# The budding yeast heterochromatic protein Sir3 modulates genome-wide gene expression through transient direct contacts with euchromatin

**DOI:** 10.1101/603613

**Authors:** Pritha Bhattacharjee, Alain Camasses, Hrvoje Galić, Ana Hrgovčić, Lara Demont, Linh Thuy Nguyen, Pauline Vasseur, Marta Radman-Livaja

## Abstract

The SIR complex (Silent Information Regulator) is the building block of heterochromatic structures that silence underlying genes. It is well established that the silenced state is epigenetically inherited but it is not known how the SIR complex is maintained through cell divisions in optimal or variable growth conditions. The biological function of heterochromatin located in subtelomeric regions is also unclear since heterochromatin coverage appears to be limited to a few kbps near chromosome ends and the expression of subtelomeric genes is only marginally affected in the absence of the SIR complex. We use a three pronged approach to address these questions. First, nanopore-MetID, an in vivo foot printing technique similar to DamID that uses nanopore sequencing technology, identified over a thousand new transient contacts between Sir3 and euchromatic genes that are not detectable by ChIP-seq and revealed a previously undocumented low-density mode of Sir3 binding to subtelomeric regions that extends 15kbps downstream of subtelomeric SIR nucleation sites. Second, our measurements of genome-wide Sir3 exchange rates after exit from starvation show that heterochromatin is a highly dynamic structure in optimal growth conditions. Third, “spike-in” RNA-seq time course experiments in the same conditions reveal that Sir3 modulates global mRNA levels in correlation with fluctuations in nutrient availability. We now propose that subtelomeric regions serve as Sir3 hubs from which Sir3 is brought over to distal sites down the chromosome arm where it transiently contacts euchromatic genes in its path. We hypothesize that contacts between Sir3 and actively transcribed genes facilitate the removal of stalled transcription complexes and allow for optimal genome-wide transcription, which gives wt cells a competitive advantage over *sir3Δ* cells when nutrients are limited.

## Introduction

Heterochromatin in budding yeast is a transcriptionally repressive structure located at the silent mating type loci (*HMR* and *HML*), telomeres and rDNA repeats. The essential component of this structure is the non-histone protein complex SIR (Silent Information Regulator), which consists mainly of Sir2, Sir3 and Sir4. Sir4 scaffolds the complex while Sir2 functions as an NAD-dependent H4K16 deacetylase, providing a high-affinity binding site for Sir3 which then recruits Sir4 (for review see (Grunstein and Gasser 2013)). In the classical polymerization model, SIR components are first recruited to silencer regions by a combination of silencer-binding factors (ORC –Origin Recognition Complex, Rap1 and Abf1). The SIR complex then spreads from the nucleation site (silencer) through cycles of histone H4 deacetylation and binding to nucleosomes, which continue until the SIR complex reaches boundary elements that prevent unwanted spreading to transcriptionally active regions (for review see (Gartenberg and Smith 2016)).

It is well established that the silent state of heterochromatic loci is epigenetically inherited but the molecular mechanisms responsible for the maintenance and renewal of the SIR complex from one cell generation to the next are not well understood. Over-expressed Sir3 can be incorporated into existing heterochromatin (Cheng and Gartenberg 2000), but beyond this bulk measurement, the locus-specific dynamics of the chromatin bound SIR complex within and from one cell generation to another have not yet been measured. How heterochromatic SIR complexes exchange their components during the cell cycle and how they are distributed to daughter chromatids after replication has important implications for how heterochromatic states are maintained and whether they may be inherited.

Likewise, the function of subtelomeric heterochromatin is still an open question. Classic RNA-seq experiments in SIR mutant backgrounds failed to detect significant changes in the expression of most subtelomeric genes (Ellahi et al. 2015) and the long standing hypothesis that the SIR complex prevents deleterious inter-chromosomal homologous recombination between repetitive subtelomeric regions similar to its role at the rDNA locus (Gottlieb and Esposito 1989) has not been experimentally corroborated (DuBois et al. 2002). On the contrary, the SIR complex stimulates the homologous recombination pathway during double strand break repair by promoting telomere clustering and bringing chromosome ends close to each other (Batté et al. 2017).

Another open question is how chromatin bound complexes that epigenetically determine gene expression states, like the SIR complex, respond to environmental challenges such as nutrient depletion. Indeed, under unfavorable conditions, yeast cells stop growing until depleted nutrients are restored. Growth arrest is characterized by transcriptional reprogramming, spatial reorganization of the genome and a 300 fold decrease in protein synthesis rates (McKnight et al. 2015). While the organization of the SIR complex in starved arrested cells has been described, the dynamics of the SIR complex during and following exit from growth arrest are poorly understood (Guidi et al. 2015). These questions have motivated us to probe SIR function and genomic localization in fluctuating nutrient conditions using new genome-wide approaches. We chose to focus on the Sir3 subunit because Sir3 is the limiting factor that determines the extent of SIR complex polymerization and the location of SIR complex boundaries (Renauld et al. 1993; Hecht et al. 1996; Radman-Livaja et al. 2011).

First, we sought to expand the catalogue of known genome-wide Sir3 binding sites by including transient and unstable contacts that are not detectable by ChIP. We reasoned that clues for the function of subtelomeric heterochromatin may lie beyond known subtelomeric nucleation sites. In order to detect these hitherto “invisible” Sir3 targets we developed Nanopore-MetID, an in-vivo genome-wide foot printing method that combines Sir3 fused to the Dam or the EcoG2 DNA-methyl transferases, which mark Sir3 targets by methylating nearby Adenines, with direct detection of methylation by long read nanopore sequencing.

Second, we measured genome wide Sir3 turnover rates after exit from nutrient deprivation with the goal to better understand the mechanisms of subtelomeric heterochromatin maintenance and renewal in variable growth conditions.

Third, in order to better understand how and if Sir3 dynamics at subtelomeric loci and transient Sir3 contacts with euchromatin, identified with Nanopore-MetID, influence global gene expression upon release from starvation we performed “spike-in” RNA-seq experiments.

Our three-pronged genome-wide approach provides a comprehensive picture of Sir3 activity before and after nutrient depletion and reveals a new role for Sir3 in the genome-wide control of gene expression.

## Results

### In vivo foot printing reveals transient Sir3 contacts at more than a thousand euchromatic genes

As mentioned above, we sought to investigate whether transient and unstable Sir3 contacts located beyond the known SIR loci that were mapped by ChIP-seq (Radman-Livaja et al. 2011) may provide clues for the function of subtelomeric heterochromatin. We have therefore developed a technique that can map transient and/or rare contacts between chromatin proteins and DNA called Nanopore-MetID for Nanopore sequencing and Methyl-adenine IDentification. This method combines nanopore sequencing (Müller et al. 2019) and in vivo methyl-Adenine foot printing similar to DamID (van Steensel and Henikoff 2000). Nanopore sequencing detects different nucleic acid bases by monitoring changes to an electrical current as nucleic acids are passed through a protein nanopore. Moreover, nanopore sequence detection can distinguish modified bases from the canonical A, T, G and C, thus making it possible to monitor DNA methylation of Cytosines or Adenine residues (McIntyre et al. 2019; Müller et al. 2019).

We fused Sir3 to the Dam or the EcoG2 DNA methyl transferases from *E.coli* (**Supplementary Figures S1 and S2)**. While Dam only methylates Adenines in GATC motifs, EcoG2 methylates all accessible Adenines. ^me^A at sites of contact between Sir3Dam or Sir3EcoG2 and DNA can then be directly read with nanopore sequencing. Unlike ChIP-seq, which detects stable long-lived Sir3-DNA/chromatin interactions, Dam and EcoG2 methylation leave a long-lived trace of short lived and/or rare contacts between Sir3 and chromatin whenever the residency time of Sir3 on chromatin is long enough to allow for Dam or EcoG2 methylation of nearby GATCs or As, respectively. Also, since nanopore sequencing does not require amplification of the isolated DNA and since we are using haploid cells, the fraction of methylated reads in the entire population of sequenced reads reflects the frequency of Sir3 contacts per cell. Given sufficient sequencing depth, this approach allows for the detection of rare contacts that occur only in a small fraction of the cell population and which would be too close to or even below the genome average read count in ChIP-seq datasets to be considered as bona fide Sir3 targets.

We first mapped ^me^A in saturated mid-log cultures of Sir3Dam cells, to minimize the number of cells undergoing replication since replication bubbles interfere with sequencing through nanopores. In addition to a wt Dam construct we also used a Dam mutant with a K9A substitution that reduces its binding affinity for GATC motifs and consequently improves Sir3 specificity of Sir3DamK9A methylation (Szczesnik et al. 2019). The ^me^A signal was determined as described in **Figure S1B** and in the “Nanopore sequencing data analysis” section of Materials and Methods in the SI Appendix. Since our yeast strains are haploid and isolated DNA fragments are sequenced directly without amplification, the ^me^A count in Sir3Dam|Sir3DamK9A cells normalized to read count and GATC content and subtracted from the normalized ^me^A count of the standalone Dam control NLSDamK9A gives us the probability that a GATC motif within a given genomic region will be specifically methylated by Sir3Dam in any cell in the population (**Figure S1C-D**). Note that the NLS sequence is crucial for the proper functioning of the standalone Dam control as there was almost no GATC methylation in a Dam standalone strain without the NLS sequence (**Figure S1E**).

The obtained GATC methylation probabilities may be an underestimate of the actual number of transient Sir3 contacts as it can only detect Sir3 contacts close to GATCs that are accessible to methylation and are for example not masked by nucleosomes and whose residency time on DNA is long enough to allow for Dam methylation. Nevertheless, our analysis reveals that at least 15% of gene promoters and at least 7% of gene coding sequences (CDS) have a non-zero probability of GATC methylation by Sir3Dam in the cell population and are consequently transiently contacted by Sir3 (**Figure S1F**). Considering that only 3% of yeast genes are located within 30kbps of chromosome ends and subtelomeric heterochromatin, the vast majority of genes that are transiently contacted by Sir3 are actually found in euchromatin.

Note that since there is on average only one GATC per 400bp in each gene promoter or CDS, and Sir3Dam foot printing does not discriminate between stable Sir3 binding within the SIR complex or transient/unstable Sir3 binding to euchromatin that occurs frequently within the cell population, the probability of GATC methylation by Sir3Dam does not reflect the density or the stability of Sir3 molecules that were bound to the methylated site. Consequently, the ^me^A signal at some euchromatic genes may be as high as or even higher than at genes located in heterochromatic regions (**Figures 1E-J and S1D**).

**Figure 1:**
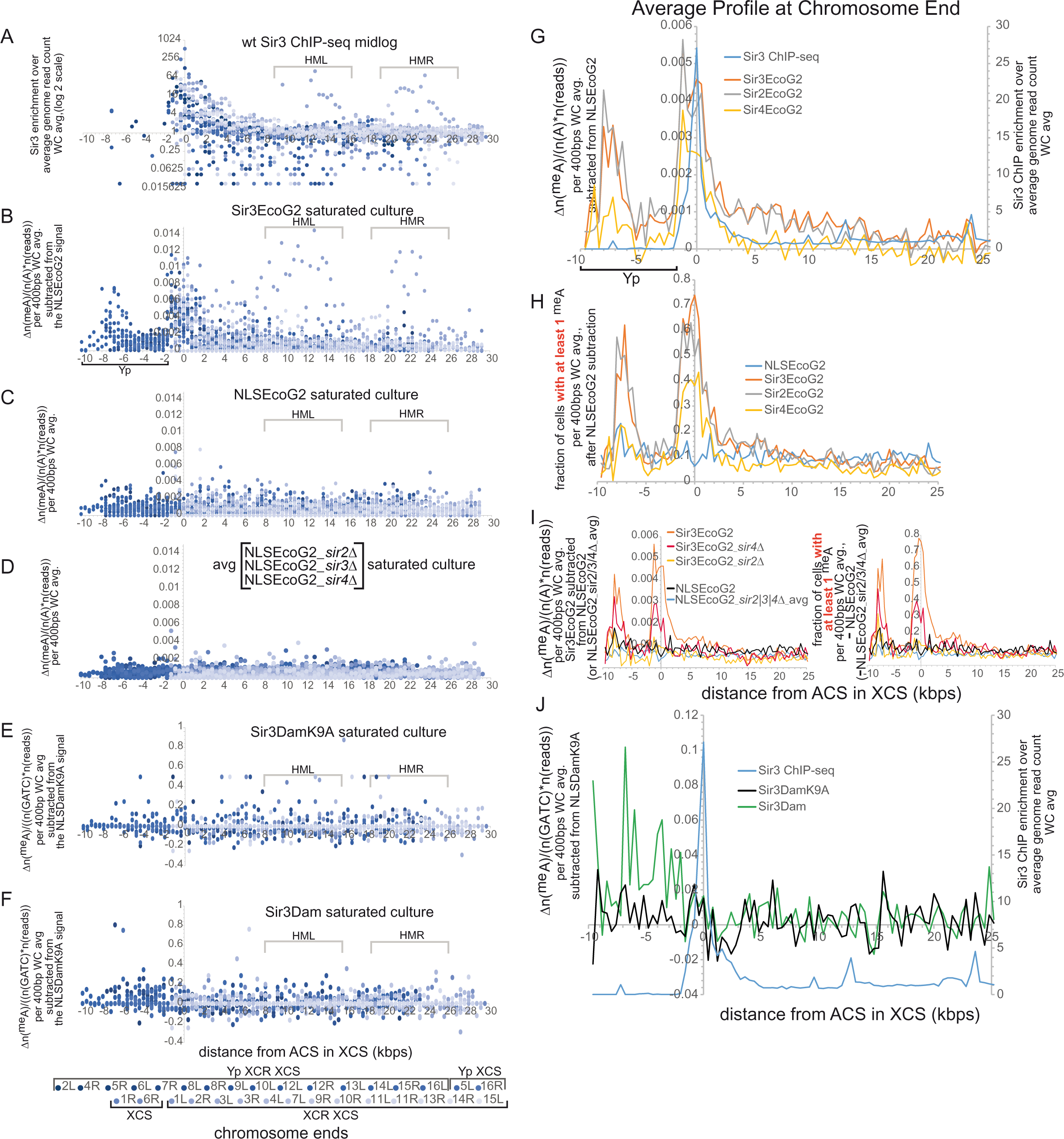
Sir3Dam and Sir3EcoG2 foot printing reveal two modes of Sir3 binding to subtelomeric regions. **A-F.** Scatter plot of Sir3 contacts 30-40kbps from all 32 chromosome ends measured by Sir3 ChIP-seq (A, from Radman-Livaja et al. (2011)), Sir3EcoG2 foot printing (B), Sir3DamK9A foot printing (E) and Sir3Dam foot printing (F). Stand-alone NLSEcoG2 in a wt genetic background and stand-alone NLSEcoG2 in *sir2Δ, sir3Δ* or *sir4Δ* cells controls are shown in C and D, respectively. Chromosome ends are aligned by the location of the ACS (ARS Consensus Sequence) motif within the XCS subtelomeric sequence. Points from different chromosome ends are drawn in different shades of blue (see legend below the graph in F). **G.** Average ^me^A density for Sir3EcoG2, Sir2EcoG2 and Sir4EcoG2 (y axis on the left) and Sir3 ChIP signals (y axis on the right) at chromosome ends. The peak at −8kbps from the ACS in the Sir3EcoG2 profile comes from the Yp region in chromosome ends that contain it (see B). Note that Yp regions are highly repetitive and cannot be mapped by ChIP-seq. **H.** Average fraction of cells in the population with at least 1 ^me^A per 400bp segment (see **Nanopore sequencing data analysis** in the SI Appendix for calculations) for Sir3EcoG2, Sir2EcoG2 and Sir4EcoG2 at chromosome ends. **I.** Average ^me^A density (left) and average fraction of cells in the population with at least 1 ^me^A per 400bp segment (right) at chromosome ends for Sir3EcoG2 in wt, *sir2Δ* and *sir4Δ* genetic backgrounds, and the standalone NLSEcoG2 control and the average of the NLSEcoG2 controls in *sir2Δ, sir3Δ* and *sir4Δ* genetic backgrounds. The Sir3EcoG2 wt signal was subtracted from the standalone NLSEcoG2 control and the Sir3EcoG2 signal from *sir2Δ, sir4Δ* and *sir2Δsir4Δ* cells was subtracted from the average NLSEcoG2 signal from *sir2Δ, sir4Δ* or *sir3Δ cells.* **J** Average GATC Adenine methylation probability per 400bp segment at chromosome ends for Sir3Dam and Sir3DamK9A (y axis on the left) and Sir3 ChIP signals (y axis on the right).

We consequently used a Sir3EcoG2 fusion to assess Sir3 density and/or stability at the binding site, both of which are directly correlated to the density of ^me^A in a given region. We also determined the frequency of specific Sir3 contacts in the cell population, which is calculated from the number of reads (equal to the number of cells for our haploid strains) that contain at least one ^me^A in a given genomic region (**Supplementary Figure S2A-C**). The Sir3EcoG2 signal has been subtracted from the standalone NLSEcoG2 control to control for EcoG2 methylation that is not specifically directed by Sir3. Since Adenines comprise ∼25% of the yeast genome, the density of ^me^A determined by the normalization of the ^me^A count by the read count and Adenine content in 400bp windows, yields a Sir3EcoG2 methylation profile that is remarkably similar to the Sir3 ChIP-seq profile (**Figures 1A-B and S2B-D**).

However, not more than 1.5% of Adenines are methylated specifically by Sir3EcoG2 (HML and HMR) even in regions that are stably bound by the SIR complex (**Figure 1B**). The ^me^A density around the XCS nucleation site is on average only 0.45%, probably because most Adenines are masked by nucleosomes and inaccessible to EcoG2 (**Figure 1G**). Thanks to long read nanopore sequencing, Nanopore–MetID with Sir3EcoG2 also revealed a new SIR nucleation site in the repetitive Yp region located 8kbps upstream of the well documented nucleation site in the XCS region (**Figure 1B, G**). The Yp nucleation site is similar in terms of Sir3 binding to the XCS site with an average ^me^A density of ∼0.4% and SIR spreading of ∼2 kbps around the highest ^me^A density peak (**Figure 1B**). Also, the Yp and XCS nucleation sites seem to be independent of each other as the ^me^A density in the XCS site is not significantly different between chromosome ends that have the Yp site and those that do not (**Figure 1B**).

The Yp and XCS nucleation sites are methylated on average in 72% of cells (**Figure 1H**) and the fraction of cells with at least one ^me^A/400bp drops down to 10%, 30kbps downstream of the XCS nucleation site. Likewise, ^me^A density decreases from 0.45% at the XCS sites down to 0.1%, 4 to 15 kbps downstream of the XCS and then down to 0.02% beyond that. Note that the Sir3 ChIP-seq signal reaches background levels by 4kbps downstream of the XCS site, a region where ^me^A density in the Sir3EcoG2 strain is still at 0.1% above background (determined by the standalone NLSEcoG2 signal). This is consistent with our hypothesis that a single-molecule, in vivo foot printing approach is better at detecting Sir3 contact sites with low Sir3 density.

In fact, Sir3EcoG2 methylates 0.08% of Adenines (an average of ∼6000 Adenines per genome) on the Watson or the Crick strand in 99% of the genome (**Figure S2E**). Sir3EcoG2 also covers more genomic sites than NLSEcoG2 as only 1% of the genome is not methylated in any Sir3EcoG2 cell while 15% of the genome is not methylated in any NLSEcoG2 cell. Most of the genomic sites are however only methylated in a subset of cells in both strains: 97% of the genome is methylated on average in 13% or 7% of Sir3EcoG2 or NLSEcoG2 cells, respectively (**Figure S2E bottom left**). Additionally, Sir3EcoG2 methylates 50% of the genome more efficiently than NLSEcoG2 (**Figure S2E bottom right).** It is important to mention here that this does not mean that the sites that are more efficiently methylated by NLSEcoG2 are actually not contacted by Sir3. It rather means that these potential Sir3 contacts are more transient and less frequent than NLSEcoG2 contacts, which makes it impossible to know if the methylation detected at these sites in Sir3EcoG2 cells is actually dependent on Sir3. We also suspect that Sir3 almost certainly makes contact with many more genomics sites in each cell, but most of those contacts are either too short lived or the underlying Adenines are inaccessible for methylation by EcoG2.

Interestingly, Adenines in the Crick strand are more methylated than As in the Watson strand perhaps reflecting differences in nucleotide accessibility between the two DNA strands wrapped around nucleosomes. The preferential methylation of the Crick strand relative to the Watson strand is specific to Sir3EcoG2 as there are no differences between methylation profiles of Watson and Crick strands in the standalone NLSEcoG2 strain (**Figure S2E top).** In the absence of Sir2 or Sir4, the Sir3EcoG2 dependent methylation of the two strands resembles more the Watson and Crick methylation profiles in the NLSEcoG2 strain (**Figure S2E**), suggesting that the genome-wide contacts observed between Sir3EcoG2 and euchromatin in the wt strain depend on the formation of the SIR complex at chromosome ends.

Promoters are more methylated by Sir3EcoG2 than CDSes, with 35% of promoters versus 26% of CDS that have at least one ^me^A in at least 5% of cells, respectively (**Figure S2F**). This probably reflects the higher chromatin accessibility of promoters, which are largely depleted of nucleosomes.

In summary, our Sir3EcoG2 mediated Adenine methylation profiles of chromosome ends show that Sir3 has two modes of binding to chromatin: a high and a low density binding mode. Sir3 binds at high density ±2kbps around SIR nucleation sites in ∼70% of cells. Beyond these regions and down to 30kbps downstream, Sir3 still contacts DNA in 10% to 20% of cells but its density drops 5 fold to levels undetectable by population based ChIP-seq. The two modes of Sir3 association with chromatin can be detected but cannot be differentiated by Sir3Dam foot printing. These three approaches produce complementary maps of genome wide Sir3 association with chromatin. Sir3 ChIP-seq establishes a map of high-density stable Sir3 binding to non-repetitive regions. Sir3Dam Nanopore-MetID produces probabilities of Sir3 contacts with genomic targets, including repetitive regions, but does not discriminate between targets to which Sir3 binds at high density and targets to which Sir3 binds with low density and high frequency. Finally Sir3EcoG2 Nanopore-MetID provides Sir3 contact maps that delineate regions of high-density Sir3 occupancy from sites with transient low density Sir3 contacts.

This transient low-density chromatin binding mode is not specific to Sir3 as ^me^A is detected at similar levels in the Sir2EcoG2 strain and at a twofold lower level in the Sir4EcoG2 strain (**Figure 1G-H**). Adenine methylation by Sir3EcoG2 also depends on the presence of Sir2 and to a lesser extent of Sir4. The *sir2*Δ deletion almost completely abolishes Sir3EcoG2 methylation in subtelomeric regions while the *sir4*Δ deletion causes a 2 fold reduction in ^me^A levels in the same region (**Figure 1I**).

Note that the ^me^A signal in Sir3EcoG2 and Sir3DamK9A strains shown in Figures 1 and 2 is specific for Sir3 and is not a product of spurious EcoG2 or DamK9A methylation that is independent of Sir3, because the GATC methylation probability and both the ^me^A density and the fraction of methylated cells for each 400bp genome segment were derived from ^me^A/read count ratios from the Sir3DamK9A or Sir3EcoG2 strains that were subtracted from ^me^A/read count ratios from the standalone NLSDamK9A or NLSEcoG2 strains, respectively (**Figures S1C and S2C**).

**Figure 2:**
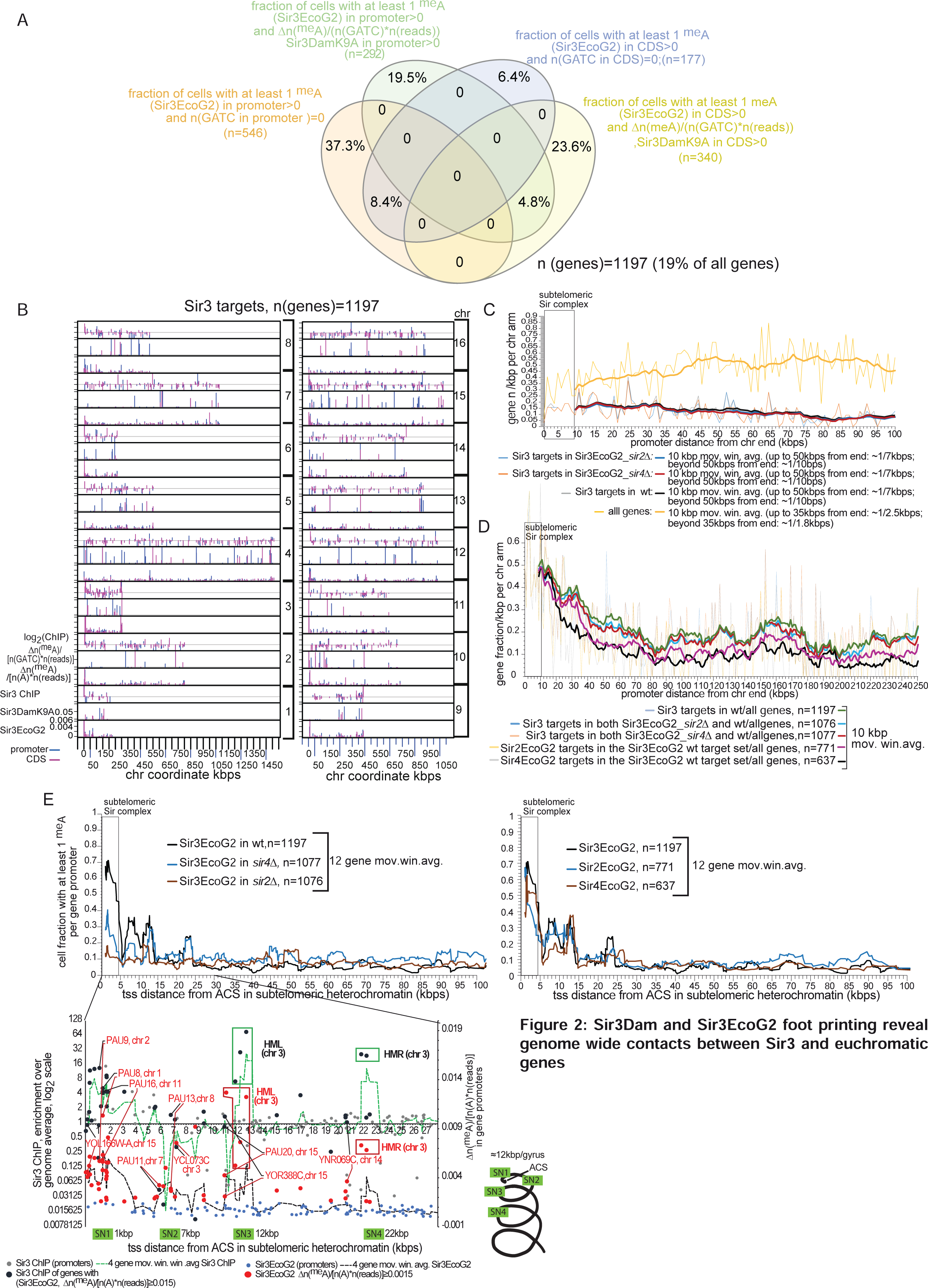
Sir3Dam and Sir3EcoG2 foot printing reveal new genome wide contacts between Sir3 and euchromatic genes. **A.** There are 632 genes that have a non-zero probability of GATC methylation by Sir3DamK9A methylation: Δ[n(meA)/(n(GATC)*n(reads))] = [[n(meA,Sir3DamK9A)/(n(GATC)*n(reads))]-[n(meA,NLSDamK9A)/(n(GATC)*n(reads))]] > 0, and a non-zero probability of having at least 1 ^me^A in the Sir3EcoG2 strain, either in their promoter or CDS or both. Additionally there are 723 genes with a non-zero probability of having at least 1 ^me^A in the Sir3EcoG2 strain that do not have GATC motifs in their promoter, CDS or both that were included in the set of 1197 genes that are contacted by Sir3. **B.** Locations of the 1197 genes from A on each chromosome. **C.** Average number of gene promoters per kbp per chromosome arm 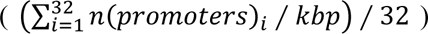 versus distance from the chromosome end (kbps) for all genes and Sir3 targets (from A) in the wt, *sir2Δ* and *sir4Δ* genetic backgrounds. Sir3 targets in *sir2Δ* and *sir4Δ* are a subset of sir3 targets in wt. **D.** Fraction of Sir3, Sir2 and Sir4 targets (as defined in A) per kbp per chromosome arm as a function of distance from chromosome ends. The fractions of Sir3 targets in a wt genetic background are calculated by dividing the gene density of Sir3 targets from C by the gene density of all genes from C, while Sir3 targets in *sir2Δ* and *sir4Δ* backgrounds, and Sir2 and Sir4 targets are are a subset of Sir3 targets in the wt background, also divided by the gene density of all genes from C. **E.** Meta-profiles of cell fractions with at least 1^me^A per gene promoter for Sir2EcoG2 and Sir4EcoG2 targets among Sir3EcoG2 wt targets (right top panel), or for Sir3EcoG2 targets in wt, *sir4Δ* and *sir2Δ* cells (left top panel) aligned by the ACS in the XCS of each chromosome arm and averaged over all 32 chromosome arms. The bottom panel compares median cell fractions with at least 1^me^A per gene promoter for Sir3EcoG2 (y axis on the right), and Sir3 ChIP-seq enrichments (y axis on the left) of genes located at the Sir3EcoG2 peaks located 1kbp (SN1), ∼7kbps (SN2), ∼13kbps (SN3) and ∼22kbps (SN4) downstream of the ACS. A schematic of a chromosome end wrapped into a spiral with 12kbps/gyrus and the locations of SN1-4 is shown on the right of the plot.

In order to identify genes that are contacted by Sir3, we determined the median GATC methylation probability, and the ^me^A density and the fraction of methylated cells for each gene promoter and CDS in the Sir3DamK9A and Sir3EcoG2 strains, respectively. We used these measurements to identify euchromatic genes that are contacted by Sir3 in at least part of the cell population (**Figure 2**). We considered genes as bona fide Sir3 contacts when their promoter or CDS has a non-zero probability of having at least 1 ^me^A in the Sir3EcoG2 strain and a non-zero probability of GATC methylation (if they contained GATCs) in the Sir3DamK9A strain (**Figure 2A**). We identified 1197 genes (19% of all genes) that fulfilled these criteria (**Table S1**).

The 1197 genes that are contacted by Sir3 are for the most part evenly distributed throughout each yeast chromosome (**Figure 2B**). **Figure 2C** compares the density distribution of all yeast genes (i.e. the average number of gene promoters per kbp per chromosome arm) with the density distribution of Sir3 targets in wt, *sir2Δ* and *sir4Δ* strains, as a function of their distance from the end of the chromosome. When considering the entire set of genes, we see that gene density increases 30% in the regions beyond 50kbps from the chromosome end. Conversely, Sir3 targets are more densely packed close to chromosomes ends: Sir3 contacts 50% of genes within 20kbps of the chromosome end. The density of Sir3 targets than gradually drops from 50% to 20% of all genes in the region between 20kbps to 50kbps from the end of the chromosome and oscillates around 20% beyond that limit (**Figure 2C-D**). While the number of Sir3 targets is only reduced by 10% in the absence of Sir2 and Sir4 (**Figure 2C-D**), their absence does affect the distribution profiles of Adenine methylation frequency in the cell population along the chromosome arm. *sir2Δ* and *sir4Δ* strains show reduced methylation frequency in the cell population compared to wt in the first 25kbps and 15kbps downstream of the XCS, respectively. Both mutants also have a somewhat higher methylation frequency than wt beyond 25kbps. Also, 64% or 53% of Sir3 targets beyond the subtelomeric SIR complex are contacted by Sir2 or Sir4, respectively (**Figure 2D**). These two observations taken together with the fact that Sir3EcoG2 differentially methylates the Watson and Crick strands only in the presence of Sir2 and Sir4 (**Figure S2E, top**), are consistent with the idea that the subtelomeric SIR complex acts as a Sir3 reservoir from which Sir3 is brought over to distal euchromatic regions through telomere looping.

Remarkably, the distribution of ^me^A density in Sir3EcoG2 cells also provides clues about the 3D structure of chromosome ends (**Figure 2E**). If we average profiles of methylation frequencies in the cell population originating from the ACS in the XCS region of all chromosome arms, we detect four discrete peaks spaced 6-7kbps from each other starting ∼1kbp downstream from the ACS (SN1 to SN4 respectively, for Secondary Nucleation site). While ^me^A densities at HML and HMR located on the left and right arm of chr 3, ∼13 and ∼22 kbps downstream from the XCS, respectively, contribute significantly to the average ^me^A profiles at SN3 and SN4, we also find genes with comparably high ^me^A densities that are located in the same regions but on other chromosomes (**Figure 2E**, bottom panel). The SIR complex is however probably not assembled in the SN3-4 regions that don’t contain HML or HMR when Sir3 is expressed at wt levels, since, contrary to HMR and HML, these genes don’t show any significant Sir3 enrichment measured with Sir3 ChIP (Figure 2E, (Radman-Livaja et al. 2011)). Likewise, the SN2 region at 7kbp downstream from the XCS contains PAU genes that show high ^me^A density in the Sir3EcoG2 strain but no significant Sir3 enrichment in the wt Sir3 ChIP-seq dataset. Peaks of Sir3 enrichment at PAU genes located in the SN2 region only appear in ChIP-seq datasets if Sir3 is over expressed (Figure 6B and (Radman-Livaja et al. 2011)), thus demonstrating the high sensitivity and specificity of Nanopore-MetID. Indeed, the only Sir3 targets that are detectable by both ChIP and Nanopore-MetID when Sir3 is not over-expressed are located in the SN1 region that is immediately adjacent to SIR nucleation sites in the XCS region.

The non-random distribution of SN1-4 is consistent with the folding of chromosome arms into a spiral with ∼12kbp/gyrus that brings SN1-4 closer to the XCS, as illustrated in the bottom panel of Figure 2E. The 12kbps/gyrus measurement estimated from Nanopore-MetID data is remarkably close to the 13.2 kbp/gyrus calculations for the spiral fold of human chromosome 10 (Sedat et al. 2022), suggesting that yeast cells are “taking advantage” of the natural propensity of chromatin fibers to fold into a spiral coil, in order to facilitate the propagation of Sir3 from the chromosome end towards the middle of the chromosome.

### Old Sir3 is rapidly degraded upon release from starvation

Sir3EcoG2 and Sir3Dam foot printing revealed that, in exponentially growing cells, Sir3 makes transient low-level contacts along the entire length of each yeast chromosome. The average Sir3Dam dependent GATC methylation of Sir3 targets identified in Figure 2, drops down to zero during carbon starvation and does not recover even 90 minutes after release (**Figure 3A**). Since nutrient deprivation appears to drastically decrease Sir3 binding to the entire genome, we decided to measure cellular dynamics of Sir3 in these conditions.

**Figure 3:**
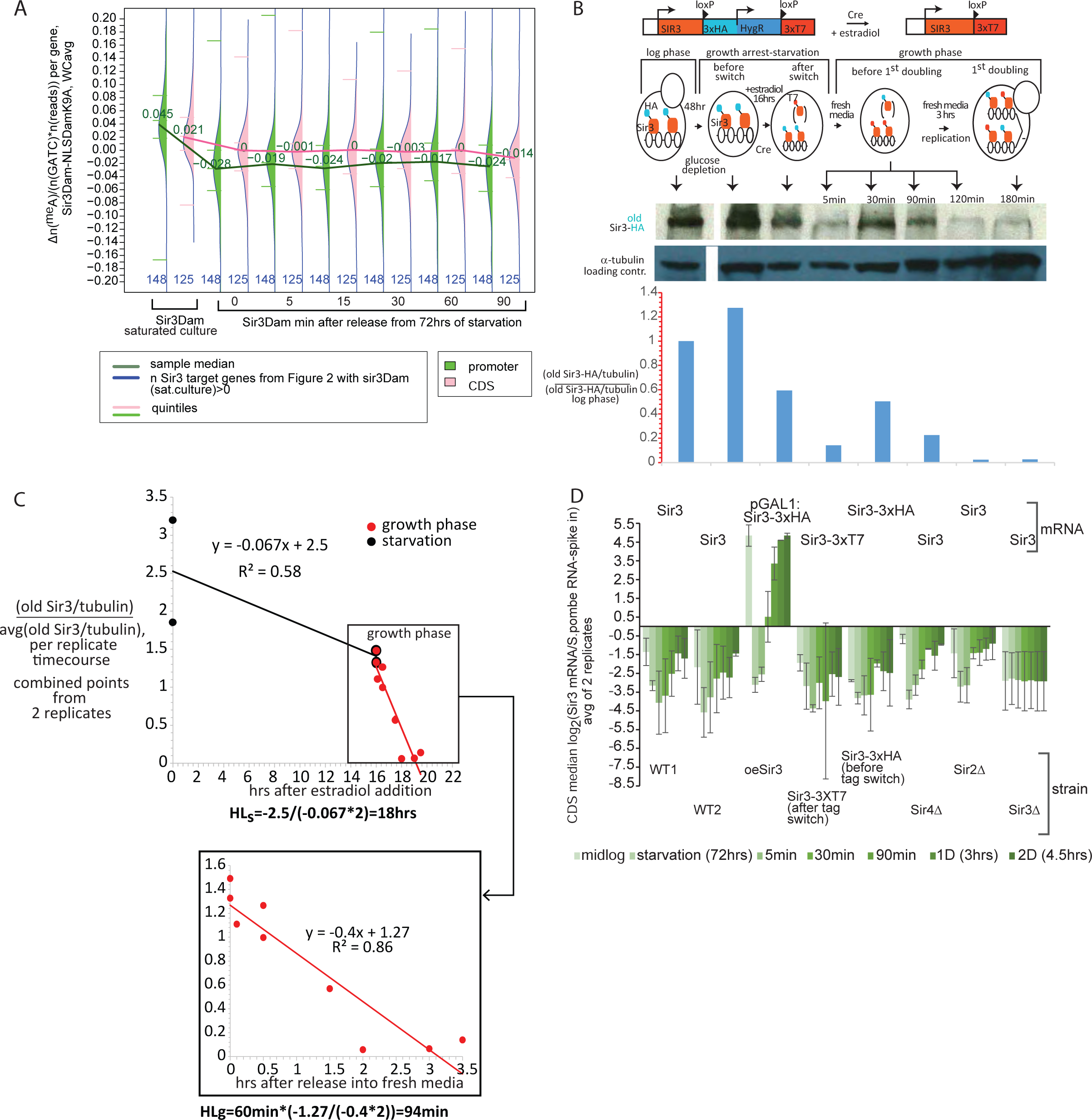
Sir3 degradation rates increase 6 fold after release from starvation A. Bean plot distribution of the probability of GATC methylation by Sir3Dam after release from starvation. We used gene subsets from the 3393 set of Sir3 targets identified in Figure 2 that had a non-zero probability of GATC methylation by Sir3Dam in saturated mid-log cultures and plotted the probability of GATC methylation by Sir3Dam of those genes during starvation and at indicated times after release from starvation. **B.** Western blot of “old” Sir3-3xHA before and after tag-switch from the Sir3-3xHA to Sir3-3xT7 RITE tag switch strain during and after release from growth arrest. The top panel shows the experimental outline and describes the time points when total cell extracts were isolated (marked by arrows above the blot). The bar graph below the blot shows the quantification of the bands from the blot. Sir3 band intensities were first normalized to the α-tubulin loading control and then divided by the normalized Sir3-HA intensity from mid-log cells. One of two biological replicates is shown. The α-tubulin and Sir3 bands were on the same gel but we had to use two different exposures for detection and quantification: weak Sir3-3xHA signals required a long exposure that saturated the strong α-tubulin signal for which we had to use a much shorter exposure. **C.** Sir3 half-life in growth arrest and after exit from growth arrest. The top plot shows the decrease in Sir3HA band intensity over time with combined points from western blots of two biological replicates. The half-life of Sir3 during starvation is calculated from the decrease in Sir3-HA band intensity after the induction of the tag switch with estradiol addition to starved cells, according to the equation: HLs=-b/(2*a), where b and a are the y cutoff and slope of the linear fit equation (black line). The half-life of Sir3 after release (red line) HLg is calculated with the same equation using the y-cutoff and slope from the linear fit in the bottom plot. **D.** Sir3 gene expression. Median Sir3 read density enrichment from the RNA-seq datasets normalized to an external spike-in *S. pombe* RNA control (from datasets in Figure 7), normalized as described in Figure S7. The error bars represent the standard error between two biological replicates.

We first wanted to assess bulk Sir3 degradation and synthesis rates during and after release from starvation. We used the RITE system (Verzijlbergen et al. 2010) to construct the Sir3 tag switch module. The genomic *SIR3* locus was engineered with a C-terminal tag switch cassette containing loxP sites, and the 3xHA and 3xT7 tags separated by a transcription termination signal and the hygromycin resistance gene (**Figure 3B**). The host strain carries the CreEBD78 recombinase, which is activated upon estradiol addition. After Cre-mediated recombination of LoxP sites, the 3xHA tag on Sir3 is switched to the 3xT7 tag.

A saturated over-night culture was diluted 10 fold in glucose rich media (YPD) and cells were incubated for 48 hrs until they stopped dividing. They were then kept in growth arrest for ∼16 hrs after estradiol addition to allow for the tag switch to complete. Whole cell extracts were taken at indicated times during the time course and the amount of old HA tagged Sir3 was quantified by Western blot (**Figure 3B**). Note that in these conditions, only ∼50% of cells have reached a quiescent state(Allen et al. 2006), which is characterized by the reorganization of subtelomeric SIR complexes into superfoci(Guidi et al. 2015). Consequently, the population in these conditions is a mixture of “pre-quiescent” cells with several subtelomeric SIR foci and quiescent cells, with one subtelomeric SIR “super focus”.

The apparent half-life of Sir3 during growth arrest was estimated to be ∼18 hrs (**Figure 3C).** Steady state Sir3 amounts in starved cells are maintained at a similar level as in mid-log cells (**Figure 3B**), probably due to residual Sir3 gene expression **(Figure 3D**) and slow protein synthesis that is sufficient to compensate for slow Sir3 protein degradation during early starvation.

Sir3 decay rates increase 6 fold immediately after release from starvation and the half-life of old Sir3 drops down to ∼94 min from 18hrs during starvation (**Figure 3C**). Considering that most yeast proteins have a half-life between 60 and 150 min in exponentially growing cells(Auboiron et al. 2021), we suspect that old Sir3 is degraded by the usual cellular protein degradation machinery, which probably resumes its activity at pre-starvation rates shortly after release.

The genome-wide loss of Sir3Dam mediated GATC methylation observed in the first 90min after release from starvation (**Figure 3A**) is consistent with the sudden increase in Sir3 degradation rates immediately after release from starvation and the slow recovery of Sir3 gene expression that only reaches mid-log levels after the first division after release (**Figure 3D**). We now hypothesize that the rapid degradation of “old” Sir3 after release from starvation, compounded with the delay in the reactivation of Sir3 gene expression, could transiently destabilize the SIR complex and impair its silencing function during the first cycle after release.

### The SIR complex is transiently destabilized upon exit from starvation

Since the dramatic decrease in old Sir3 levels immediately after release from starvation that is accompanied by slow synthesis of new Sir3 (**Figure 3**) could compromise heterochromatin formation, we performed the “α-factor assay” to directly test the silencing function of SIR complexes after release from starvation (**Figure 4**).

**Figure 4:**
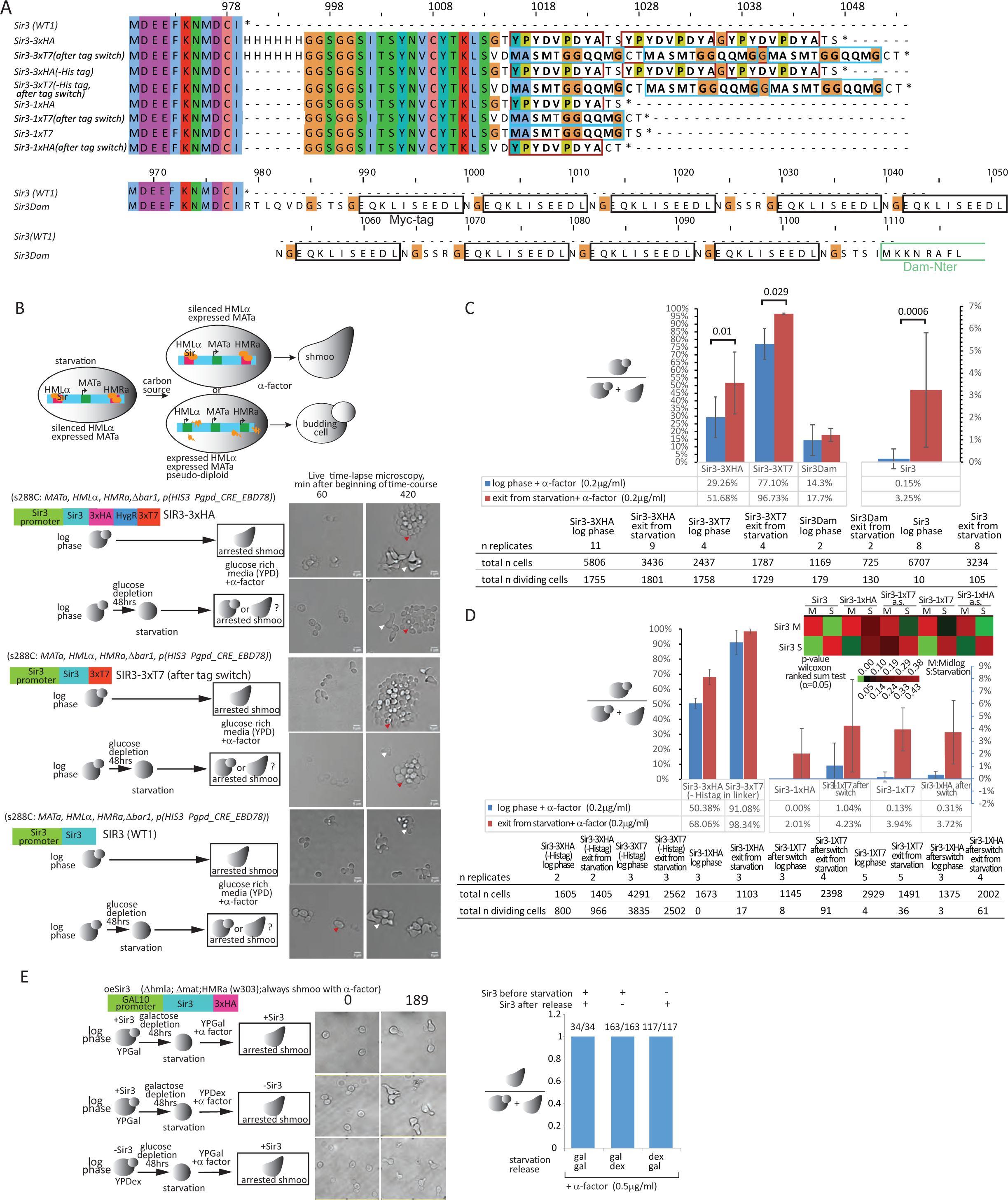
The SIR complex is destabilized upon exit from starvation. **A.** Amino-acid sequence alignment of the C-terminus of Sir3 with different epitope tags used in this study. The Sir3 strain (WT1) is the parent strain of all the RITE Sir3 tagged strains below. The Sir3-3xHA strain is the RITE strain used throughout the study. The tag switch was not induced in this experiment. All the strains marked with “after tag switch” were selected from their respective RITE strains (the strain listed in the line above the “after the tag switch strain”) after the tag switch was completed. The Sir3-3xHA (-His tag) is another RITE strain with a Sir3-3xHA to 3xT7 switch that does not have a 6xHis linker between the C-terminus and the first LoxP site. The Sir3Dam strain is the strain used in Figures 1 and S1. The linker between the Dam sequence and the C-terminus of Sir3 has 9 Myc tags. **B**. α-factor heterochromatin stability test. The diagram on top shows the expected response of MATa cells to α-factor added after release from starvation. If the SIR heterochromatic complex is unstable HMLα and HMRa will be transcribed along with MATa, thus creating pseudo-diploid cells that don’t respond to α-factor and consequently do not become shmoo but start budding instead. The images show examples of frames from a live cell imaging experiment that follows Sir3-3xHA, Sir3-3xT7 and Sir3 (WT1) log phase cells or starved cells in the first cell cycle after release in the presence of 0.2 µg/ml α-factor 60 and 420 min after the beginning of the time course. Red arrows: budding cells; White arrows: shmoos. **C.** The bar graph shows the fraction of budding cells out of the total number of cells (budding and shmoo) for each strain and growth condition from B. and the Sir3Dam strain. The error bars represent the standard deviation from the mean of all the replicates summarized in the table below the graph. P-values for the Wilcoxon ranked sum test (α=0.05) are listed above the corresponding bars in the graph. **D.** same as C for the remaining set of strains from A that were not shown in B-C. **E.** α-factor test with the oeSir3 strain which shmoos independently of Sir3 in the presence of α-factor because of the deletion of HML and MAT loci.

*MATa* (Mating Type a) cells respond to α-factor (mating pheromone α) by arresting in G1 and forming a mating projection termed shmoo. When the *HML* locus is not fully silenced, cells behave as pseudo-diploids and do not respond to α-factor. These cells therefore keep on budding in the presence of pheromone. Exponentially growing cells predominantly respond to α-factor and shmoo. Our hypothesis on the transient instability of the SIR complex during the starvation to growth transition predicts that populations exposed to α-factor will have a higher fraction of cells that do not shmoo upon exit from starvation. We observed that 52% of cells from the RITE strain used in Figure 3 do not shmoo upon release from growth arrest, which is nearly twice the rate of budding cells in mid-log populations (29%) exposed to α-factor (**Figure 4**).

However, the high proportion of exponentially growing cells that were unresponsive to α-factor was surprising. Suspecting an adverse effect of the 3xHA epitope tag on mRNA stability or the stability or function of the protein, we performed the same experiment using strains with an untagged Sir3 (WT1) and the Sir3 RITE strain isolated after the tag switch (Sir3-3xT7) (**Figure 4A-C**). Insensitivity to α-factor does increase 20-fold after exit from growth arrest in cells with the untagged Sir3 compared to exponentially growing untagged cells but the overall effect is greatly diminished compared to tagged Sir3 cells since only 3.25% and 0.15% of the cell population released from arrest and mid-log cells, respectively, did not shmoo in the presence of α-factor (**Figure 4C**).

The degree of insensitivity to α-factor apparently depends on the size and type of the epitope tag and the linker sequence that connects the tag to the C-terminus of Sir3 (**Figure 4A**). While a triple HA tag substantially reduced HML silencing, the triple T7 tag almost completely obliterated it with 77% and 97% of mid-log and starved cells, respectively, insensitive to α-factor. Additionally, the absence of a 6xHis tag in the linker region upstream of the 3xHA and 3xT7 tags impaired Sir3 function even more, while Sir3 with a single HA or T7 tag were both functionally indistinguishable from untagged Sir3 (**Figure 4D**). The C-terminal Dam fusion in the Sir3Dam strain (**Figure 1**) has a mild hypo-morph effect, with a 2 fold and 5 fold better response to α-factor than the two RITE strains, respectively (**Figure 4C**). Note that, our single cell α-factor assay for SIR complex function is more stringent than the more commonly used population based tests that measure the level of repression at silent mating type loci using RT-qPCR. RT-qPCR is not sensitive enough to detect increased mRNA expression in a small subpopulation of cells like our Sir3Dam strain in which the vast majority of cells (i.e. ∼85%) are fully functional. Consequently, Sir3 tagged strains that appear fully functional with an RT-qPCR based test like the Sir3EcoG2 strain described in a recent article(Brothers and Rine 2022) may actually have mild hypo-morph phenotypes that only become apparent when we use a quantitative single cell based test. Furthermore, considering that the Sir3EcoG2 construct has the same Myc tag linker between the C-terminus of Sir3 and EcoG2 as the Sir3Dam construct, which is fully functional in 85% of mid-log cells, and that the Sir3EcoG2 methylation pattern at canonical SIR loci mirrors wt Sir3 enrichment measured by ChIP-seq (**Figure 1**), we conclude that, at the cell population level, Sir3EcoG2 is functionally equivalent to the untagged wt Sir3.

We also confirmed that the increased inability of cells to shmoo upon release from growth arrest was directly linked to the de-repression of the *HML* locus caused by Sir3 instability and not by an indirect effect linked to release from starvation. Cells that shmoo in log phase in the presence of α-factor irrespective of the presence of Sir3 due to a *HMLα/MATa* double deletion, also shmoo with 100% efficiency upon release from growth arrest even in the absence of Sir3 (**Figure 4E**).

Cells resumed budding upon refeeding in the absence of α-factor in the same experimental set up as the α-factor test described above, with an average efficiency of 98% ± 0.9% for all strains tested in Figure 4. This indicates that the triple tags that impair the silencing function of Sir3 at silent mating type loci do not otherwise affect the cell’s ability to resume growth after starvation.

Next, we wanted to test if fluctuations in cellular Sir3 levels (**Figure 3**) and the weakening of the SIR silencing function (**Figure 4**) after release from starvation correlate with changes in Sir3 levels at silent mating type loci. We consequently used the Sir3-3xHA to 3xT7 and the Sir3 1xT7 to 1xHA RITE strains to measure Sir3 turnover rates at silent mating type loci (**Figure S3**). Our results show that exchange rates increase dramatically after release from starvation resulting in a complete replacement of old Sir3 with the newly synthesized Sir3 by the end of the first cell cycle after release. Moreover, Sir3 OFF rates are comparable between the triple and single tag strains, while the ON rates are 1.7 fold higher in the single tagged strand at the HMR-E locus, but are not different at the HMLα locus. Since the hypo-morph phenotype of Sir3-3xHA correlates with a six fold lower Sir3 occupancy at heterochromatic loci compared to wt mid-log cells (**Figure S4**) and since the single tagged Sir3 strains do not exhibit a hypo-morph phenotype and consequently probably have a similar occupancy of heterochromatic loci as wt Sir3, we conclude that, as expected, Sir3 occupancy does not influence OFF rates. Also unsurprisingly, ON rates appear to be influenced by Sir3 levels in a locus specific manner. We therefore conclude that at the very least the OFF rates of Sir3 subunit exchange within the SIR complex are not affected by the hypo-morph phenotype of the triple tag.

The triple tag may either destabilize the mRNA or the protein or interfere with either the ability of Sir3 to interact with other SIR complex subunits or with its ability to bind to chromatin. An increase in mRNA instability is probably not the main cause of the more than 200 fold increase in insensitivity to α-factor in mid-log Sir3-3xHA|3xT7 cells compared to mid-log wt cells (**Figure 4C**), since RNA-seq datasets normalized to an external *S. pombe* RNA “spike-in” (**Figure 7**) show that, during exponential growth, *SIR3* mRNA levels are on average only 2 fold lower in the two tagged Sir3 strains than in wt strains (**Figure 3D**). The slow recovery to mid-log *SIR3* mRNA levels that takes the entire first cell cycle after exit from starvation is consistent with the increase in α-factor insensitivity relative to mid-log cells during this period in all three strains (**Figure 3D**). Sir3 enrichment at nucleation sites in subtelomeric and silent mating type loci is 6 to 8 fold lower in the 3xHA strain compared to an untagged WT strain (Radman-Livaja et al. 2011). The extent of Sir3 spreading from nucleation sites is nevertheless similar between the untagged and tagged strain (**Figure S4**). This suggests that the triple tag either destabilizes the Sir3 protein and causes an overall decrease in cellular Sir3 levels and/or impairs its ability to bind to nucleation sites but does not interfere with SIR polymerization. Consequently the hypo-morph phenotype of the triple tagged Sir3 is due to low Sir3 occupancy at subtelomeric and silent mating type loci.

Our results led us to conclude that the transient de-repression of the HML locus after release from starvation is directly linked to a decrease in Sir3 occupancy caused by more than a 1.5 to 2 fold increase in Sir3 OFF rates and 6 fold increase in Sir3 decay rates that is not immediately compensated by the recruitment of newly synthesized Sir3 to silent mating type loci.

### Sir3 exchange rates in subtelomeric regions increase after release from starvation

Next, we wanted to investigate whether the rapid increase in Sir3 exchange rates upon release from starvation observed at silent mating type loci can be generalized to subtelomeric heterochromatin. We consequently repeated the Sir3 turnover experiment from Figure S3 and processed ChIP-ed DNA for Illumina sequencing. We had to use the triple tag RITE strain for the ChIP-seq experiments as the single tag RITE strain did not yield sufficient amounts of ChIP-ed DNA to produce Illumina sequencing libraries of good quality. However, since the triple tag and the single tag strains had similar Sir3 OFF rates in our ChIP-qPCR experiments while it affected the ON rate only at HMR-E and not at HMLα, we surmised that at least the OFF rates obtained from ChIP-seq experiments with the triple tag strain should be comparable to the Sir3 OFF rates of the untagged Sir3.

ChIP-seq datasets of old and new Sir3 showed that Sir3 dynamics are similar at all heterochromatic loci, including silent mating type loci and subtelomeric regions (**Figure 5B-D and replicate experiment in Supplementary Figure S5**).

**Figure 5:**
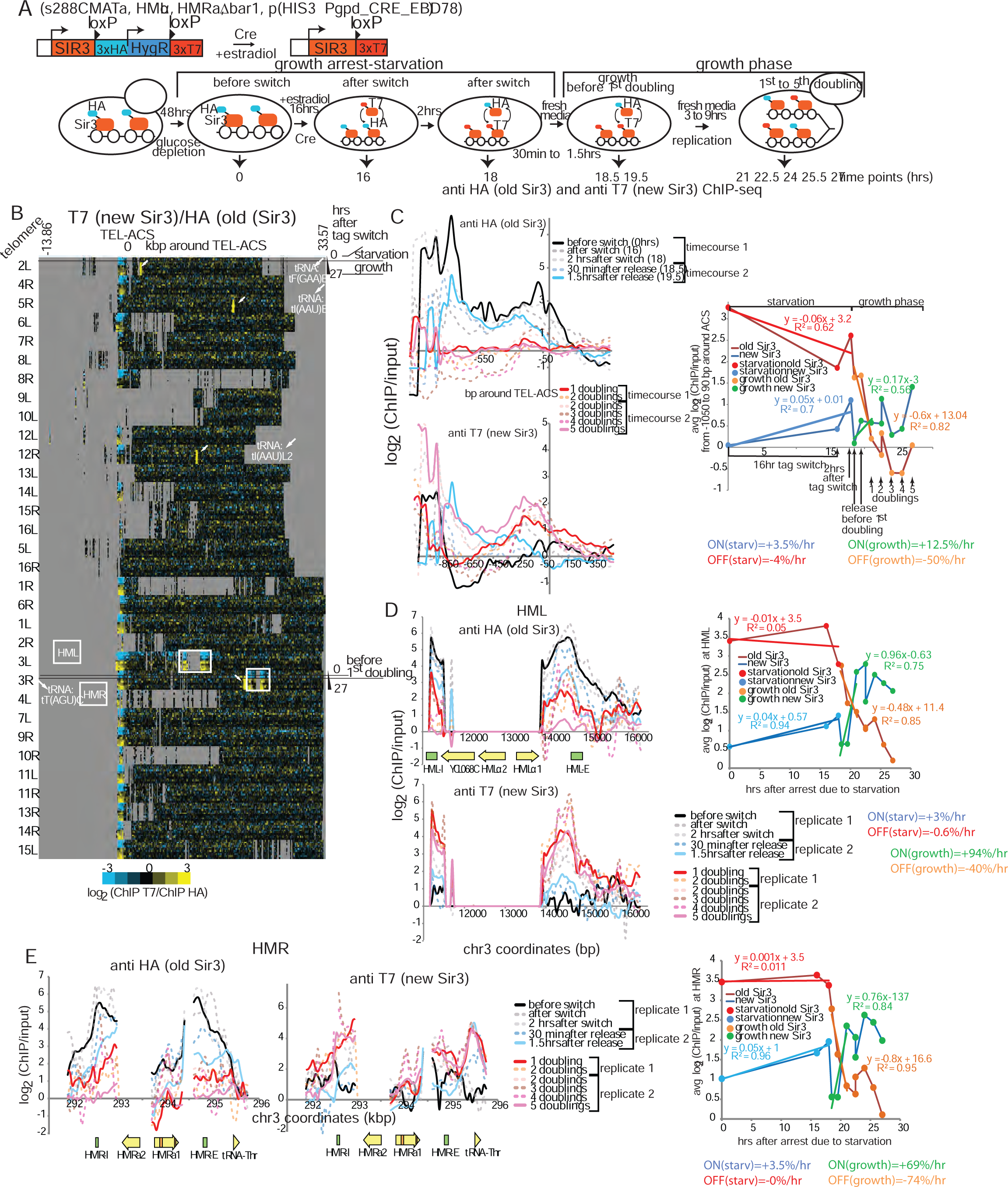
Rapid Sir3 turnover after exit from starvation induced growth arrest. **A.** Diagram of the Sir3 tag switch construct (top left). Bottom: Experiment outline. Cells were arrested by glucose depletion before the tag switch, induced with estradiol addition (recombination efficiency: 98.1%). Cells were then released from arrest with addition of fresh media and allowed to grow for one to five doublings (monitored by OD measurements). Parallel anti-HA and anti-T7 (polyclonal) ChIPs were then performed with cell aliquots that were fixed at times indicated below the diagram. **B.** Heat map of new Sir3 (T7 tag) enrichment over old Sir3 (HA tag) during and after exit from growth arrest, at all yeast telomeres (30 kbp from chromosomes ends). Time points are aligned by the ARS Consensus Sequence (TEL-ACS) located in telomeric silencer regions of the XCS (SIR complex nucleation sites). White arrows show tRNA genes where new Sir3 binds after exit from growth arrest. New T7 tagged Sir3 appears to be significantly enriched at all but 20 tRNA genes immediately upon release from growth arrest. The biological significance of this binding is not clear as the replicate experiment from Supplementary Figure S6 using a different anti-T7 antibody (a monoclonal one versus the polyclonal used here), while confirming Sir3-T7 binding to tDNAs, shows enrichment levels that are 8 to 16 fold lower than the ones measured in this experiment. Silent mating type loci HML and HMR, on 3L and 3R, respectively, are framed with a white rectangle. Sir3 is enriched in a small 1kb region upstream of the TEL-ACS at all telomeres. Repetitive and unmapped regions are shown in grey. The HMLα reads have been eliminated as repetitive sequences during alignment to the reference genome which is MATα. **C.** Old (top left) and new (bottom left) Sir3 enrichment around TEL-ACS averaged for all 32 telomeres at indicated time points during starvation arrest and the renewed growth phase. The right panel shows average enrichment around the TEL-ACS for old and new Sir3 over time with Sir3 on and off rates during growth arrest and during the first cell cycle after release calculated from the slope of the linear fit as shown. **D-E.** Old and new Sir3 enrichment at HML (D) and HMR (E) at indicated time points (same color code for D and E) during starvation and re-growth after release. The right panel shows average enrichment over the entire silent mating type locus for old and new Sir3 over time with on and off rates as in C. The time points marked replicate 1 and 2 come from two different time-course experiments.

Our measurements of Sir3 ON and OFF rates (see **Materials and Methods** in the SI Appendix) show that the old Sir3 stays bound to chromatin in arrested cells even after the tag switch because of a slow OFF rate of 4%, 0.6% and 0% decrease in Sir3-3xHA enrichment per hour and a slow ON rate of 3.5%, 3% and 3.5% increase in Sir3-3xT7 enrichment per hour for subtelomeres, HML and HMR, respectively (**Figure 5C-E**). This results in very slow exchange rates of −1%/hr, +2.1%/hr and +3.5%/hr for subtelomeres, HML, and HMR respectively. Overall, Sir3 enrichment after 64 hrs of starvation is decreased ∼2 to 4-fold relative to the genome average compared to the pre-arrest mid-log phase as shown in the replicate experiment in **Supplementary Figure S5B-C**. As previously observed in the ChIP-qPCR experiment (**Figure S3**), old Sir3 completely disappears by the first cell doubling after release into fresh media and is replaced by new Sir3, due to an increase in Sir3 enrichment ON rates to +12.5%/hr (from +3.5%/hr), +94%/hr (from +3%/hr) and +69%/hr (from +3.5%/hr) and an increase in OFF rates to −50%/hr (from −4%/hr), −40%/hr (from −0.6%hr) and −74%/hr (from −0%/hr) at subtelomeres, HML and HMR, respectively, compared to Sir3 exchange rates in arrested starved cells (**Figure 5C-E**). Also of note is that the first appearance of significant new Sir3 enrichment at heterochromatic loci at the 90min time point after release from starvation (**Figure 5C-E**) is consistent with the absence of transient Sir3 contacts with euchromatic genes for the first 90min after release from starvation (**Figure 3A**).

The enrichment in new Sir3 reaches steady state after the first division after release resulting in lower Sir3 occupancy relative to values before starvation. New Sir3 enrichment after the 12^th^ doubling is still 3 fold lower than old Sir3 immediately after release and ∼7 fold lower than Sir3 in mid-log cells, most probably due to low cellular levels of the hypo-morphic Sir3-3xT7 protein (**Figure 5B-C and Supplementary Figure S4B-C**).

The observed pattern of Sir3 binding dynamics after exit from starvation suggests that old Sir3 is rapidly removed from heterochromatin when cells resume growth before the first cell division and is completely replaced with new Sir3 by the end of the first cycle after release. Sir3 replacement at silent mating type loci is faster than at subtelomeric loci, probably because of the stronger SIR complex nucleation capacity of HML and HMR silencers compared to subtelomeric silencers (**Figure 5D-E, Supplementary Figure S5D-E**). Subtelomeric heterochromatin is consequently a highly dynamic structure with fast Sir3 exchange rates in optimal growth conditions.

We saw that the triple tagged Sir3 RITE construct has a hypo-morph phenotype with a 7 to 15 fold lower Sir3 enrichment in subtelomeric heterochromatin of mid-log cells than the wt untagged Sir3 (**Figure S4**). We also saw that the defect in Sir3 binding to nucleation sites conferred by the triple T7 tag caused a 1.7 fold decrease in the Sir3 ON rate at HMR-E compared to Sir3 with a single HA tag (**Figure S3**). We therefore used the same oeSir3 strain with a *SIR3* construct with a 3xHA tag under the control of the galactose inducible GAL1 promoter as in Figure 4E and (Radman-Livaja et al. 2011) to test whether *SIR3* overexpression can rescue the observed defect in Sir3 ON rates and Sir3 occupancy conferred by the triple HA tag (**Figure 6**). Cells were first allowed to grow exponentially in galactose to induce a 128 fold increase in SIR3 mRNA expression (**Figure 3D**). Cells were then arrested due to galactose depletion.

**Figure 6:**
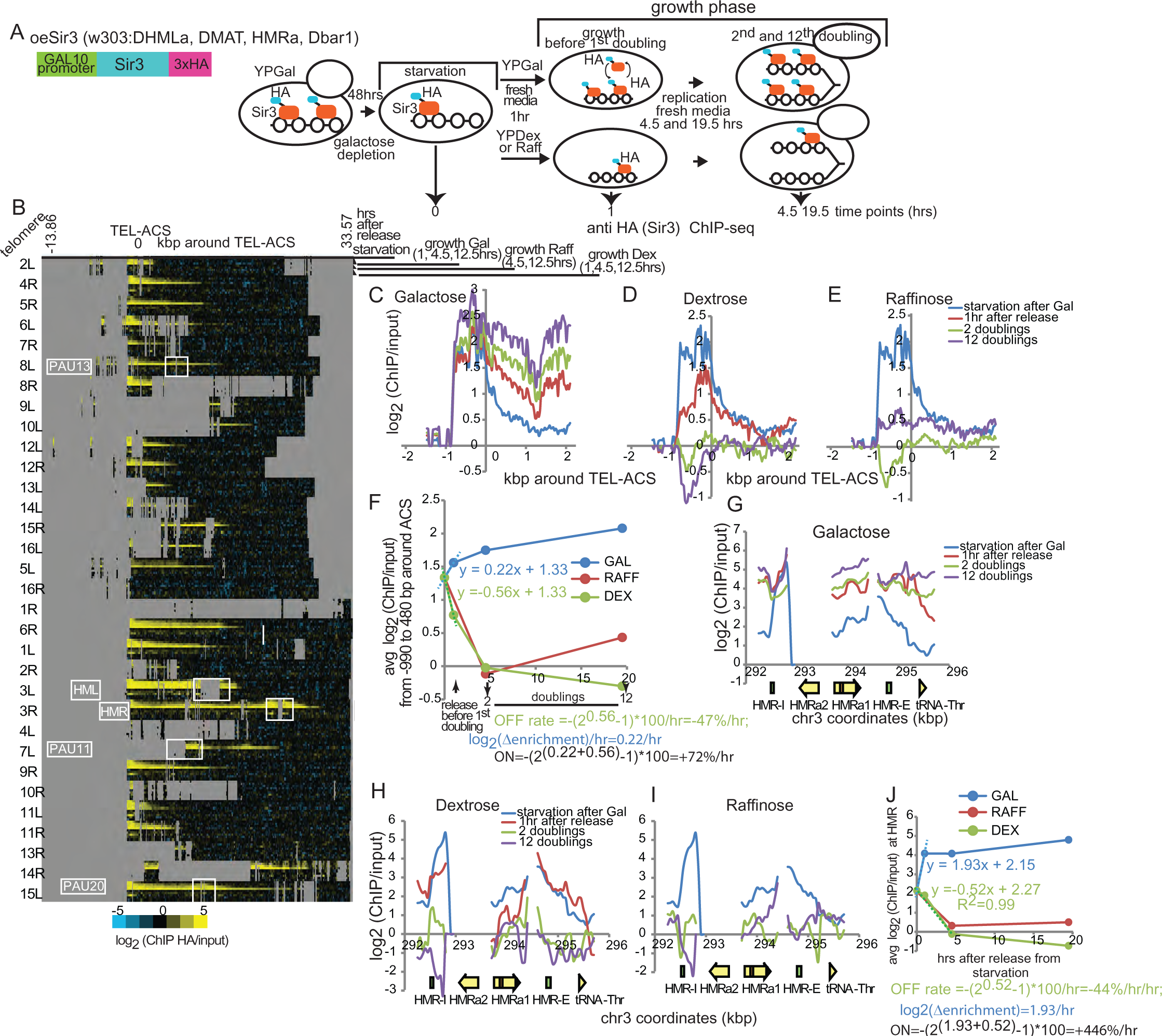
Sir3 turnover in condition of SIR3 overexpression A. Diagram of the Sir3 gene construct controlled by a Galactose inducible promoter in the oeSir3 strain (OverExpression) (top left). Bottom: Experiment outline. Cells were arrested after galactose depletion, released into fresh media with the indicated carbon source (2%), and allowed to grow for 2 and 12 doublings (monitored by OD measurements). Anti-HA ChIPs were then performed with cell aliquots fixed at times indicated below the diagram. **B.** Heat map of Sir3 (HA tag) enrichment over input during and after exit from starvation, at all yeast telomeres (30 kbp from chromosomes ends). Time points are aligned by the ARS Consensus Sequence (TEL-ACS) in XCS regions as in Figure 4B. Silent mating type loci HML (HML is deleted in this strain) and HMR, on 3L and 3R, respectively, are framed with a white rectangle. Repetitive and unmapped regions are shown in grey. **C-E.** Sir3 enrichment around TEL-ACS averaged for all 32 telomeres after release into Galactose-over expression of Sir3 (C), Dextrose-inhibition of Sir3 expression (D) or Raffinose-low Sir3 expression (E). **F.** Average Sir3 enrichment around the TEL-ACS over time in indicated carbon sources. **G-I.** Sir3 enrichment at HMR after release into Galactose (G), Dextrose (H) or Raffinose (I). **J.** Average Sir3 enrichment over the entire HMR over time.

Since galactose depletion stops Sir3 production and dextrose represses the GAL1 promoter, we can estimate genome-wide Sir3 ON and OFF rates after exit from growth arrest in conditions of *SIR3* overexpression simply by performing two parallel ChIP-seq time-courses in cells released from arrest into galactose or dextrose media, respectively.

Curiously, the 128 fold increase in Sir3 mRNA expression in the oeSir3 strain does not result in a proportionally higher Sir3 enrichment at SIR nucleation sites compared to the RITE Sir3-3xHA strain (compare 12^th^ doubling points in galactose from **Figure 5 C-H** to mid-log points in **Figure S5 C-E**), suggesting that the triple HA tag in oeSir3 is also interfering with Sir3 nucleation similar to the 3xHA tag in the RITE strain. Nevertheless, Sir3 over-expression does increase Sir3 ON rates as expected, but compared to the increase in mRNA expression levels, the effect is relatively modest with ∼6 fold higher ON rates at subtelomeres and HMR, for the overexpressed Sir-3XHA construct compared to the RITE switch Sir3-3xT7 (compare **Figures 5 and S5 C-E with 6 F-J**). This increase in ON rates is nevertheless sufficient for Sir3 enrichment to rapidly reach steady-state. In contrast to the RITE Sir3-3xT7 construct, Sir3 enrichment in the oeSir3 strain has already reached equilibrium 1 hr after release into galactose, suggesting that SIR complex renewal does not directly depend on DNA replication or cell division but is likely driven by the rate of Sir3 synthesis. As previously observed in mid-log cells (Radman-Livaja et al. 2011), the most dramatic consequence of *SIR3* over-expression after exit from growth arrest is the extension of subtelomeric Sir3 binding domains by as much as 15 kbp by the 12^th^ doubling after release from starvation, suggesting that low Sir3 levels in the wt and RITE strain, which are controlled by the endogenous Sir3 promoter, are the main factor that limits Sir3 spreading from subtelomeric nucleation sites (**Figure 6B**). Furthermore, our experiments with the oeSir3 construct also confirm that Sir3 polymerization is not impaired by the triple HA tag, as observed previously for the RITE 3xHA tag (**Figure S4**).

Sir3 OFF rates on the other hand are comparable between cells with wt Sir3 levels and cells with Sir3 over-expression (compare **Figures 5 and S5 C-E with 6 F-J**). It is however important to note that the process of old Sir3 removal occurs independently of Sir3 synthesis as evidenced by the complete disappearance of chromatin bound old Sir3 by the second doubling upon release into dextrose or raffinose in which Sir3 is either not produced or produced at low levels, respectively (**Figure 6H-I**). In other words the removal of old Sir3 from heterochromatin is not “driven” by its replacement with new Sir3.

### Thousands of genes restart transcription faster in the absence of Sir3

Transitions in and out of stationary phase or starvation induced growth arrest elicit dramatic changes in the cellular transcription program (Gasch et al. 2000; Martinez et al. 2004; Radonjic et al. 2005). Our experiments on the effect of Sir3 turnover on SIR complex function (**Figure 4**) also suggest that heterochromatin is transiently destabilized and heterochromatic genes are temporarily de-repressed immediately after exit from growth arrest. Moreover, Sir3 Nanopore-MetID revealed that Sir3 transiently targets at least a thousand euchromatic genes in exponentially growing cells. We consequently decided to investigate how Sir3 dynamics during and after starvation affect genome-wide gene expression levels.

In order to be able to directly compare mRNA levels between time points and different yeast strains, we performed duplicate RNA-seq experiments with external “spike in” normalization using RNA from *S. pombe* as described in Materials and Methods (**Figure 7**). We decided to take advantage of the Sir3 triple tag hypo-morph phenotype with reduced Sir3 enrichment at heterochromatic loci and measured mRNA dynamics during exit from growth arrest in the following strains: two WT strains with untagged Sir3 (WT1 and WT2), the Sir3-3xHA RITE strain before the tag switch, the Sir3-3xT7 RITE strain after the tag switch, the *sir2Δ*, *sir4Δ* and *sir3Δ* strains, and the *SIR3* over expression strain (oeSir3).

**Figure 7:**
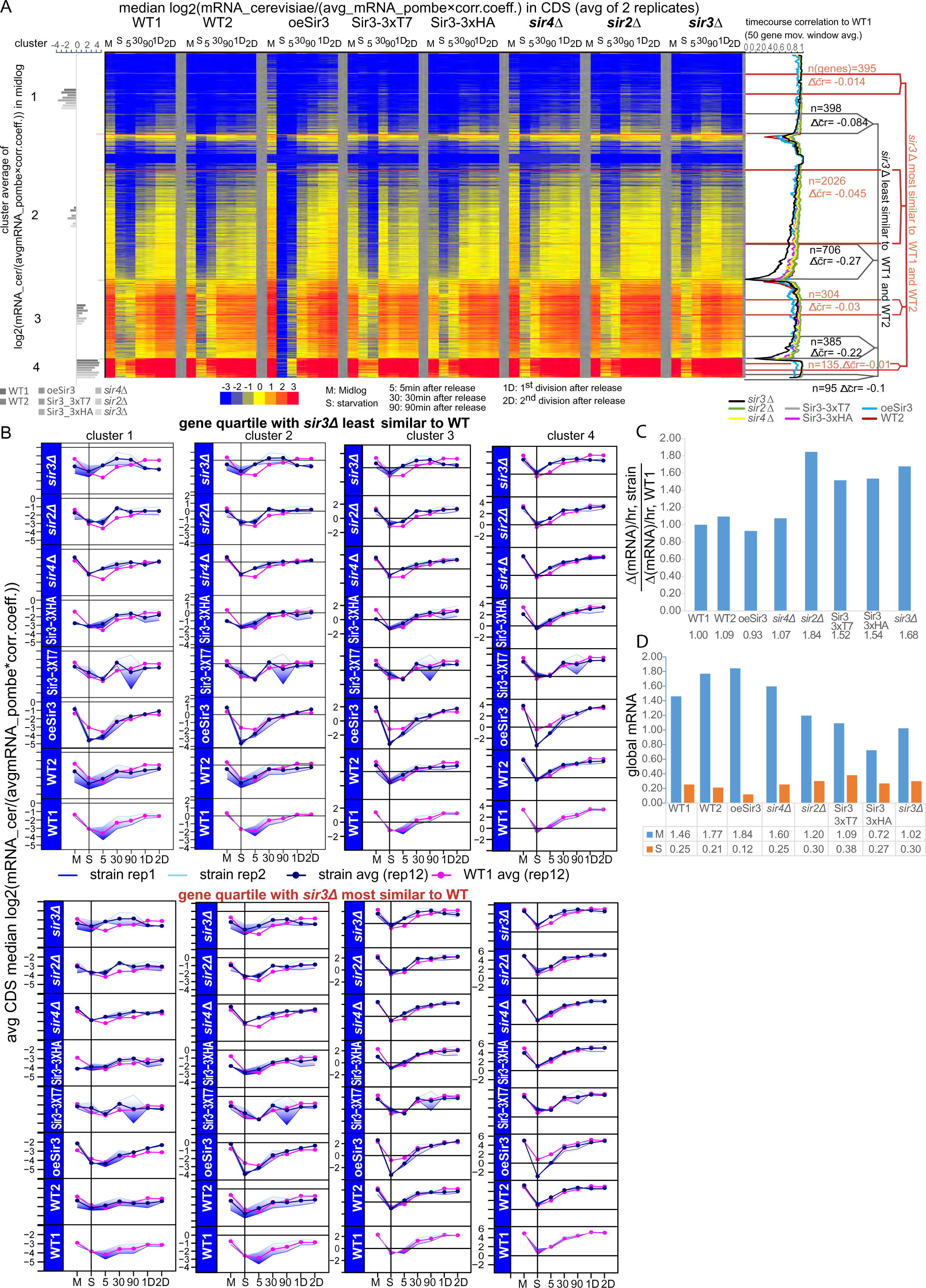
Genome-wide transcription is reactivated faster after exit from growth arrest in Sir3 and Sir2 mutants. **A.** Heat map of median mRNA enrichment averaged from two biological replicates (normalized to an external spike in control of *S. pombe* RNA as described in Supplemental Figure S8 and Materials and Methods in the SI Appendix) in indicated strains before and after release from growth arrest for ∼5800 yeast genes. Genes were grouped into four clusters based on average mRNA abundance in mid-log WT1 cells (bar graph on the left). The Pearson correlation between the time course expression profile in each strain and the WT1 strain was determined for each gene. Genes in each cluster were then ordered by increasing difference between the *sir3Δ* correlation to WT1 and the WT2 correlation to WT1 (Δcorrelation(*sir3Δ*-WT2)). The gene quartile within each cluster with the most negative Δcorrelation(*sir3Δ*-WT2) was considered to contain genes whose expression profiles between Sir3Δ and WT cells were the least similar (marked in black). The gene quartile within each cluster with the Δcorrelation(*sir3Δ*-WT2) closest to 0 was considered to contain genes whose expression profiles between *sir3Δ* and WT cells were the most similar (marked in red) **B.** Gene expression profiles were averaged for most and least similar gene quartiles in each cluster. The average profile from two biological replicates of each strain indicated in the strip header on the left (dark blue) is compared to the average expression profile of WT1 (magenta). The shaded blue area around the curve is delimited by average expression values from replicate 1(light blue) and replicate 2 (blue). **C.** The bar graph shows the rate of increase of global mRNA levels after exit from starvation for all analyzed strains normalized to WT1. It was calculated from the slope of the linear fit of average global mRNA read counts (average of two replicates from Supplementary Figure S7D) versus time (0, 5 and 30min after release from starvation). **D.** The bar graphs show the average global mRNA read counts from two biological replicates in Mid-log (M) and during Starvation (S) from Supplementary Figure S7D for all analyzed strains normalized as described in Supplementary Figure S7.

In order for “spike in” normalization to work, we need to add the same amount of *S. pombe* mRNA relative to *S. cerevisiae* mRNA for each time point from each strain. We therefore calibrated *pombe* mRNA levels relative to *cerevisiae* mRNA levels by mixing total RNA extracts at a 10:1 weight to weight ratio of *cerevisiae* to *pombe*, before the mRNA purification step. We opted for total RNA calibration instead of the alternative method, which consists of mixing cerevisiae and pombe cells at a constant cell count ratio prior to RNA extraction, because we wanted to avoid the variability between samples that could potentially be introduced by mechanical disruption of the cell wall by bead beating. Since starvation conditions change the composition and structure of the cell wall and since seriPAUperin genes, coding for cell wall mannoproteins, are more expressed during starvation and in SIR mutants (Ai et al. 2002), we reasoned that it would be technically very difficult to control for the variability in the efficiency of cell wall disruption by bead beating between different time points and strains. This could then introduce systematic errors in pombe spike-in normalization. We consequently preferred to use total RNA calibration because measurements of the total RNA amount by Qubit fluorescence assays or by Nanodrop spectrometry, and of its composition by BioAnalyzer (Agilent) or LabChIP (Perkin-Elmer) are accurate and reproducible. Since mRNAs represent only ∼5% of total RNA and since total RNA is ∼80% rRNA, we are essentially using rRNA to calibrate the cerevisiae to pombe total RNA ratio. **Figure S6** shows that the rRNA fraction does not vary significantly during the time course in all examined strains. We nevertheless used the information on RNA content obtained from BioAnalyzer/LabChip assays from each total RNA sample to correct for the small differences in rRNA content between different time points and strains by multiplying the measured genome wide average *pombe* mRNA read density with the correction coefficient calculated as shown in Figure S6 before *S. pombe* spike in normalization (see Materials and Methods). We also performed a second normalization to the genome-wide average *S.cerevisiae* to *S.pombe* mRNA ratio for the whole time course to account for variability in the total RNA *cerevisiae* to *pombe* ratios between replicates and strains (**Figure S7**).

We identified genes whose gene expression trajectory during the time course is most dependent on the presence of Sir3. First, we divided genes into four clusters according to their average mid-log gene expression levels in the WT1 strain. Second, we determined the Pearson correlation between the gene expression time course of WT1 and every other tested strain for each gene. Finally, in order to find the genes whose gene expression profiles in *sir3Δ* cells were the least similar to WT, we calculated the difference between the *sir3Δ*|WT1 and WT2|WT1 correlations for each gene (Δcorrelation(*sir3Δ*-WT2)).We then ordered genes in each cluster by their Δcorrelation(*sir3Δ*-WT2). Genes in the quartile with the most negative average Δcorrelation(*sir3Δ*-WT2) in each cluster are the ones whose gene expression across the time course is the least similar between *sir3Δ* cells and both WT1 and WT2 cells. Conversely, the quartile from each cluster with an average Δcorrelation(*sir3Δ*-WT2) that is closest to 0 contains genes whose expression in *sir3Δ* cells is the most similar to either WT strain (**Figure 7A**).

We found 1584 genes whose gene expression dynamics after exit from growth arrest are the most affected by the absence of Sir3 and Sir2 or by reduced amounts of Sir3 in Sir3-3xHA|T7 RITE strains. The expression profiles of these genes averaged per cluster and per strain show that the main difference between WT and the other strains lies in the speed at which mRNAs return to mid-log levels after the drop observed during growth arrest (**Figure 7B**). It takes on average one whole cell cycle for mRNAs to go back up to mid-log levels in both WT strains. This happens much sooner in *sir3Δ* cells. Typically, mRNA levels are back to pre-starvation amounts only 5 min after release from growth arrest. In Sir3 hypo-morphs-Sir3-3xHA, Sir3-3xT7- and *sir2Δ* cells, mRNA levels reach a plateau a little later, at the 30 min time point, which is still 4 hrs before mRNAs level off in WT cells. The trajectory of gene expression re-activation in *sir4Δ* cells appears to by a hybrid between WT cells and *sir3* mutants, with an initial jump in mRNA levels at the 5min time point similar to *sir3* mutants followed by a more gradual rise to mid-log levels akin to wt cells. *oeSIR3* cells on the other hand follow similar kinetics of reactivation as wt cells despite starting from a ∼2 fold lower mRNA baseline during starvation. mRNAs also level off faster in SIR complex mutants and Sir3 hypo-morphs than in WT cells, in the quartiles with genes whose expression is the most similar between *sir3Δ* and WT cells.

In fact, the genome-wide average rate of gene expression re-activation is 60% higher in *sir3Δ*, *sir3* hypo-morphs and *sir2Δ* cells compared to WT, *oeSIR3* and *sir4Δ* cells (**Figure 7C**). Sir3 and Sir2 also participate in global gene expression regulation in starved and mid-log cells as evidenced by 0.3 fold higher or 0.6 fold lower genome-wide average mRNA levels during starvation and exponential growth, respectively, in Sir3 and Sir2 mutants compared to WT and *sir4Δ* (**Figures 7D** and **S7D**). Consistent with these observations, over-expression of Sir3 causes a 2-fold decrease and a 14% increase in global mRNA levels, during starvation and exponential growth, respectively. Consequently, it appears that the levels of Sir3 and Sir2 that are found in wt cells are not only needed to delay the resumption of full transcriptional activity till the end of the first cell cycle after release from growth arrest, but are also necessary for global transcription regulation during exponential growth and during growth arrest.

The simultaneous expression of a1 and α2 transcription factors from the de-repressed silent mating type loci creates a pseudo-diploid phenotype that silences haploid specific genes and activates diploid specific genes in Sir mutants(Fraser and Heitman 2003). There are three reasons why we are inclined to dismiss the possibility that the global effect on transcription that we see in the *sir3Δ* mutant is an indirect consequence of the pseudo-diploid phenotype of our strain. First, our results show that SIR3 deletion affects the transcription of all genes to different degrees while haploid and diploid specific genes that are directly controlled by the a1 and α2 transcription factors represent only a small fraction of all yeast genes. Indeed, Ellahi et al. (2015)(Ellahi et al. 2015) identified only 55 genes that were differentially expressed in pseudo-diploids (*sir2Δ, sir3Δ* and *sir4Δ*) compared to wt. Second, there should be no difference between *sir2Δ, sir3Δ* and *sir4Δ* mutants if the effect we saw was only due to a pseudo-diploid phenotype as they are all three pseudo-diploids. We however measure a much smaller effect of the *SIR4* deletion on genome-wide transcription reactivation, which resembles wt more than s*ir3Δ* or *sir2Δ* (Figure 7 C-D)*. sir2Δ* also does not behave exactly like *sir3Δ* and has an intermediate phenotype between wt and *sir3Δ* (Figure 7). Third, the two fold decrease in global mRNA production during starvation, observed in the oeSir3 strain cannot be due to a pseudo-diploid phenotype because HML and MAT loci are deleted in the oeSir3 strain.

GO annotation analysis shows that genes that are the most affected by suboptimal SIR complex levels are involved in protein transport and localization, nucleotide/nucleoside binding, and Golgi apparatus function. The least affected genes are mostly involved in transcription, RNA processing, ribosome function, and mitochondrial function and respiration (**Table S2**).

We also looked at subtelomeric genes whose expression is more likely to be directly affected by changing Sir3 levels. Consistent with previous reports (Ellahi et al. 2015), wt levels of the SIR complex do not control the expression of most subtelomeric genes because we found only 16 genes out of 132 in our RNA-seq datasets that were located within 15 kbp of subtelomeric SIR nucleation sites, whose gene expression profiles in SIR mutants and Sir3 hypo-morphs were significantly different from WT cells (Δcorrelation(Sir3Δ-WT2) ≤ −0.35, **Figure S8**). We observed that mid-log expression of some of these genes is somewhat higher in Sir3 mutants, but is overall not significantly different as observed before (Ellahi et al. 2015). All 16 genes are however more expressed in the *SIR3* mutant strains than in either WT strain during starvation and immediately after release from growth arrest. The increase in expression observed in *SIR3* mutants is only temporary and mRNA levels fall back to low mid-log levels by the 1^st^ division after release. It consequently seems that the proximity of the SIR complex is not a determining factor for the low expression of subtelomeric genes in optimal growth conditions. The SIR complex is instead needed to slow down transcription reactivation after release from starvation. We speculate that controlled transcription reactivation may be needed to delay “full-blown” genome-wide transcription until the cell is ready for a complete re-establishment of its transcription program.

SIR dependent attenuation of transcription of at least some subtelomeric genes in growth arrest and after exit from growth arrest might have been expected because of their proximity to SIR bound loci. The fact that thousands of euchromatic genes located far away from “canonical” SIR loci exhibit a similar SIR dependent delay in transcription re-activation, is however more puzzling.

One possible explanation for the faster reactivation of transcription in the absence of the SIR complex that is consistent with higher mRNA levels during starvation in Sir2 and Sir3 mutants (**Figure 7D**), is that transcriptional activity does not fully shut down during starvation in these mutants and might therefore be able to restart faster upon release from arrest. If there was still some lingering low-level genome-wide transcription during growth arrest, we reasoned that the apparent mRNA half-lives in SIR mutants and hypo-morphs would be longer than in WT cells. Conversely, Sir3 overexpression would cause a more efficient shutdown of transcription during starvation, resulting in lower global mRNA amounts as observed in Figure 7D and shorter apparent mRNA half-lives than in wt cells. For simplicity sake, we assumed that the decay pattern for most mRNAs is exponential, even though it is probably more complex for a number of mRNAs (Deneke et al. 2013). Of course, our prediction would only hold if the true mRNA decay rates are the same in all tested strains, which we assumed should be the case because decay rates are for the most part determined by mRNA sequence and secondary structure, which are identical in all strains (Miller et al. 2011). We consequently determined apparent mRNA half-lives during growth arrest using the datasets from Figure 7.

We detect a genome-wide drop in mRNA abundance during growth arrest in all tested strains, as would be expected if global transcription activity has largely stopped during that period (**Figures 7D** and **S9**). We calculated apparent mRNA decay rates and half-lives directly from the decrease in mRNA levels from mid-log to growth arrest, as shown in **Figure S9A**. Somewhat surprisingly and in contrast to decay rates in mid-log cells(Miller et al. 2011), mRNA decay rates during starvation are directly correlated with mid-log mRNA levels: the more an mRNA was abundant before growth arrest the shorter its half-life during growth arrest (**Figure S9**). As predicted above, apparent mRNA half-lives are globally ∼60 to 70% longer in Sir3Δ and Sir3 hypo-morphs compared to wt, and they are two times shorter in the Sir3 over expression strain but they are only marginally affected in Sir2 and Sir4 deletes (**Figure S9B**).

Most importantly, the Sir3 dependent global modulation of transcription has physiological significance as demonstrated by a competitive fitness assay in **Supplementary Figure S10**. Sir3 gives a twofold growth advantage to WT cells compared to *sir3Δ* mutants when cells have to compete for resources during starvation (**Figure S10B, C**). *sir3Δ* cells are on the other hand not competed out by WT cells in optimal growth conditions during exponential growth (**Figure S10A, C**). Note that the predominance of WT cells in starvation conditions is not due to increased lethality of starved *sir3Δ* cells because both strains resume growth at comparable rates when starved cells are diluted down to the same OD with fresh media (**Figure S10B**). Most importantly, the growth advantage of wt cells is not due to the pseudo-diploid phenotype of *sir3Δ* because wt cells have no competitive advantage over *sir4Δ* cells that also exhibit a pseudo-diploid phenotype but unlike *sir3Δ* cells don’t have a significant impact on transcription efficiency (**Figure S10C**).

In conclusion, transcription dynamics in SIR complex mutants before, during and after exit from starvation is consistent with the idea that Sir3 globally stimulates transcription in exponentially growing populations (Figures 7C and S7D) and is involved in the global downregulation of transcription during nutrient deprivation (Figures 7D, S7D and S9B). Higher residual transcription during starvation in Sir3 mutants suggests that the transcription machinery remains bound to genes at higher levels in the absence of Sir3, which facilitates a more rapid transcription reactivation when nutrients are restored upon release.

### A model for global modulation of transcription efficiency by Sir3

Our RNA-seq results suggest that Sir3 regulates the dynamics of global reactivation of transcription after exit from growth arrest caused by nutrient starvation. Remarkably, more than 99% of genes whose reactivation is delayed in the presence of Sir3 are located in euchromatin far away from previously known Sir3 binding sites and the vast majority of affected genes are not known to be directly or indirectly controlled by the SIR complex. This raises the intriguing possibility that Sir3 regulates gene expression through transient direct contacts with euchromatic genes, revealed by Sir3 Nanopore-MetID (**Figure 2**).

The absence of genome wide Sir3 contacts after release from starvation (**Figure 3A**) suggests that the slower reactivation of transcription after release from starvation in WT compared to Sir3 mutants may be due to the genome-wide binding activity of Sir3 before starvation induced growth arrest.

Our results are consistent with the hypothesis that a so far unknown Sir3 activity resulting from its genome-wide contacts with euchromatic genes during exponential growth prior to nutrient starvation influences transcription activity during and after release from starvation. We consequently wanted to see if this putative Sir3 activity was different between genes whose transcription after exit from starvation is the most or the least affected by Sir3 mutations, or if the same Sir3 activity had a different effect on transcription dynamics of these two groups of genes. We consequently looked for differences in the density and/or probability of Sir3 contacts with euchromatic genes identified in Figure 2, depending on whether these contacts are found in proximity of genes whose transcription after exit from starvation is the most or the least affected by Sir3 mutations, i.e., whose transcription dynamics in *sir3Δ* are least or most similar to WT, respectively (see Figure 7).

The set of 1197 Sir3 targets identified in Figure 2 overlaps by ∼20% with sets of genes whose transcription is the most (1579 genes, Group 2) or the least (2855 genes, Group 1) affected by Sir3 mutations, respectively (**Figure 8B**). As mentioned previously, since Sir3 binding to euchromatin is transient we probably missed a number of Sir3 targets whose residency time is too short to be reliably detected by Nanopore-metID. Considering that the transcription dynamics of practically all yeast genes appear to be affected by Sir3 mutations albeit to different degrees, the number of identified Sir3 targets is probably an underestimate of the actual number of transient Sir3 contacts with euchromatic genes.

**Figure 8:**
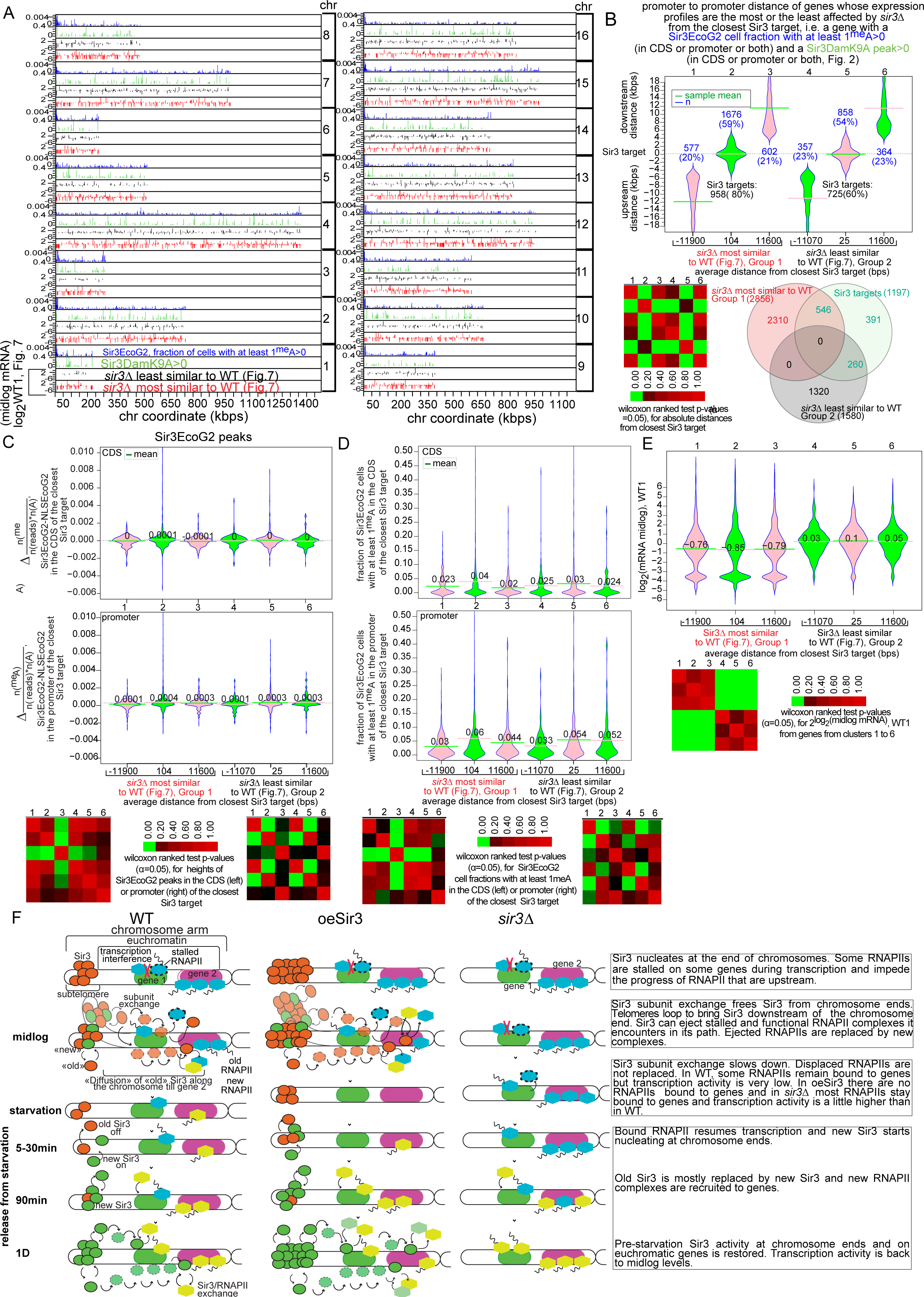
Model of Sir3 mediated global turnover of transcription complexes. **A.** Chromosome maps of: 1. WT1 mid-log mRNA levels of genes least affected by a Sir3 deletion (most similar to WT, red) 2. WT1 mid-log mRNA levels of genes most affected by a Sir3 deletion (least similar to WT, black) 3. Median Sir3DamK9A peak height (Δ(n(^me^A) /(n(reads)*n(GATC)), Sir3DamK9A-NLSDamK9A) of genes with Sir3DamK9A>0 in its CDS or promoter, whichever is higher (green). 4. Median Sir3EcoG2 peak height (Δn(^me^A) /(n(reads)*n(A), Sir3EcoG2-NLSEcoG2) of the promoter or CDS (whichever is higher) of genes with a cell fraction with at least 1^me^A >0(blue). **B.** Density plots of distances between promoters of genes that are least (clusters 1-3, Group 1) or most (clusters 4-6, Group 2) affected by *sir3Δ* and the closest gene from the 1197 gene set of Sir3 targets from Figure 2. Clusters 2 and 5 contain genes that are found within half a Standard Deviation from the mean upstream or downstream of the closest Sir3 target. Since the Wilcoxon ranked sum test excludes negative values, the minimum value in each cluster was subtracted from each value in the cluster to convert all values to positive values. **C.** Density plots of Sir3EcoG2 peak heights (^me^A density) in the CDS (top) or the promoter (bottom) of the gene from the 1197 gene set contacted by Sir3 that is closest to each gene from clusters 1-6 from B. **D.** As C but for the fraction of cells with a Sir3EcoG2 methylated CDS (top) or promoter (bottom) of the closest Sir3 target. **E.** density plots of WT1 mid-log mRNA levels (Figure 7 and A) of genes in clusters 1 to 6 from B. **F.** A model of the effect of Sir3 contacts on the transcription of underlying genes in mid-log and after exit from starvation. Note that telomeres are shown to loop only in mid-log cells, for clarity sake, we expect that telomere looping probably occurs during and after release from starvation as well.

Similar to what we saw for the localization of Sir3 targets along the chromosome, genes from Groups 1 and 2 are distributed homogenously on each chromosome (**Figure 8A**). Genes from Group 1 are interspersed with genes from Group 2 and do not appear to cluster away from each other in specific locations on the chromosome. To further characterize the relationship between Sir3 contact sites and genes from Groups 1 and 2 (least and most affected by *sir3Δ*, respectively), we sorted genes from Groups 1 and 2 by their promoter-to-promoter distance from the closest gene contacted by Sir3 from the set identified in Figure 2 (**Figure 8B**). More than 50% of the genes from the least affected (cluster 2 in Group 1) or the most affected group (cluster 5 in Group 2) are found within 5kbp of the closest gene contacted by Sir3, with an average distance of ∼1kbps, i.e. more than half of genes from groups 1 and 2 appear to be direct Sir3 targets. There are also no statistically significant differences between clusters that contain genes that are more than 4kbps away from Sir3 targets (compare clusters 1 and 3 in Group 1 with clusters 4 and 6 in Group 2) nor between clusters 2 and 5 that contain genes from Group 1 and 2 that are directly contacted by Sir3, respectively. This analysis shows that genes from either group are equally likely to be found within or close to regions contacted by Sir3. Likewise, there are no significant differences in Sir3 contact probability or frequency in the cell population between Sir3 targets that overlap with Group 1 genes (least affected by *sir3Δ*) or Group 2 genes (most affected by *sir3Δ*) (**Figure 8C-D**). Sir3 contacts tend to be somewhat more frequent (the peaks are higher on average) in regions close to genes from clusters 2 and 5 but this is expected since these clusters include Sir3 targets that are close to chromosome ends, where Sir3 targets are more densely packed and have high Sir3 contact probabilities and Sir3 densities. There is however no statistically significant difference in Sir3 contact probability or density between clusters 2 and 5, again suggesting that Sir3 exerts the same level of activity on genes in Groups 1 or 2.

Why does then a Sir3 mutation, have a quantitatively different effect on genes in Group 1 compared to genes in Group 2? A comparison of mid-log mRNA levels between the two groups provides a clue (**Figure 8E**). Genes that appear least affected by a Sir3 deletion (clusters 1-3, Group 1) are enriched for genes with either very low (log_2_(mRNA)< −3) or very high (log_2_(mRNA)>3) expression, while genes that appear more affected by *sir3Δ* (clusters 4-6, Group 2) are mostly moderately transcribed (−2<log_2_(mRNA)<2). The analysis in Figure 8 shows that all genes are equally likely to be contacted by Sir3 but the effect of a Sir3 contact is more pronounced and therefore more “visible” on moderately transcribed genes.

We now propose a model for the genome-wide Sir3 activity that takes into account all the results presented in this study (**Figure 8F**). In our model, subtelomeric regions are reservoirs of Sir3 from which Sir3 molecules are brought over to distal chromosomal regions through random looping of the chromosomes arm, and then released onto euchromatic genes thanks to a steady turnover of Sir3 subunits within the SIR complex at chromosome ends. Sir3 molecules thus come randomly and transiently into contact with genes along the chromosome arm. While we do not have direct evidence that Sir3 impacts RNAPII occupancy on transcribed genes, our RNA-seq and nanopore-MetID data are consistent with the idea that Sir3 improves transcription efficiency in mid-log cells by displacing stalled transcription complexes from chromatin when it transiently comes into contact with them, as discussed below.

## Discussion

Our study sheds light on two long standing questions in yeast heterochromatin biology: 1. How are heterochromatic structures maintained in response to changes in growth conditions and 2. What is the cellular function of subtelomeric heterochromatin.

Our measurements of Sir3 dynamics within heterochromatin after exit from growth arrest show a dramatic increase in Sir3 exchange rates, as well as Sir3 degradation rates immediately after release from arrest (**Figures S3, 5** and **S5,** and **Figure 3**). We show that Sir3 subunits exchange very slowly during growth arrest and Sir3 protein degradation is also slowed down relative to exponentially growing cells (**Figure 3**). The supply of fresh nutrients triggers the reactivation of all cellular processes including faster Sir3 protein degradation and faster Sir3 turnover within the SIR complex. Consequently, all Sir3 proteins bound to subtelomeric and silent mating type loci are eventually replaced with newly synthesized Sir3 by the end of the first cell cycle after release. We demonstrate for the first time that budding yeast heterochromatin is a highly dynamic structure that is continuously renewed throughout the cell cycle. Consequently, high Sir3 turnover rates in the first cell cycle after release from growth arrest don’t allow us to conclude if Sir3 is epigenetically inherited and if it facilitates the reconstitution of the SIR complex after replication. The RITE tag switch system cannot be used to differentiate between replication dependent and replication independent Sir3 dynamics in exponentially growing cells because Cre mediated recombination of epitope tags in at least 95% of cells takes 16 hrs, during which time cells double 8 to 11 times. A different strategy that relies on fast and temporally controlled labelling of Sir3 should therefore be used to assess Sir3 dynamics in exponentially growing cells, but we suspect that Sir3 exchange rates in exponentially growing cells are not much different from the ones we measured in the first cycle after release from starvation.

Even though the RITE strain with a Sir3-3xHA to Sir3-3xT7 tag switch that we used to measure genome-wide Sir3 exchange rates is a hypo-morph with lower Sir3 enrichment at heterochromatic loci than in WT cells, we show that the genome-wide Sir3 OFF rates measured in this strain are not significantly different from Sir3 OFF rates measured in a strain that does not exhibit the “hypo-morph” phenotype (**Figure S3**). Sir3 OFF rates are therefore not affected by reduced Sir3 density at heterochromatic loci observed in the “hypo-morphs”. Sir3 ON rates are independent of OFF rates and are directly proportional to Sir3 dosage (compare **Figures S3, 5** and **S5** with **Figure 6**). We consequently conclude that Sir3 subunits “leave” the SIR complex at similar rates in the Sir3-3xHA to 3xT7 RITE strain as in WT cells but are replaced with new Sir3 subunits at slower rates in the hypo-morph strains than in WT.

The α-factor test for the gene silencing function of the SIR complex revealed that the sudden increase in Sir3 turnover rates upon exit from growth arrest that follows Sir3 depletion during starvation compromises SIR function at silent mating type loci in a small but significant fraction of the wt cell population. The rapid increase in Sir3 turnover after release from starvation appears to make the SIR complex relatively more permissive to transcription of silent mating type loci. If the supply of new Sir3 is not optimal as in the Sir3 hypo-morphs, SIR dependent silencing of silent mating type loci is 15 to 30 fold less efficient after exit from growth arrest and 200 to 500 fold less efficient in mid-log cells, compared to WT cells in the same conditions (**Figure 4**).

We found that the SIR complex directly represses only a small subset of subtelomeric genes (**Figure S8**). SIR complex spreading from subtelomeric nucleation sites does not however explain the repression of all documented genes as most of them are located beyond documented heterochromatin boundaries that are positioned ∼2kbps around nucleation sites (**Figure S8**). We propose instead that the enhanced Sir3 dependent repression of these genes immediately after release from starvation is the consequence of a thorough removal of transcription complexes by a Sir3 sweeping activity that is made more efficient by the proximity of SIR nucleation sites from which according to our model in Figure 8F, Sir3 is brought over to the rest of the chromosome.

The current generally accepted view is that the principal function of heterochromatin is to repress underlying genes whose expression would be potentially deleterious to cellular function and viability. The function of the SIR complex at silent mating type loci in haploid cells aligns itself with that canonical function of heterochromatin: it prevents the simultaneous expression of genes for both mating types and thus enables mating between cells of opposite type. The biological function of the SIR complex in subtelomeric regions has on the other hand long been somewhat of a mystery. Results from this study and others argue against a primary role in the silencing of subtelomeric genes through position effect variegation (Ellahi et al. 2015) in exponentially growing cells where the absence of Sir3 has a marginal effect on the expression of most subtelomeric genes (**Figures 7 and S8**).

Our analysis of gene expression dynamics using RNA-seq with *S. pombe* RNA “spike in” normalization now reveals wide-spread faster reactivation of transcription of almost all euchromatic genes, immediately after exit from growth arrest in SIR mutants and “hypo-morphs” (**Figure 7**). This result is all the more unexpected, since the vast majority of affected genes are not known to be directly or indirectly controlled by the SIR complex, as measured by ChIP-seq. More than a thousand euchromatic genes located throughout the genome were only now revealed as direct Sir3 targets thanks to our in vivo foot printing technique Nanopore-MetID (**Figures 1** and **2**). This led us to hypothesize that transient and sporadic Sir3 contacts with euchromatic genes, which cannot be detected by ChIP-seq, are involved in the global control of gene expression before and after exit from starvation. Sir3 localization to euchromatin is not unprecedented. ChIP-seq experiments have detected Sir3 at highly transcribed genes and euchromatic replication origins (Radman-Livaja et al. 2011; Thurtle and Rine 2014; Hoggard et al. 2018). While some of the Sir3 signal at highly transcribed genes may be attributed to a ChIP artefact (Teytelman et al. 2013), our Nanopore-MetID results suggest that a good number of these genes represent bona fide Sir3 targets.

How would Sir3 be “dispatched” throughout the genome? We propose that SIR nucleation sites at chromosome ends serve as Sir3 hubs where the continuous exchange of Sir3 subunits during exponential growth provides a steady stream of Sir3 molecules that are brought over to distal sites along the chromosome arm through telomere looping. This model is consistent with our Nanopore-MetID results, which show a gradual decrease in the density of Sir3 contacts that correlates with the distance from the SIR nucleation site and 50% to 60% overlap between Sir3 and Sir4 or Sir2 targets, respectively. The analysis from **Figure 2D** shows that ∼50% of genes are contacted by Sir3 in the zone up to 20kbps downstream from the subtelomeric SIR nucleation site. The density of Sir3 targets drops gradually to 20% in the following 20kbps and then stays at 20% beyond that, all the way up to the centromere. This distribution of Sir3 targets along chromosome arms would be consistent with a mechanism that delivers Sir3 to distal chromosomal regions through loop formation bringing subtelomeric regions in proximity of regions further away from chromosome ends. These loops may however be difficult to detect if they are short lived. Nevertheless, a few of these bridges have recently been identified by Hi-C in mid-log cells when Sir3 was overexpressed (Ruault et al. 2021). It would therefore be interesting to explore this possibility with a more sensitive method than Hi-C, like DamC (Redolfi et al. 2019).

In our model, subtelomeric heterochromatin does not perform a direct biological function, it serves instead as a reservoir of Sir3, which than performs its biological function away from subtelomeric heterochromatin and the SIR complex, at hundreds of distal sites throughout the chromosome. The importance of subtelomeric heterochromatin formation for the Sir3 dependent control of mRNA expression efficiency, is supported by Sir3EcoG2 Nanopore-MetID and spike in RNA-seq experiments in *sir2Δ* and *sir4Δ* strains (**Figures 1, 2 and 7**). Indeed, in the absence of Sir2 or Sir4, Sir3 binding to subtelomeric nucleation sites is, respectively, close to background levels or 2 fold lower than in wt cells (**Figure 1I**). While the drop in the number of euchromatic Sir3 targets in *sir2Δ* and *sir4Δ* is only 10% compared to wt (Figure 2D), the cell population frequency of Sir3 contacts in the subtelomeric SN1-SN4 regions identified in Figure 2E, is reduced 2 to 3 fold in *sir4Δ* compared to wt, while in *sir2Δ* the peaks completely disappear. The absence of Sir2 consequently, eliminates SIR complex formation while the absence of Sir4 merely impairs it. The difference in SIR complex formation between *sir4Δ* and *sir2Δ* is consistent with the differential effect of each mutation on gene expression with *sir4Δ* that still forms SIR complexes in subtelomeric regions albeit less efficiently, being closer to wt, and *sir2Δ*, in which SIR complexes are no longer detectable, being closer to *sir3Δ.* Our results therefore suggest that SIR complex formation at chromosome ends is the decisive factor that ensures optimal delivery of Sir3 to euchromatic genes (**Figure 2D**).

The next question is: what does Sir3 do once it contacts euchromatic genes? We propose that Sir3 acts as a genome-wide “sweeper” of the components of the transcription machinery and facilitates the turnover of transcription complexes at actively transcribed genes (**Figure 8F**). Note that we use RNAPII as proxy for any transcription associated factor or complex in the illustration in Figure 8F and in the text below. Our current working hypothesis is that Sir3 can evict any transcription associated protein, including transcription factors at promoters. While the displacement of a stalled RNAPII is more likely, because Sir3 is more likely to come into contact with a non-moving target, Sir3 should also be able to displace an actively transcribing RNAPII during its «travels».

In exponentially growing cells the displaced RNAPII can readily be replaced by new RNAPII complexes with minimal disruptions to global transcription. The «sweeping» of stalled RNAPIIs by Sir3 is actually beneficial because it allows for the RNAPIIs that are backed up upstream of the stalled complex to resume transcription, which would explain why global mid-log mRNA levels are on average ∼70% lower in Sir2 and Sir3 mutants compared to wt, oeSir3 and Sir4Δ strains (Figures 7D and S7D). Since Sir3 can potentially sweep away indiscriminately any RNAPII it contacts, moderately transcribed genes in wt cells will be mostly cleared of RNAPII after a period of starvation when the displaced RNAPIIs cannot be replaced. Cells will therefore require some time after release from starvation to repopulate their RNAPII levels and return to pre-starvation levels of activity. On highly transcribed genes with a high density of bound RNAPII, Sir3 would clear up a smaller fraction of bound RNAPII complexes than on moderately transcribed genes. Some of these complexes will stay bound to highly transcribed genes during starvation (and some of them will still be active) and allow for a more rapid reactivation after release from starvation. Consequently, Sir3 mediated sweeping of bound RNAPIIs will have a bigger effect on moderately transcribed genes than on highly transcribed genes. High levels of Sir3 in the oeSir3 strain will result in a more thorough sweeping of bound RNAPII, resulting in genes mostly depleted of RNAPII, thus explaining a 2 fold drop in residual transcription during starvation, and ∼14% higher transcription in mid-log compared to WT, due to more efficient displacement of stalled RNAPIIs. In the absence of Sir3 in the *sir3Δ* strain, all genes will have more bound RNAPII during starvation and transcription reactivation after release will be faster, as observed.

Our RNA-seq and Sir3Dam/EcoG2 Nanopore-MetID results suggest that the rate of transcription reactivation after release from starvation is influenced by genome-wide Sir3 binding during exponential growth before the onset of starvation. Indeed, Sir3 contacts with euchromatin are reduced below the detection threshold for at least 90 min after release from arrest or practically for the entire time it takes for the transcription program to get back to mid-log levels (**Figure 3A**). Consequently, it is while nutrients are still available and cells are still dividing that the Sir3 activity in euchromatin determines how fast genes will be reactivated after global transcription has been shut down because of nutrient shortage. So, Sir3 is affecting transcribing genes in a way that generally improves the efficiency of transcription genome-wide in exponentially growing cells (Figure 7D), but decreases residual transcription during starvation (Figures 7D and S9B) and finally slows down reactivation of transcription after release from starvation (Figure 7C). These processes respond to Sir3 dosage since mRNA levels are respectively higher or lower in mid-log or during starvation, and transcription reactivation is slightly slower after release in oeSir3 cells relative to wt. Our hypothesis that Sir3 acts as a general purpose “sweeper” of transcription complexes would be consistent with all these observations. Sir3 would periodically and randomly clear out all or a fraction of bound proteins, depending on the expression level of the underlying gene. Since this kind of indiscriminate sweeping would also remove stalled transcription complexes it would generally improve transcription efficiency in conditions of optimal growth when new complexes are readily available to restart transcription. The complexes that were cleared out by Sir3 towards the end of the growth phase should not be replaced during starvation, which explains lower residual transcription in starved wt cells compared to *sir3Δ*. It would then take some time for new transcription complexes to rebind after nutrients are replenished, resulting in generally slower transcription reactivation. In the absence of Sir3 on the other hand transcription complexes should not be cleared out as efficiently and would still stay bound to genes during starvation, causing transcription to jumpstart as soon as nutrients become available again.

The molecular mechanisms underlying the “sweeper” activity of Sir3 are not known at this stage. Sir3 has both a histone and a DNA binding activity and while H4K16Ac, H3K4me and H3K79me have a negative effect on Sir3 binding affinity in the context of the SIR complex (Behrouzi et al. 2016), the presence of these modifications in transcribing genes does not exclude the possibility of a transient Sir3 contact. Whether Sir3 can displace transcription complexes on its own or whether it requires additional factors for its “sweeper” activity is obviously an open question that requires further study.

Our results from the competitive fitness test in Figure S10 are consistent with the idea that the “sweeper” activity of Sir3 improves transcription efficiency enough to give wt cells a competitive advantage over Sir3 mutants and increases their fitness in conditions when resources are becoming scarce. While the need for a mechanism that facilitates the elimination of non-functioning stalled transcription complexes from chromatin seems obvious for optimal growth, the necessity to control the speed at which transcription is reactivated after growth arrest may seem less evident. One could nevertheless imagine that a controlled and gradual restart of cellular processes is preferable to an uncoordinated “every gene for itself” approach. Since, according to our model, Sir3 removes only a portion of gene bound transcription complexes, genes that were highly transcribed in mid-log such as ribosomal protein genes or genes involved in cellular metabolism, will be reactivated faster than genes that were moderately transcribed because they would have kept more transcription complexes during starvation than moderately expressed genes. Consequently, the expression of moderately transcribed genes that are mostly not essential for restarting cell growth after starvation would be naturally delayed, which would allow highly transcribed genes involved in transcription, translation and metabolism to have sufficient quantities of newly available nutrients to jump-start cell growth. Radonjic et al. (Radonjic et al. 2005) have identified 769 transcripts that were induced more than 2 fold above their mid-log expression levels within 3 min after exit from stationary phase and that are mostly involved in transcription and protein synthesis. We find 41% and 25% of those transcripts in the group least and most affected by Sir3 mutations, respectively, which is consistent with the idea that the Sir3 “sweeper” activity disproportionally affects the reactivation of genes that are not immediately needed to restart growth.

Thanks to our three-pronged approach of measuring Sir3 exchange rates using ChIP-seq and Sir3 dependent changes in mRNA levels using spike-in RNA-seq, and mapping genome-wide transient Sir3 binding using Nanopore-MetID, we were able to uncover a new role for Sir3 in the regulation of gene expression and demonstrate that yeast heterochromatin is a highly dynamic structure that indirectly influences transcription on a global scale. Our model for the biological function of subtelomeric heterochromatin in budding yeast represents a paradigm shift from the generally internalized ideas on how chromatin architecture and chromatin binding proteins influence DNA based processes such as transcription. In contrast to the widely accepted view that chromatin structure directly influences the transcriptional activity of its underlying sequences i.e. that the principal function of heterochromatin is to repress transcription at the locus where it is assembled, we postulate that, unlike heterochromatin at mating type loci, subtelomeric heterochromatin in yeast does not act directly on the sequences where the SIR complex is assembled. It is instead a reservoir for Sir3, whose biological function is to modulate transcription throughout the chromosome far away from its “hub” at chromosome ends. We propose that the main biological function of subtelomeric heterochromatin is to compensate for the weak euchromatin binding affinity that Sir3 has on its own, by attracting and concentrating Sir3 at each chromosome end from where Sir3 can be “dispatched” throughout the chromosome, apparently through telomere looping, since 50% of Sir3 targets are also Sir2 and Sir4 targets (Figure 2). In other words, subtelomeric heterochromatin is needed to maximize the probability of stochastic non-specific contacts between Sir3 and most genes in the genome.

The idea that Sir3 mediated eviction of transcription complexes relies entirely on transient, low-affinity, non-specific binding of Sir3 to euchromatic genes challenges the prevalent view on how chromatin binding proteins perform their regulatory or enzymatic functions. The current “dogma” is that DNA based processes rely on the stable, high-affinity, locus-specific binding of proteins that either recruit other proteins that perform enzymatic reactions or perform them themselves. This idea has of course been consolidated by more than a decade worth of studies that use ChIP-seq to measure genome-wide binding of chromatin associated proteins. The fact that the undeniably powerful ChIP-seq based approaches repeatedly confirm the “dogma” in a wide range of organisms and systems is however not because stable locus-specific binding is the only functionally and biologically significant “way” for proteins to act but rather because it is the only “way” that ChIP-seq can reliably detect. Nanopore-MetID has now revealed over a thousand transient non-specific contacts between Sir3 and the yeast genome. We postulate that these transient Sir3 interactions with euchromatic genes that were previously either undetected or characterized as background noise or experimental artefacts of ChIP assays, are in fact important for optimal genome-wide transcription and cellular fitness (**Figures 7, 8 and S10**).

The future challenge of chromatin biology research will be to expand our understanding of the dynamics of chromatin structure maintenance and renewal that goes beyond the information typically afforded by classical single time point ChIP-seq experiments. In light of our study, we now need to start including transient/unstable associations between proteins and DNA into our catalogue of protein-DNA interactions with a biological function, if we want to fully understand how chromatin shapes transcription programs and cellular phenotypes in response to environmental or developmental cues.

## Materials and Methods

Detailed Experimental Procedures are listed in the SI Appendix.

### Yeast Strains

Genotypes and strain construction are described in the SI Appendix.

### Sir3 Nanopore MetID cell cultures and genomic DNA preparation for nanopore sequencing

For mid-log cultures, cell were grown overnight until they reached saturation (OD>1). For the release from starvation time course (Figure 3A), cells were grown for 72hrs, an aliquot was taken for the starvation time point. Cells were then pelleted and re-suspended in fresh YPD and aliquots were then taken at indicated times and pellets were kept on ice until spheroplasting.

Pelleted cells were counted and divided into aliquots of 10^9^ cells, the amount needed for one sequencing run. Pellets were then spheroplasted with Zymolyase and genomic DNA was extracted with the MagAttract HMW DNA Kit (Qiagen) according to the manufacturer’s protocol. Nanopre sequencing libraries were prepared with the Rapid Barcoding Kit (Oxford Nanopore), according to the manufacturer protocol. The library mix was loaded on the R9.4.1 Flow cell (Oxford Nanopore) and sequenced with the Minion device (Oxford Nanopre).

### Cell culture for release from starvation

Cells were kept at 30°C in rich media for 48 hrs after a 10 fold dilution with fresh YPD media of a saturated overnight culture, until carbon source depletion caused growth arrest, as described in the SI Appendix. For the tag switch experiments estradiol was added and cells were incubated for another 16 hrs before release. For the experiments without tag switch cells were released into fresh media after 72 hrs and fixed with formaldehyde at indicated time points.

### Microscopy and image analysis

Cells were injected under a 0.8% agarose/growth media layer in 8-well glass bottom microscopy plates (BioValley) and visualized with a Nikon Ti2 Eclipse widefield inverted microscope in the triple channel LED DIC mode.

### Chromatin Sonication and ChIP

Cell walls were mechanically disrupted and the entire cell lysate was sonicated. ChIPs were done as described in the SI Appendix using anti-HA (Abcam, ab9110 (lot# GR3245707-3) and polyclonal anti-T7 (Bethyl A190-117A (lot# A190-117A-7) (Figure 5) or monoclonal anti-T7 (Cell Signaling Technology, DSE1X (lot#1)) (Supplementary Figure S5). Purified DNA was treated with RNAse A and used for NGS library construction.

### RNA isolation and RNA-seq library construction

Cells were flash frozen in liquid N_2_ and total RNA was isolated from frozen cell pellets with Trizol and treated with DNAseI. RNA samples were then used for NGS library preparation with the Illumina TruSeq Stranded mRNA kit or the Illumina Stranded mRNA Prep Ligation kit according to the manufacturer’s protocol. Libraries were sequenced on the Illumina NextSeq550 (2×75bp) (Plateforme Transcriptome, IRMB, Montpellier, France) or NovaSeq 6000 (2×75bp) (Illumina) at the CNAG, Barcelona.

### NGS Input and ChIP library construction and Illumina sequencing

DNA fragments were blunt ended and phosphorylated with the Epicentre End-it-Repair kit. Adenosine nucleotide overhangs were added using Epicentre exo-Klenow. Illumina Genome sequencing adaptors with in line barcodes were then ligated using the Epicentre Fast-Link ligation kit. Ligated fragments were amplified using the Phusion enzyme Library pools were sequenced on the HiSeq 2000 (2×75bp) (Illumina) at the CNAG, Barcelona, Spain or the NextSeq 550 (2×75bp) (Plateforme Transcriptome, IRMB, Montpellier, France).

### ChIP-seq and RNA-seq data analysis

Sequences were aligned to the *S. Cerevisiae* genome with BLAT (Kent Informatics, http://hgdownload.soe.ucsc.edu/admin/). Read count distribution was determined in 1bp windows and then normalized to 1 by dividing each base pair count with the genome-wide average base-pair count.

RNA-seq normalized read densities for each gene were aligned by the transcription start site and divided into sense and antisense transcripts. The median read density was determined for each transcript as above and normalized to the genome average read count of the *S.pombe* spike-in, as described in the SI Appendix and Figure S8. Intron regions were excluded from the calculation.

### Sir3 Nanopore-MetID nanopore sequencing data analysis

Basecalling, demultiplexing and aligning to the *S.cerevisiae* genome was performed with the guppy software from Oxford Nanopre. Fastq files and modification probability tables were extracted from basecalled fast5 files using the ont-fast5-api package. Adenines with a methylation probability higher or equal to 0.95 were considered as positive signals.

## Data availability

NGS and ONT (Oxford Nanopore Technology) datasets were submitted to the NCBI GEO database. GEO accession numbers for ChIP-seq and RNA-seq datasets are

## Author Contributions

PB and AH performed experiments in Figures 4 and 7, PB performed the experiments in Figure S6 and helped with cell culture for Figure 3A. AC and LD performed the experiments from Figures 1, 2, S1 and S2 and AC performed experiments in Figure 3 (replicate 2). HG performed experiments in Figures 3B-C (replicate 1), 5, 6, S3 and S5. LTN performed the experiment with the standalone NLSEcoG2 control for experiments in Figures 1, 2 and S2. PV performed the microscopy in Fig 4E. MRL, PB, AH, HG and AC designed the experiments, MRL analyzed the data, wrote the manuscript and performed the experiment in Figure S10.

## Acknowledgments

We thank Fred van Leeuwen, Kevin Struhl and Laure Crabbe for yeast strains and plasmids. Thank you to Marta Gut and Julie Blanc from CNAG (Barcelona, Spain) and Véronique Pantesco (IRMB, Montpellier, France) for Illumina sequencing services. We thank Virginie Georget and Leslie Bancel-Vallée (MRI, Biocampus, Montpellier) for their assistance with the microscope set-up. Thanks to Sylvie Fromont and Christelle Anguille (MGC platform, IGMM-CRBM, Montpellier) for the LabChip experiments. Thank you to Fabrice Caudron for critical reading of the manuscript. Two previous versions of this article are part of the doctoral theses “Heterochromatin dynamics upon release from growth arrest in budding yeast” by Hrvoje Galić and “Functional analysis of heterochromatin dynamics after exit from growth arrest in budding yeast” by Ana Hrgovčić. This work was supported by the ERC-Consolidator (NChIP 647618) (MRL), Canceropole GSO_Emergence 2020 (NanoRep-MetAID; n°2020-E11; MRL) and the CNRS, MITI “Evènements rares 2022-2023” (MRL) grants.

## References

Ai W, Bertram PG, Tsang CK, Chan TF, Zheng XF. 2002. Regulation of subtelomeric silencing during stress response. Mol Cell 10: 1295–1305.

Allen C, Buttner S, Aragon AD, Thomas JA, Meirelles O, Jaetao JE, Benn D, Ruby SW, Veenhuis M, Madeo F et al. 2006. Isolation of quiescent and nonquiescent cells from yeast stationary-phase cultures. The Journal of cell biology 174: 89–100.

Auboiron M, Vasseur P, Tonazzini S, Fall A, Castro FR, Sučec I, El Koulali K, Urbach S, Radman-Livaja M. 2021. TrIPP-a method for Tracking the Inheritance Patterns of Proteins in living cells-reveals retention of Tup1p, Fpr4p and Rpd3L in the mother cell. iScience: 102075.

Batté A, Brocas C, Bordelet H, Hocher A, Ruault M, Adjiri A, Taddei A, Dubrana K. 2017. Recombination at subtelomeres is regulated by physical distance, double-strand break resection and chromatin status. EMBO J 36: 2609–2625.

Behrouzi R, Lu C, Currie MA, Jih G, Iglesias N, Moazed D. 2016. Heterochromatin assembly by interrupted Sir3 bridges across neighboring nucleosomes. Elife 5.

Brothers M, Rine J. 2022. Distinguishing between recruitment and spread of silent chromatin structures in. Elife 11.

Cheng TH, Gartenberg MR. 2000. Yeast heterochromatin is a dynamic structure that requires silencers continuously. Genes Dev 14: 452–463.

Deneke C, Lipowsky R, Valleriani A. 2013. Complex degradation processes lead to non-exponential decay patterns and age-dependent decay rates of messenger RNA. PLoS One 8: e55442.

DuBois ML, Haimberger ZW, McIntosh MW, Gottschling DE. 2002. A quantitative assay for telomere protection in Saccharomyces cerevisiae. Genetics 161: 995–1013.

Ellahi A, Thurtle DM, Rine J. 2015. The Chromatin and Transcriptional Landscape of Native Saccharomyces cerevisiae Telomeres and Subtelomeric Domains. Genetics 200: 505–521.

Fraser JA, Heitman J. 2003. Fungal mating-type loci. Curr Biol 13: R792–795.

Gartenberg MR, Smith JS. 2016. The Nuts and Bolts of Transcriptionally Silent Chromatin in Saccharomyces cerevisiae. Genetics 203: 1563–1599.

Gasch AP, Spellman PT, Kao CM, Carmel-Harel O, Eisen MB, Storz G, Botstein D, Brown PO. 2000. Genomic expression programs in the response of yeast cells to environmental changes. Mol Biol Cell 11: 4241–4257.

Gottlieb S, Esposito RE. 1989. A new role for a yeast transcriptional silencer gene, SIR2, in regulation of recombination in ribosomal DNA. Cell 56: 771-776.

Grunstein M, Gasser SM. 2013. Epigenetics in Saccharomyces cerevisiae. Cold Spring Harb Perspect Biol 5.

Guidi M, Ruault M, Marbouty M, Loiodice I, Cournac A, Billaudeau C, Hocher A, Mozziconacci J, Koszul R, Taddei A. 2015. Spatial reorganization of telomeres in long-lived quiescent cells. Genome Biol 16: 206.

Hecht A, Strahl-Bolsinger S, Grunstein M. 1996. Spreading of transcriptional repressor SIR3 from telomeric heterochromatin. Nature 383: 92–96.

Hoggard TA, Chang F, Perry KR, Subramanian S, Kenworthy J, Chueng J, Shor E, Hyland EM, Boeke JD, Weinreich M et al. 2018. Yeast heterochromatin regulators Sir2 and Sir3 act directly at euchromatic DNA replication origins. PLoS Genet 14: e1007418.

Martinez MJ, Roy S, Archuletta AB, Wentzell PD, Anna-Arriola SS, Rodriguez AL, Aragon AD, Quiñones GA, Allen C, Werner-Washburne M. 2004. Genomic analysis of stationary-phase and exit in Saccharomyces cerevisiae: gene expression and identification of novel essential genes. Mol Biol Cell 15: 5295–5305.

McIntyre ABR, Alexander N, Grigorev K, Bezdan D, Sichtig H, Chiu CY, Mason CE. 2019. Single-molecule sequencing detection of N6-methyladenine in microbial reference materials. Nat Commun 10: 579.

McKnight JN, Boerma JW, Breeden LL, Tsukiyama T. 2015. Global Promoter Targeting of a Conserved Lysine Deacetylase for Transcriptional Shutoff during Quiescence Entry. Mol Cell 59: 732–743.

Miller C, Schwalb B, Maier K, Schulz D, Dumcke S, Zacher B, Mayer A, Sydow J, Marcinowski L, Dolken L et al. 2011. Dynamic transcriptome analysis measures rates of mRNA synthesis and decay in yeast. Mol Syst Biol 7: 458.

Müller CA, Boemo MA, Spingardi P, Kessler BM, Kriaucionis S, Simpson JT, Nieduszynski CA. 2019. Capturing the dynamics of genome replication on individual ultra-long nanopore sequence reads. Nat Methods 16: 429–436.

Radman-Livaja M, Ruben G, Weiner A, Friedman N, Kamakaka R, Rando OJ. 2011. Dynamics of Sir3 spreading in budding yeast: secondary recruitment sites and euchromatic localization. Embo J 30: 1012–1026.

Radonjic M, Andrau JC, Lijnzaad P, Kemmeren P, Kockelkorn TT, van Leenen D, van Berkum NL, Holstege FC. 2005. Genome-wide analyses reveal RNA polymerase II located upstream of genes poised for rapid response upon S. cerevisiae stationary phase exit. Mol Cell 18: 171–183.

Redolfi J, Zhan Y, Valdes-Quezada C, Kryzhanovska M, Guerreiro I, Iesmantavicius V, Pollex T, Grand RS, Mulugeta E, Kind J et al. 2019. DamC reveals principles of chromatin folding in vivo without crosslinking and ligation. Nat Struct Mol Biol 26: 471–480.

Renauld H, Aparicio OM, Zierath PD, Billington BL, Chhablani SK, Gottschling DE. 1993. Silent domains are assembled continuously from the telomere and are defined by promoter distance and strength, and by SIR3 dosage. Genes Dev 7: 1133–1145.

Ruault M, Scolari VF, Lazar-Stefanita L, Hocher A, Loïodice I, Koszul R, Taddei A. 2021. Sir3 mediates long-range chromosome interactions in budding yeast. Genome Res.

Sedat J, McDonald A, Kasler H, Verdin E, Cang H, Arigovindan M, Murre C, Elbaum M. 2022. A proposed unified mitotic chromosome architecture. Proc Natl Acad Sci U S A 119: e2119107119.

Szczesnik T, Ho JWK, Sherwood R. 2019. Dam mutants provide improved sensitivity and spatial resolution for profiling transcription factor binding. Epigenetics Chromatin 12: 36.

Teytelman L, Thurtle DM, Rine J, van Oudenaarden A. 2013. Highly expressed loci are vulnerable to misleading ChIP localization of multiple unrelated proteins. Proceedings of the National Academy of Sciences of the United States of America 110: 18602–18607.

Thurtle DM, Rine J. 2014. The molecular topography of silenced chromatin in Saccharomyces cerevisiae. Genes Dev 28: 245–258.

van Steensel B, Henikoff S. 2000. Identification of in vivo DNA targets of chromatin proteins using tethered dam methyltransferase. Nat Biotechnol 18: 424–428.

Verzijlbergen KF, Menendez-Benito V, van Welsem T, van Deventer SJ, Lindstrom DL, Ovaa H, Neefjes J, Gottschling DE, van Leeuwen F. 2010. Recombination-induced tag exchange to track old and new proteins. Proc Natl Acad Sci U S A 107: 64–68.

