## Supplemental Figures for "The budding yeast heterochromatic protein Sir3 modulates genome-wide gene expression through transient direct contacts with euchromatin"

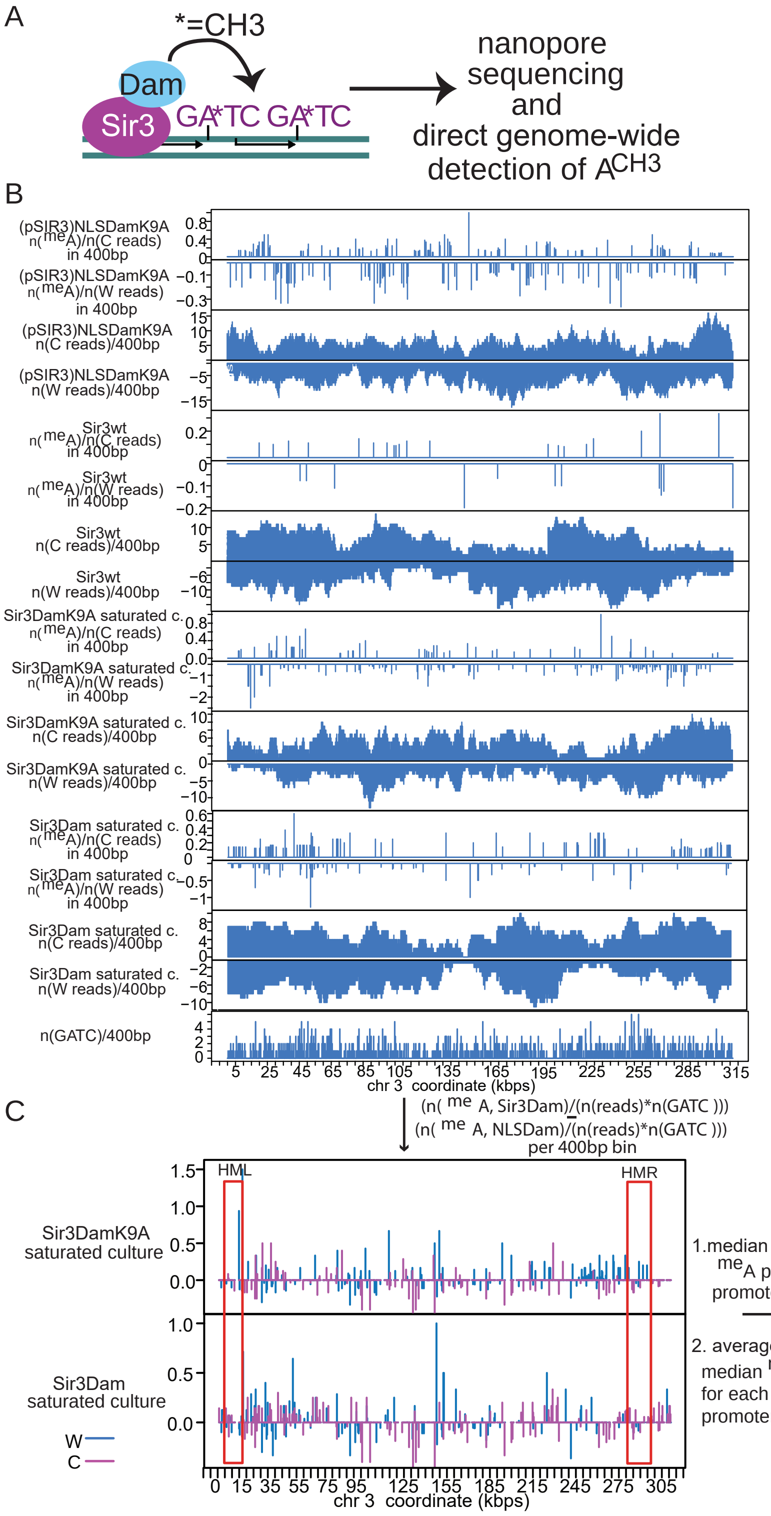


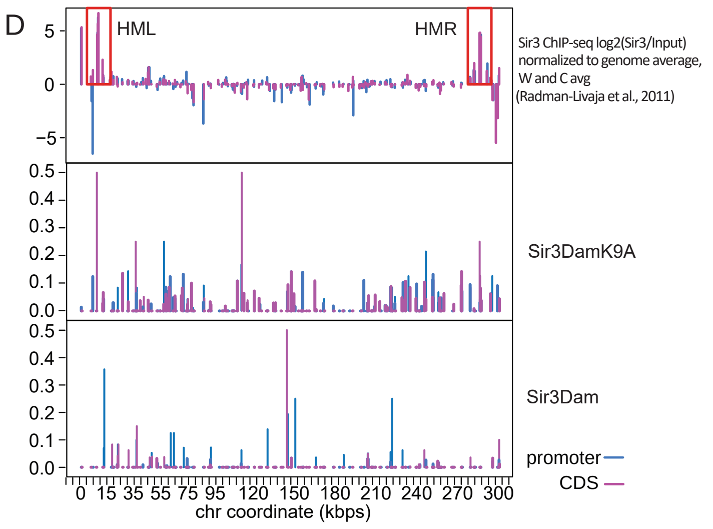


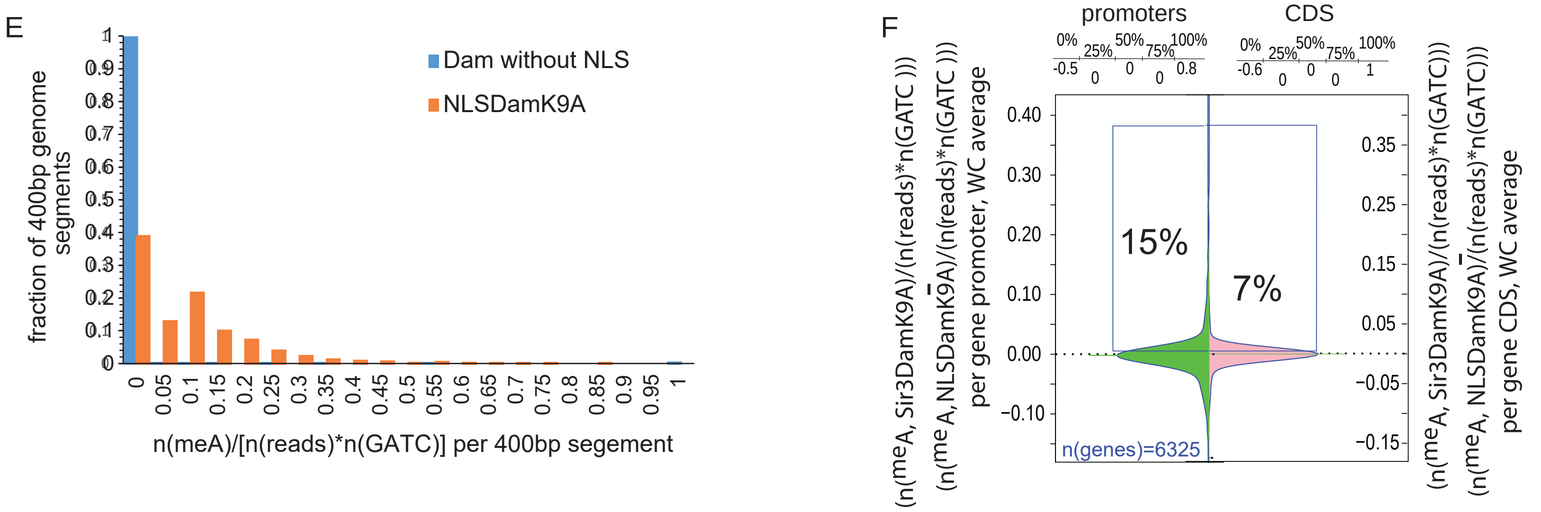


**Figure S1:Analysis of Sir3Dam nanopore sequencing data. A.** Diagram of Dam Adenine methylation within GATC sequences in proximity of Sir3 contact sites. **B.** Distribution of read or feature count number (as indicated on the left of the plot) per 400bp bins on chromosome 3. DNA from Sir3Dam, Sir3DamK9A, Sir3wt (untagged Sir3) or (pSIR3)NLSDamK9A ( «standalone» DamK9A with an NLS sequence under the control of the SIR3 promoter) strains was extracted from saturated cultures at the end of the exponential growth phase. W-Watson strand, C-Crick strand. Read counts from the W strand are shown as negative values for easier visualization. **C.** Normalized meA counts (n(meA)/n(reads)) from Sir3Dam, Sir3DamK9A and (pSIR3)NLSDamK9A from B were divided by the GATC count (n(GATC)) in every 400bp bin (if there are no GATCs in the bin the signal was considered as NA) and the normalized meA signal from the (pSIR3)NLSDamK9A strain was subtracted from the meA Sir3Dam and Sir3DamK9A signals. Signals from the W(atson) (blue) and the C(rick) (magenta) strands are shown in the same panel . **D.** The median meA signal of signals from C were first determined for each gene promoter and each gene CDS and then these signals were averaged between the Watson and the Crick strands if neither was marked as NA, otherwise only the non NA value was kept. The WC average Sir3 enrichment per gene promoter and CDS from a wt Sir3 ChIP-seq dataset from Radman-Livaja et al. (2011) is shown in the top panel for comparison. **E.** Comparison of GATC methylation probabilities between a strain that contains a stand alone Dam gene without the NLS sequence and the NLSDamK9A strain. Both constructs are under the control of the Sir3 promoter. We first selected 400bp segements from the genome whose sequences were present in both datasets, than we determined the distribution of GATC methylation probabilities in the set of selected 400bp segements from the Dam_without_NLS (blue) and NLSDamK9A (orange) strains. **F.** Density plots of normalized Sir3DamK9A signals at gene promoters and CDS from saturated cultures from D. 15% and 7% of gene promoters and CDS, respectively, have a normalized Sir3DamK9A signal >0

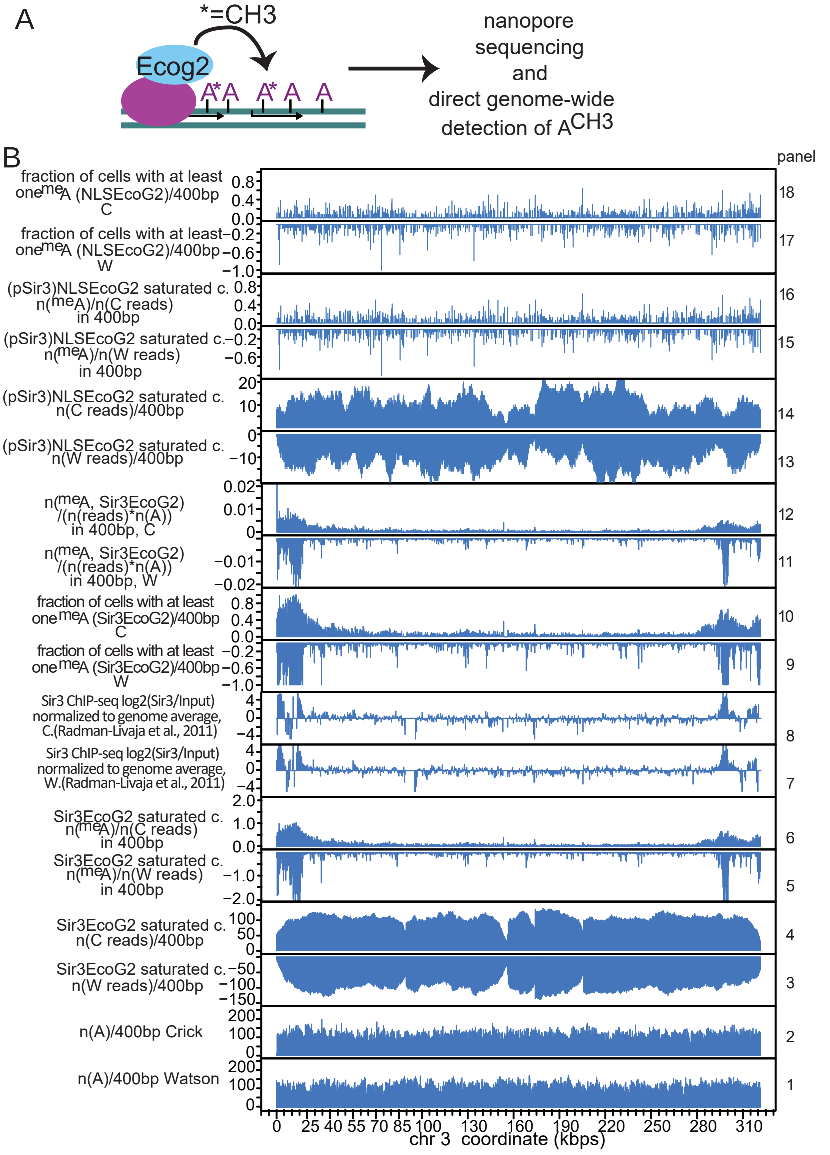


**Figure S2:Analysis of Sir3EcoG2 nanopore sequen-cing data. A.** Diagram of EcoG2 Adenine methylation in proximity of Sir3 binding sites. **B**. Distribution of read or feature count number (as indicated on the left of the plot) per 400bp bins on chromosome 3. DNA from the Sir3EcoG2 and (pSir3)NLSEcoG2 («stand-alone» EcoG2 with an NLS under the control of the Sir3 promoter) strains was extracted from a saturated culture (sat.c.) at the end of the exponential growth phase. Panels 11-12: Normalized meA counts (n(meA)/n(reads), panels 5 - 6) were divided by the Adenine count (n(A)) in every 400bp bin (panels 1 and 2). Panels 9-10 and 17-18: For the fraction of cells with at least one meA/400bp, we use normalized meA counts (panels 5 , 6 and 15,16) if the counts are ≤1 or convert them to 1 if they are >1. The Sir3 enrichment from the wt Sir3 ChIP-seq dataset from Radman-Livaja et al. (2011) for the Crick and the Watson strand are shown in panels 7 and 8, respectively.W-Watson strand, C-Crick strand. Read counts from the W strand are shown as negative values for easier visualization.**C.** WC averages of normalized W(atson) and C(rick) signals from B. subtracted from the meA signal of the strain with (pSir3)NLS-EcoG2. For the fraction of cells with at least one meA/400bp, we used normalized meA counts that were (panels 5 , 6 in B) subtracted from normalized meA counts from NLSEcoG2 (panels 15,16 in B); if the sum of the W and C counts is ≤1 and >0, we use the avg of W and C counts; if the sum of the W and C counts is >1, we use 1 (meaning that there is at least 1 meA on the W or the C strand in that 400bp segement in all cells); if the sum of the W and C counts is ≤ 0, we use 0 (meaning that this 400bp segment is not specifically methylated by Sir3EcoG2 in any cell). **D.** median meA signals from C per gene promoter or CDS **E.** Differences in Sir3EcoG2 mediated Adenine methylation on W and C strands . The plot shows the genomic meA density distribution (n(meA)/ (n(reads)* n(A)) **F.** Density plots of normalized Sir3EcoG2 signals at gene promoters and CDS from D. The average population density of meA in promoters and CDS that are specifically targeted by Sir3EcoG2 (33% of all promoters and 24% of all CDS) is low: 0.07% and 0.06%, respectively (left panel). Targeted promoters and CDS are on average specifically contacted by Sir3EcoG2 in 8% and 6% of the cell population, respectively (right panel).

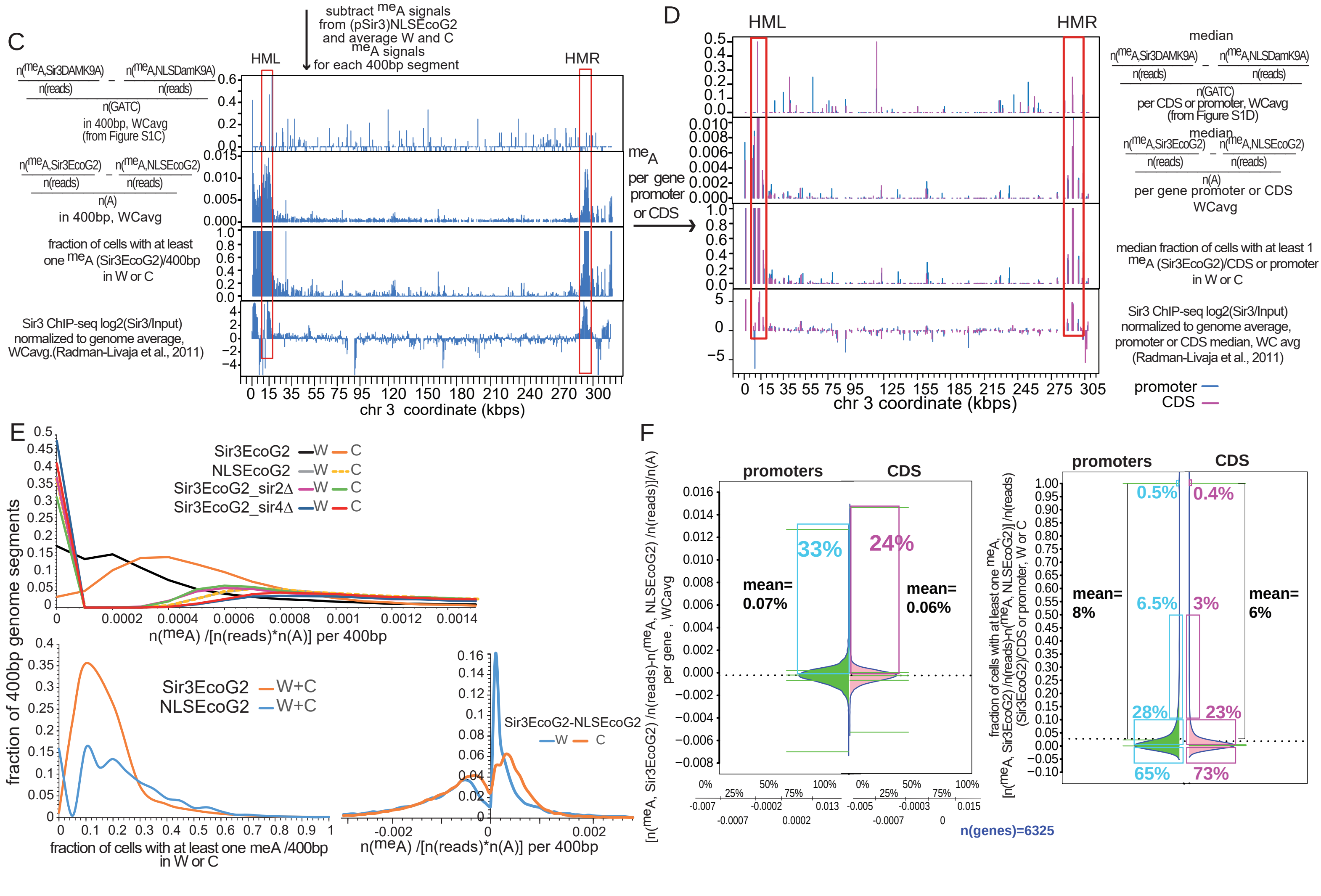

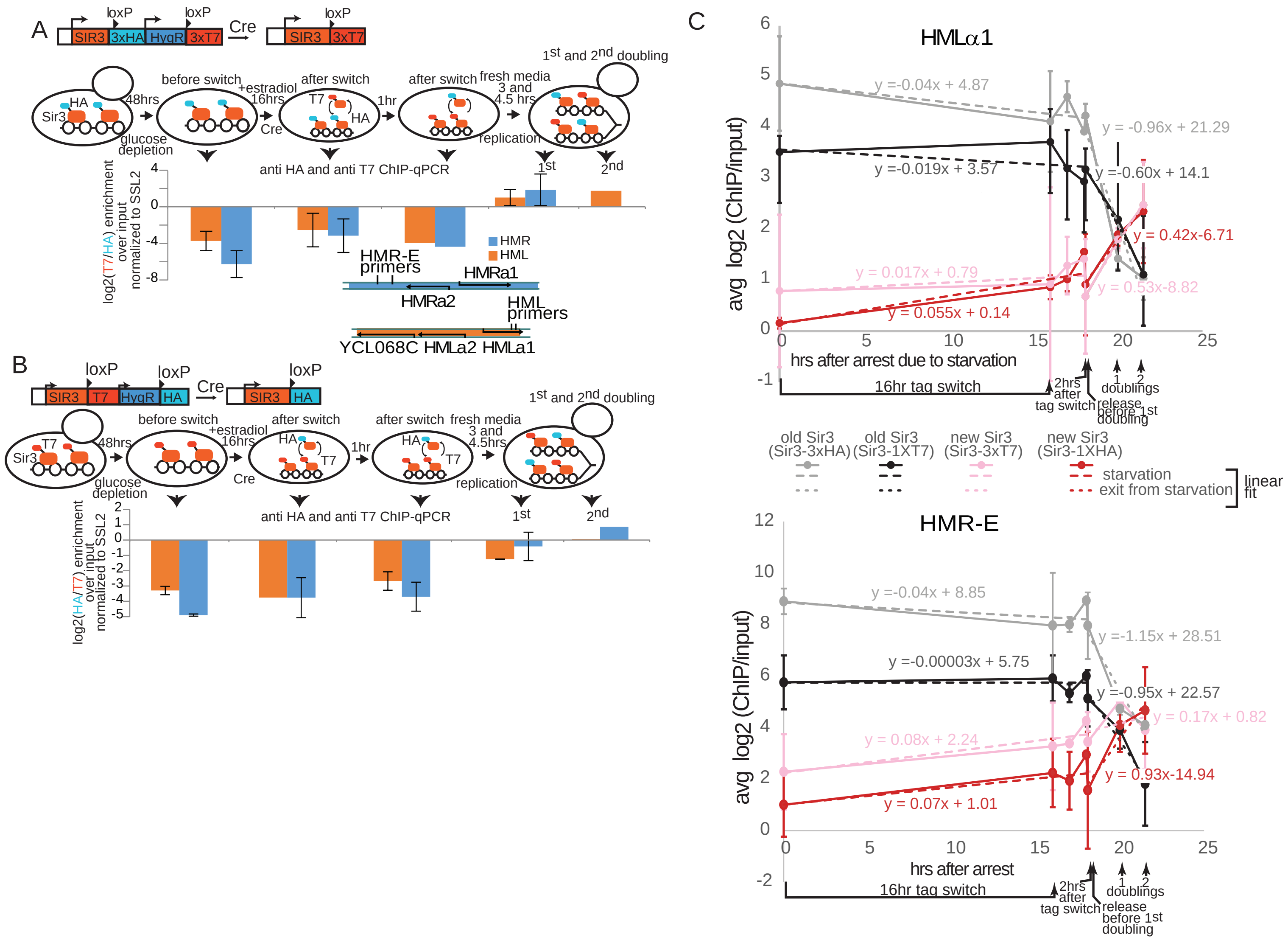

**Figure S3: Cell growth dependent and independent Sir3 turnover at silent mating type loci. A.-B.** Sir3 tag switch ChIP qPCR experiments for 3xHA to 3xT7 (A) and 1xT7 to 1xHA tag switches (B). Cells are arrested by glucose depletion before the tag switch. The switch is induced with estradiol addition to arrested cells (98% of cells have switched after a 16hr incubation). Cells are then released from arrest with addition of fresh media and allowed to grow for one or two doublings. Cell aliquots were fixed with 1% formaldehyde for 20min at times indicated in the diagrams and anti-HA and anti-T7 ChIPs were performed on sonicated chromatin. Bar graphs show the ratios of new (T7 in A or HA in B) over old (HA in A or T7 in B) Sir3 enrichment at HMR (blue) and HML(orange) silent mating type loci measured by qPCR (Cts were normalized to the SSL2 locus (where Sir3 does not bind) for ChIPs and inputs, separately, and the ChIP ΔCts were then normalized to input ΔCts). The location of qPCR primers are shown in the diagram below the bar graph in A. The bar graphs show the average of two biological replicates (error bars are deviations from the average). **C.** The scatter plots show the apparent ON (change in new Sir3p (ChIP/Input) log2 ratio in $hr^{-1}$) and OFF (change in old Sir3p (ChIP/Input) log2 ratio in $hr^{-1}$) Sir3p rates at HML (top) and HMR (bottom), calculated from the slopes of linear fits before and after release, as shown. Error bars are standard deviations from the average of two biological replicates.

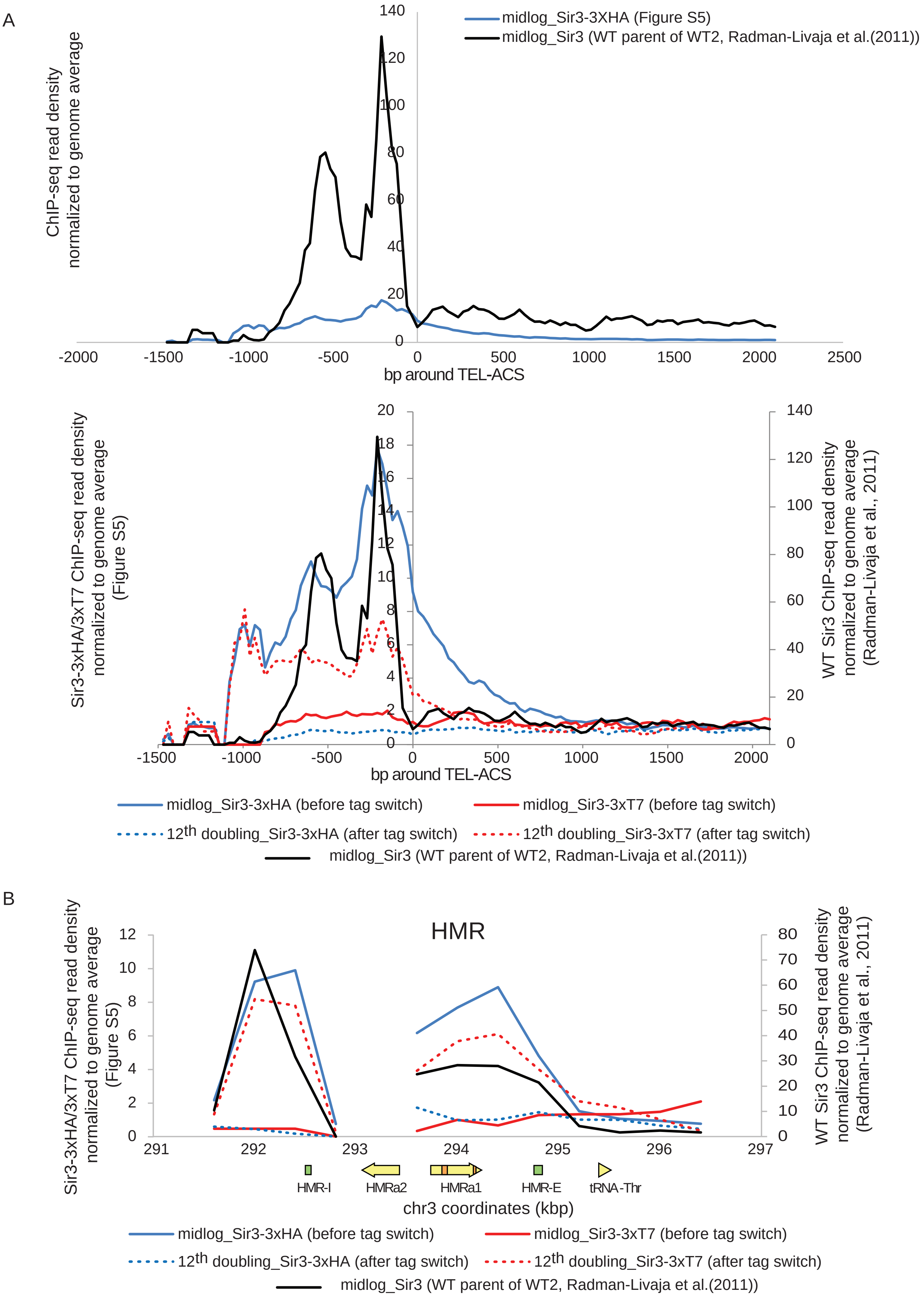


Figure S4: Comparison of untagged and 3xHA/3xT7 tagged Sir3 ChIP-seq in subtelomeric regions (A) and at the HMR silent mating type locus (B)

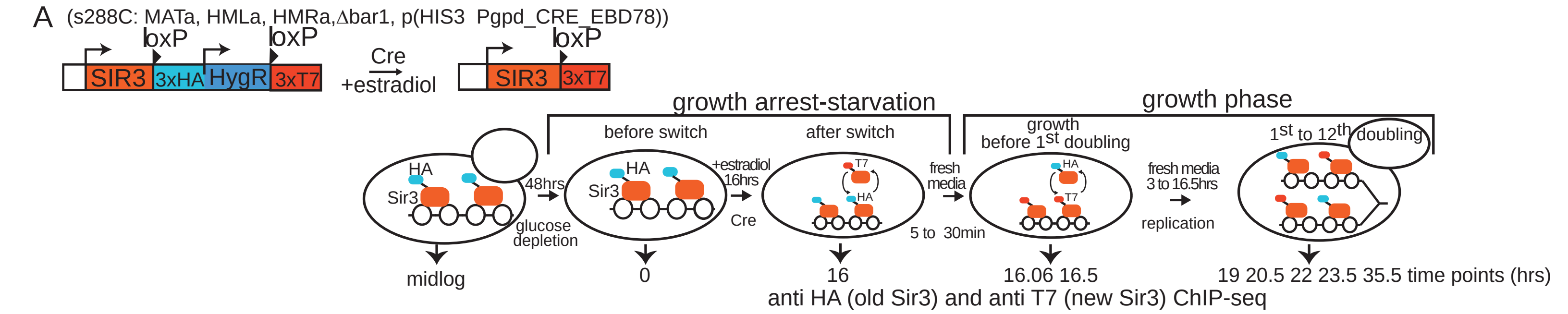


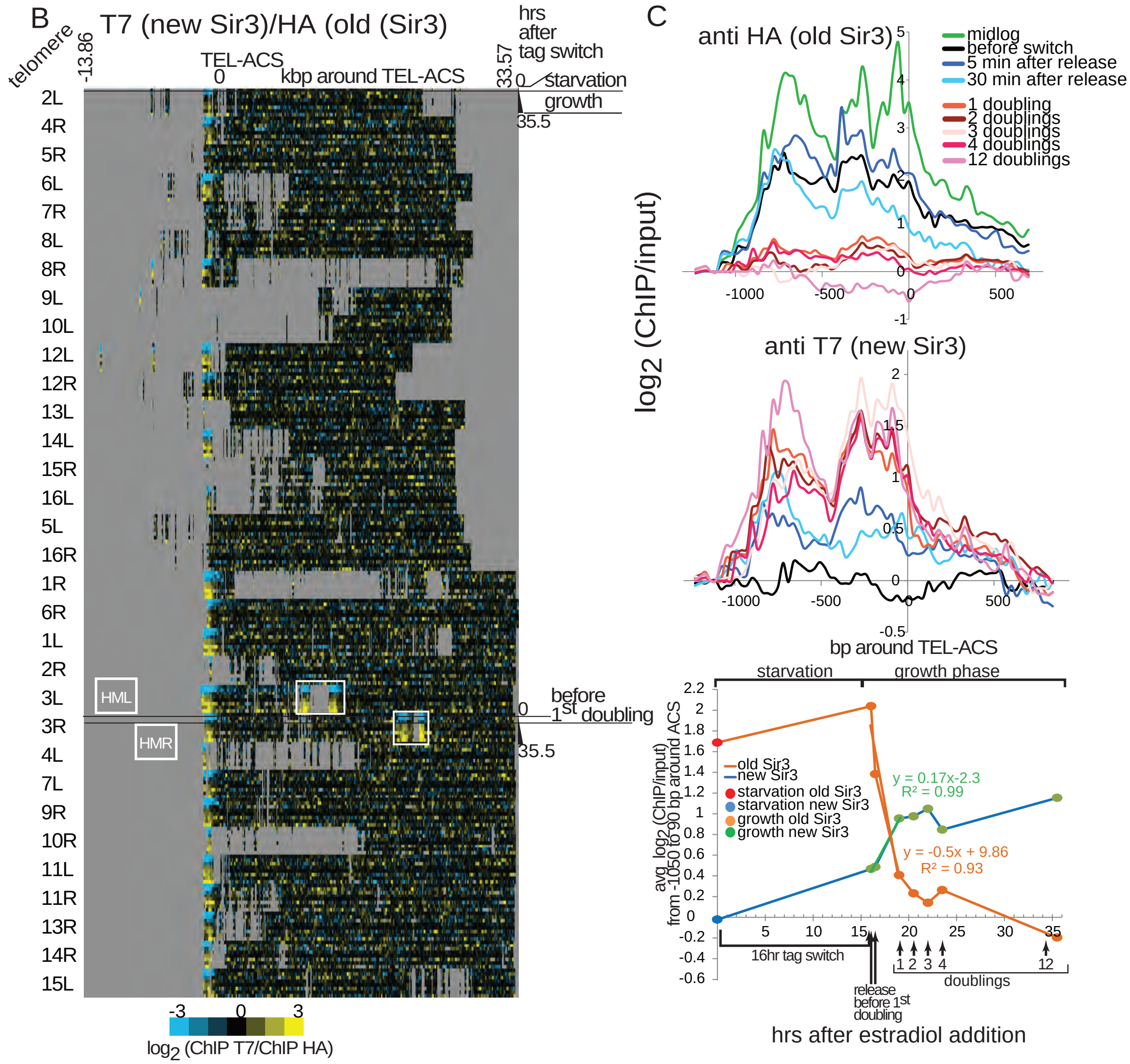


**Figure S5: Replicate of experiment from Fig. 5. A.** Diagram of the Sir3 tag switch construct (top left). Bottom: Experiment outline. Cells were arrested by glucose depletion before the tag switch, which is induced with estradiol addition to arrested cells (recombination efficiency:97.5%). Cells were then released from arrest with addition of fresh media and allowed to grow for one to twelve doublings (monitored by OD measurements). Cell aliquots were fixed with 1% formaldehyde for 20min at times indicated below the diagrams and anti-HA and anti-T7 (monoclonal) ChIPs were performed on sonicated chromatin. **B**. Heat map of new Sir3 (T7 tag) enrichment over old Sir3 (HA tag) during and after exit from starvation, at all yeast telomeres (30kbp from chromosomes ends). Time points are aligned by the ARS Consensus Sequence (TEL-ACS) located in telomeric silencer regions. Silent mating type loci HML and HMR, on 3L and 3R, respectively, are framed with a white rectangle. Sir3 is enriched in a small 1kb region upstream of the TEL-ACS at all telomeres. Repetitive and unmapped regions are shown in grey. The HMLa reads have been eliminated as repetitive sequences during alignment to the reference genome which is MATa. **C**. Old (top) and new (middle) Sir3 enrichment around TEL-ACS averaged for all 32 telomeres at indicated time points during starvation and the renewed growth phase. The bottom panel shows average enrichment around the TEL-ACS for old and new Sir3 over time with Sir3p ON (green) and OFF (orange) rates during the first cycle after release calculated from slopes of linear fits of time points 5min, 30min and 1 doubling after release as in Figure 4. **D-E.** Old (top left) and new (bottom left) Sir3 enrichment at HML (D) and HMR (E) at indicated time points (same color code for D and E) during starvation and the growth phase after release from starvation. The right panel shows average enrichment over the entire silent mating type locus for old and new Sir3p over time.

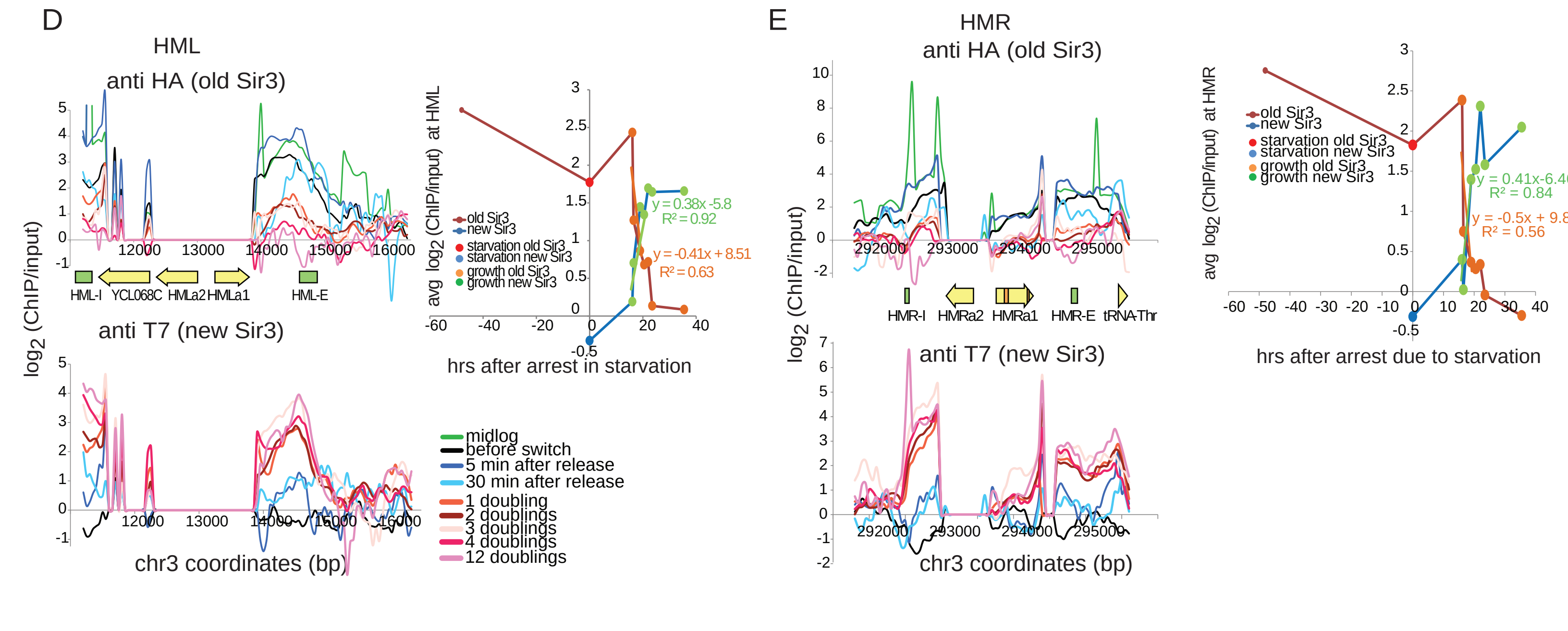

A

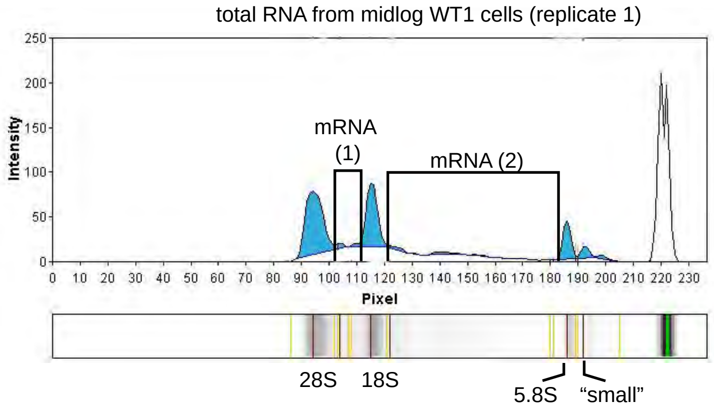


| | 28S | mRNA (1) | 18S | mRNA (2) | 5.8S | small | total RNA |
|---|---|---|---|---|---|---|---|
| pixel intensity | 538 | 14 | 348 | 51 | 153 | 60 | 1164 |

$$\text{mRNA fraction}=\frac{\text{mRNA(1)+mRNA(2)}}{\text{total RNA}}=0.056$$

$$\text{rRNA fraction}=\frac{\text{28S+18S+5.8S}}{\text{total RNA}}=0.89$$

total RNA=28S+18S+5.8S+mRNA(1)+mRNA(2)+"small"

B

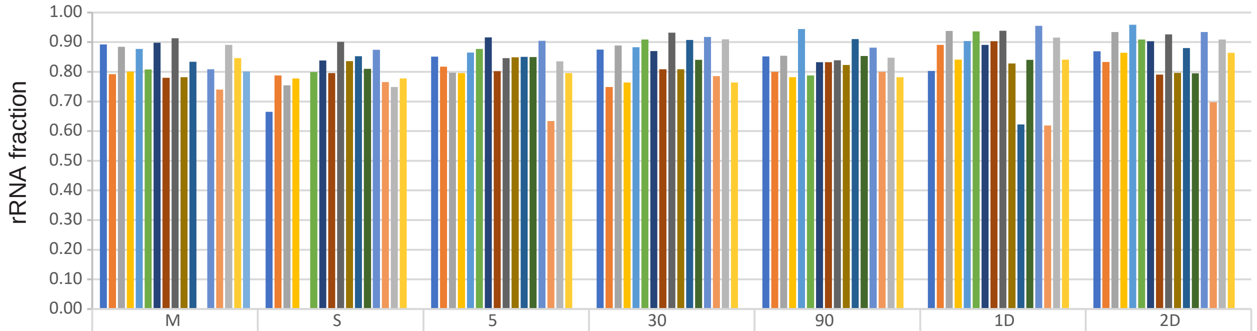


| | M | S | 5 | 30 | 90 | 1D | 2D |
|---|---|---|---|---|---|---|---|
| WT1 1 | 0.89 | 0.67 | 0.85 | 0.88 | 0.85 | 0.80 | 0.87 |
| WT1 2 | 0.79 | 0.79 | 0.82 | 0.75 | 0.80 | 0.89 | 0.83 |
| WT2 1 | 0.88 | 0.75 | 0.80 | 0.89 | 0.85 | 0.94 | 0.93 |
| WT2 2 | 0.80 | 0.78 | 0.80 | 0.76 | 0.78 | 0.84 | 0.86 |
| oeSir3 1 | 0.88 | ND | 0.87 | 0.88 | 0.94 | 0.90 | 0.96 |
| oeSir3 2 | 0.81 | 0.80 | 0.88 | 0.91 | 0.79 | 0.94 | 0.91 |
| Sir3_3xT7 1 | 0.90 | 0.84 | 0.92 | 0.87 | 0.83 | 0.89 | 0.90 |
| Sir3_3xT7 2 | 0.78 | 0.80 | 0.80 | 0.81 | 0.83 | 0.90 | 0.79 |
| Sir3_3xHA 1 | 0.91 | 0.90 | 0.85 | 0.93 | 0.84 | 0.94 | 0.93 |
| Sir3_3xHA 2 | 0.78 | 0.84 | 0.85 | 0.81 | 0.82 | 0.83 | 0.80 |
| Sir4Δ 1 | 0.83 | 0.85 | 0.85 | 0.91 | 0.91 | 0.62 | 0.88 |
| Sir4Δ 2 | ND | 0.81 | 0.85 | 0.84 | 0.85 | 0.84 | 0.80 |
| SIR2Δ 1 | 0.81 | 0.87 | 0.90 | 0.92 | 0.88 | 0.95 | 0.93 |
| SIR2Δ 2 | 0.74 | 0.77 | 0.63 | 0.79 | 0.80 | 0.62 | 0.70 |
| SIR3Δ 1 | 0.89 | 0.75 | 0.84 | 0.91 | 0.85 | 0.92 | 0.91 |
| SIR3Δ 2 | 0.85 | 0.78 | 0.80 | 0.76 | 0.78 | 0.84 | 0.86 |
| pombe | 0.80 | | | | | | |

time points

M: Midlog
S: starvation

5: 5min after release
30: 30min after release
90: 90min after release

1D: 1st division after release
2D: 2nd division after release

$$\text{rRNA corr.coeff}=\frac{\text{rRNA(S.cer.)}}{\text{rRNA(S.pombe)}}$$

| | M | S | 5 | 30 | 90 | 1D | 2D |
|---|---|---|---|---|---|---|---|
| WT1 1 | 1.12 | 0.83 | 1.06 | 1.09 | 1.07 | 1.00 | 1.09 |
| WT1 2 | 0.99 | 0.98 | 1.02 | 0.94 | 1.00 | 1.11 | 1.04 |
| WT2 1 | 1.11 | 0.94 | 1.00 | 1.11 | 1.07 | 1.17 | 1.17 |
| WT2 2 | 1.00 | 0.99 | 1.05 | 1.06 | 1.04 | 1.10 | 1.10 |
| oeSir3 1 | 1.07 | ND | 1.08 | 1.10 | 1.18 | 1.13 | 1.20 |
| oeSir3 2 | 1.02 | 1.00 | 1.10 | 1.14 | 0.98 | 1.17 | 1.14 |
| Sir3_3xT7 1 | 1.12 | 1.05 | 1.15 | 1.09 | 1.04 | 1.11 | 1.13 |
| Sir3_3xT7 2 | 0.97 | 1.00 | 1.00 | 1.01 | 1.04 | 1.13 | 0.99 |
| Sir3_3xHA 1 | 1.14 | 1.13 | 1.06 | 1.17 | 1.05 | 1.17 | 1.16 |
| Sir3_3xHA 2 | 0.98 | 1.04 | 1.06 | 1.01 | 1.03 | 1.03 | 1.00 |
| Sir4Δ 1 | 1.05 | 1.07 | 1.06 | 1.13 | 1.14 | 0.78 | 1.10 |
| Sir4Δ 2 | ND | 1.01 | 1.06 | 1.05 | 1.07 | 1.05 | 0.99 |
| SIR2Δ 1 | 1.01 | 1.09 | 1.13 | 1.15 | 1.10 | 1.19 | 1.17 |
| SIR2Δ 2 | 0.92 | 0.96 | 0.79 | 0.98 | 1.00 | 0.77 | 0.87 |
| SIR3Δ 1 | 1.11 | 0.94 | 1.04 | 1.14 | 1.06 | 1.14 | 1.14 |
| SIR3Δ 2 | 1.06 | 0.97 | 0.99 | 0.96 | 0.98 | 1.05 | 1.08 |

**Figure S6: Calculation of the correction coefficient for variability in rRNA content between different strains and timepoints. A.** We used Gel Analyzer (Istvan Lazar, www.gelanalyzer.com) to quantify the rRNA content from total RNA samples that were ran on the LabChip Nucleic Acid Analyzer. Total RNA from midlog WT1 cells is shown as an example. Background correction was performed with the rolling disk method (10% radius). The positions of 28S, 16S and 5.8S rRNA bands are indicated in the gel image below the pixel intensity profile. The rRNA bands are aligned with their corresponding pixel intensity peaks in the graph above the gel image.The rRNA fraction was determined with the formulas below the graph **B.**rRNA fraction for all RNAseq samples from Figure 7. The rRNA correction coefficient was obtained by dividing the rRNA fraction from all S.cerevisiae samples with the rRNA fraction from *S.pombe* midlog cells.

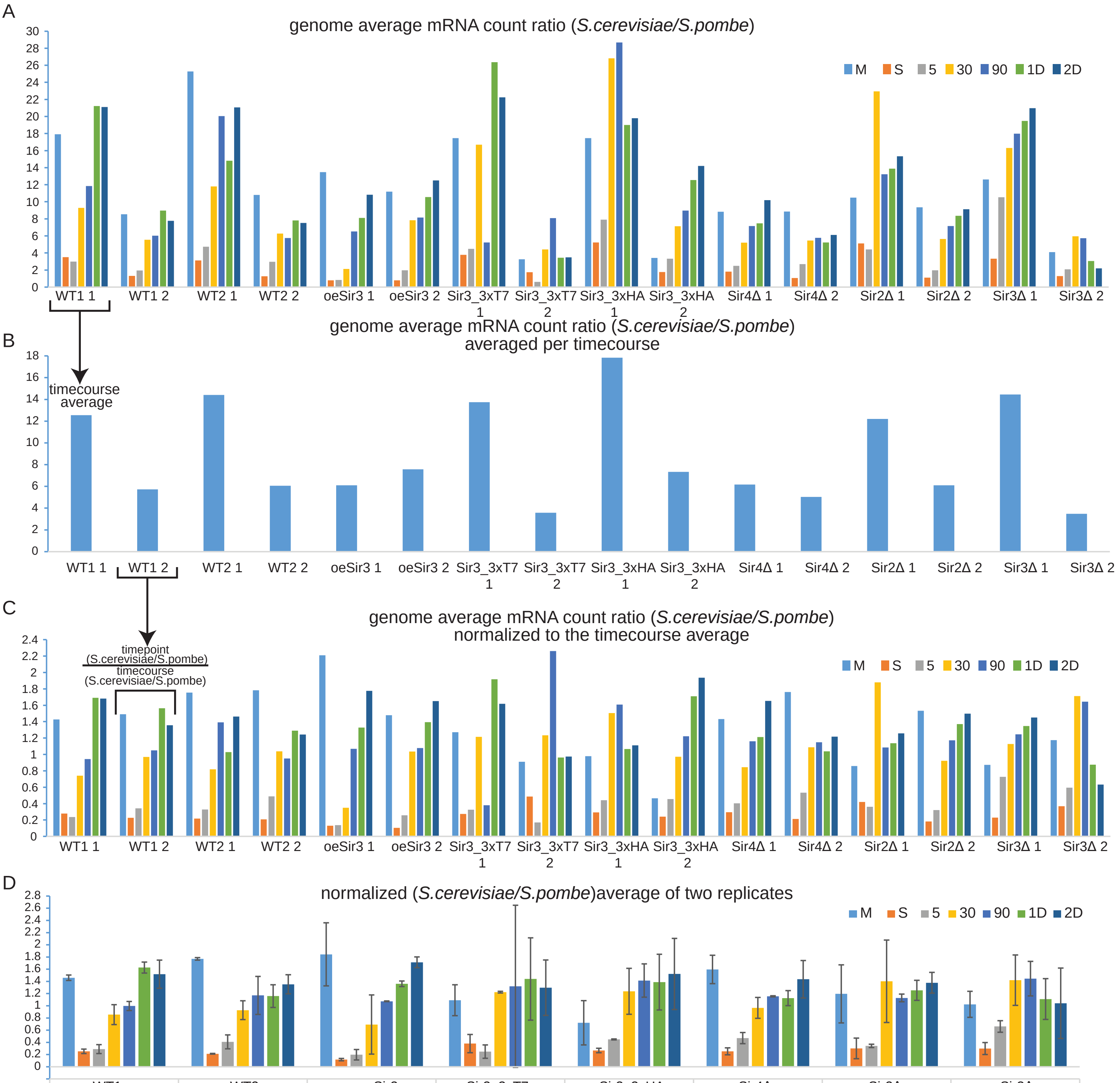


| | WT1 | WT2 | oeSir3 | Sir3_3xT7 | Sir3_3xHA | Sir4Δ | Sir2Δ | Sir3Δ |
|---|---|---|---|---|---|---|---|---|
| M | 1.46 | 1.77 | 1.84 | 1.09 | 0.72 | 1.60 | 1.20 | 1.02 |
| S | 0.25 | 0.21 | 0.12 | 0.38 | 0.27 | 0.25 | 0.30 | 0.30 |
| 5 | 0.29 | 0.41 | 0.20 | 0.25 | 0.45 | 0.47 | 0.34 | 0.66 |
| 30 | 0.86 | 0.93 | 0.69 | 1.22 | 1.24 | 0.97 | 1.40 | 1.42 |
| 90 | 1.00 | 1.17 | 1.07 | 1.32 | 1.41 | 1.15 | 1.13 | 1.44 |
| 1D | 1.63 | 1.16 | 1.36 | 1.44 | 1.39 | 1.13 | 1.25 | 1.11 |
| 2D | 1.52 | 1.35 | 1.71 | 1.30 | 1.52 | 1.44 | 1.38 | 1.04 |

**Figure S7: Faster restoration of global mRNA to pre-starvation midlog levels in Sir complex mutants and Sir3 hypomorphs A.** ratios of *S.cerevisiae* and *S.pombe* genome-wide average mRNA read counts for each time point in each strain/replicate as indicated in the x-axis. **B.** average *S.cerevisiae/S.pombe* mRNA count ratios of all time points within each time course. These average ratios were used to normalize median mRNA densities of each gene for each time point within each timecourse from Fig. 7. **C.** The ratios of each time point from A were divided by the corresponding average ratio from the corresponding time course from C, to allow for direct comparisons between strains and replicates. **D.** The normalized time point ratios from C were averaged between biological replicates of the same strain. The error bars represent the standard error between two replicates. Pre-starvation midlog mRNA levels are restored 30 min after release from starvation in Sir mutants and Sir3 hypomorphs versus 90min for WT1, WT2 and oeSir3.

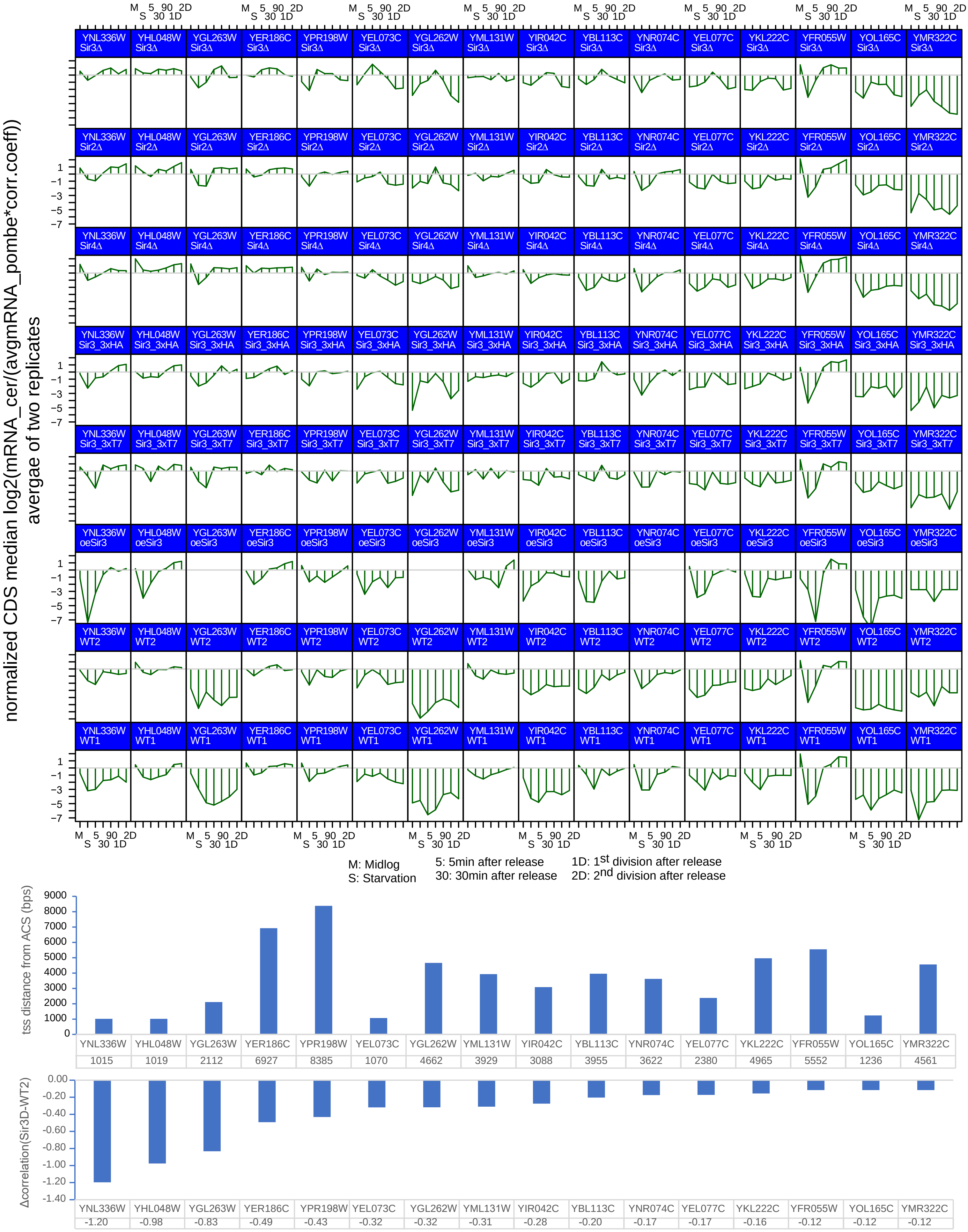


**Figure S8:** Gene expression dynamics after exit from starvation of subtelomeric genes responsive to Sir3 levels (top). We selected genes within 15kbp of subtelomeric Sir3p nucleation sites from the RNA-seq datasets in Figure 7 (middle panel) and kept the genes whose expression profiles were the most different between wt and Sir3Δ (Δcorrelation(Sir3Δ-WT2) between -0.12 and -1.2) as shown in the bar graph on the bottom.

A

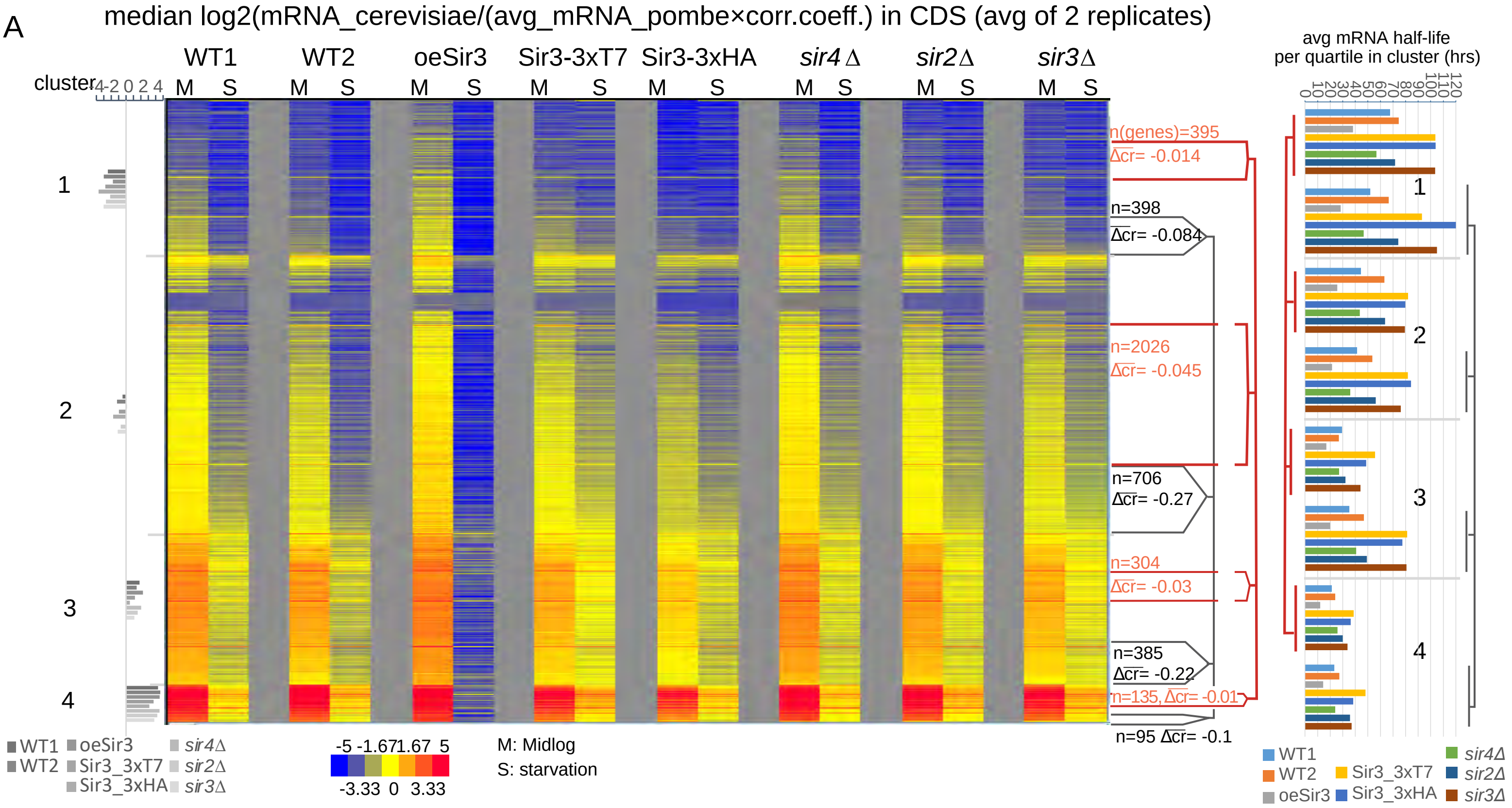


$$2^{mRNA_S}=(2^{mRNA_M})*e^{(k_S-k_d)*t}$$

$mRNA_S$=log2(mRNA_cerevisiae/(avg_mRNA_pombe*corr.coeff.)) in starvation
$mRNA_M$=log2(mRNA_cerevisiae/(avg_mRNA_pombe*corr.coeff.)) in midlog

for synthesis rate $k_S$(starvation)~0 and t=72hrs =>

$$\text{apparent decay rate (hrs}^{-1}\text{)}=k_{da}=\frac{\ln(2^{(mRNA_S-mRNA_M)})}{-72\text{hrs}}$$

$$\text{half-life (hrs)}=\frac{\ln(2)}{k_{da}}$$

B

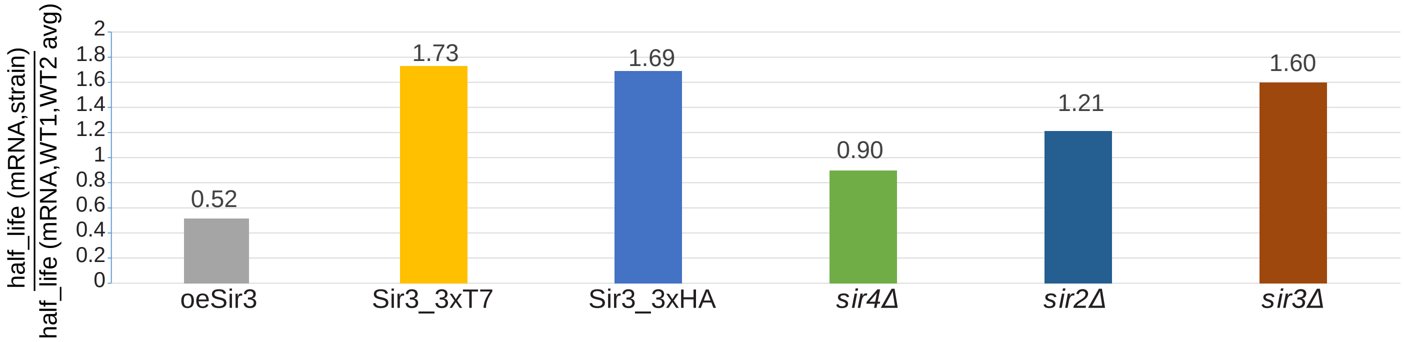


**Figure S9: Apparent mRNA half-lives during growth arrest caused by starvation. A.** The heat map shows mRNA levels in midlog (M) and starvation (S) from the RNA-seq experiment from Figure 7. Genes are organized as in Figure 7A. mRNA decay rates ($k_d$) and half lives for each gene were determined from the decrease in mRNA levels between midlog( mRNAM) and starvation (mRNAS) according to the equation shown in red below the heat map. We assumed that mRNA synthesis and decay are exponential and that the mRNA synthesis rate (ks) is close to 0 during growth arrest (starvation) for the majority of yeast genes. The average mRNA half lives in gene quartiles with *sir3*Δ most similar to WT1/2 (framed in red) or least similar to WT1/2 (framed in grey) in each cluster are shown in the bar graph on the right. The apparent mRNA half lives in *sir3*Δ and Sir3 hypomorphs (3xT7 and 3xHA) are longer than in WT1and 2 in all clusters and gene quartiles. **B.** genome average apparent mRNA half life ratios between indicated strains and the average of WT1 and WT2 strains. If we assume that the true genome average mRNA decay rate ($k_d$) is the same in all strains, the differences in the measured genome average apparent decay rate ($k_{da}$) between different strains will be determined by $k_S$>0 in each strain except oeSir where $k_S$≥0. It consequently follows from $k_{da}$(oeSir3) > $k_{da}$(*sir4*Δ) > $k_{da}$(wt) > $k_{da}$(*sir2*Δ) > $k_{da}$(*sir3* Δ)> $k_{da}$(sir3_3xHA) > $k_{da}$(Sir3_3xT7) that the genome average mRNA synthesis rates during starvation are $k_S$(oeSir3) < $k_S$(*sir4*Δ) < $k_S$(wt) < $k_S$(*sir2*Δ) < $k_S$(*sir3*Δ) < $k_S$(Sir3_3xHA) < $k_S$(Sir3_3xT7).strain

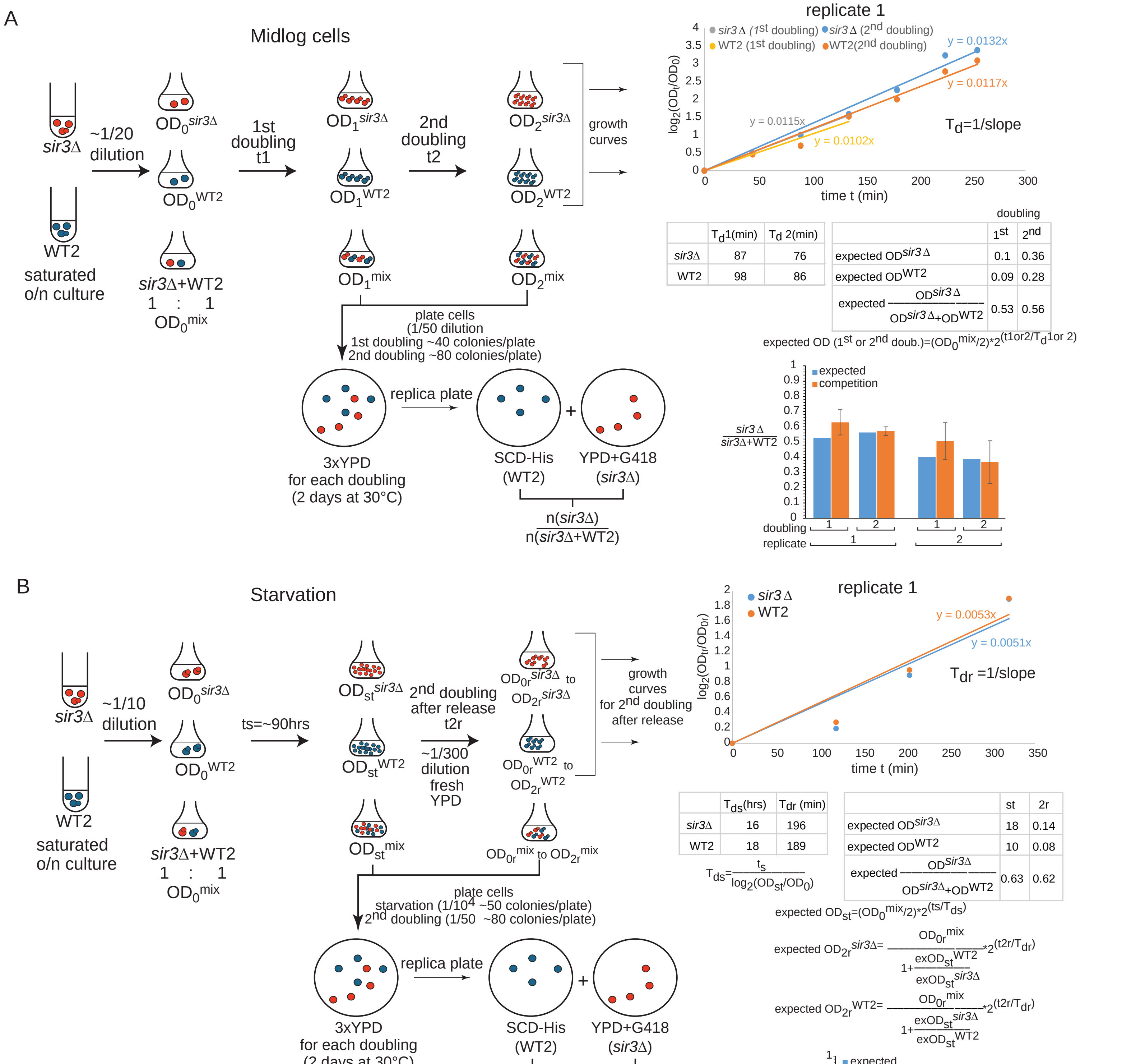


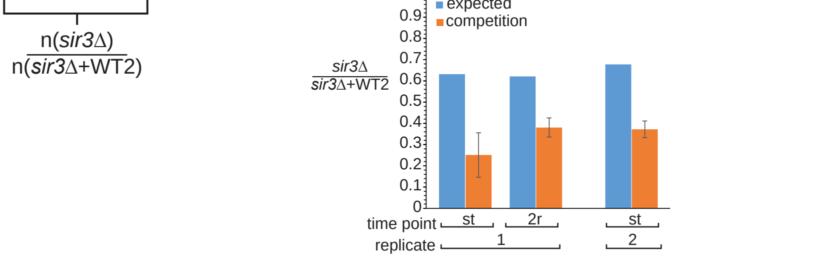

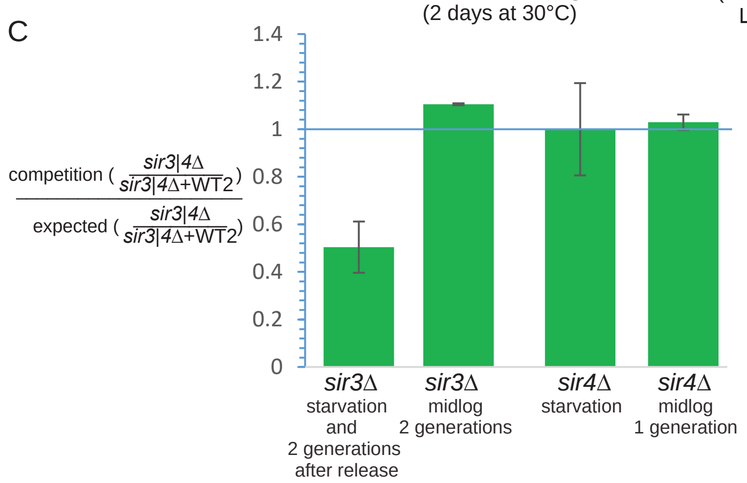


**Figure S10: Competitive fitness test between *sir3*Δ or *sir4*Δ and WT2. A.** Competition test during exponential growth. Left: outline of the experiment.Saturated *sir3*Δ and WT2 cultures were diluted 20fold in fresh YPD separately or mixed at a 1:1 OD ratio and grown for 2 generations. Individual cultures are used to calculate individual growth rates for the 1st and 2nd doubling(scatter plot on the right), used to determine the expected *sir3*Δ fraction in the mix (expected ($OD^{sir3\Delta}/(OD^{sir3\Delta}+OD^{WT2})$) if neither strain is influenced by the presence of the other strain in the mix (the tables on the right below the scatter plot shows values obtained from the first biological replicate). To determine the *sir3*Δ fraction when the two strains are competeing with each other, the mixed culture was grown in parallel with the individual cultures and cells from the mix were plated on 3 YPD plates after the 1st and the 2nd doubling. Each YPD plate was than replica plated on a SCD-His (selects for WT2) and a YPD+G418 plate (selects for *sir3*Δ) and individual colonies were counted on each selection plate to get the experimental *sir3*Δ fraction (n(*sir3*Δ)/n(*sir3*Δ+nWT2)) . The bar graph on the bottom right shows the expected (blue) and the experimental (competition, orange) *sir3*Δ fractions in the mixed culture for two biological replicates and two generations. The error bars represent the standard error from the average of 3 plates.**B.** As in A but for starvation conditions. Saturated o/n *sir3*Δ and WT2 cultures were diluted ($OD_0$) and mixed before starvation (cells were left at 30°C for ts=~90hrs). An aliquot was plated on 3xYPD plates after 90hrs of starvation (the st time point) and another aliquot from the starved culture was diluted ($OD_{0r}$) into fresh YPD. After two doublings the mixed culture was plated on 3xYPD plates. Individual *sir3*Δ and WT2 cultures were grown in parallel in the same conditions as the mix to determine expected *sir3*Δ fractions as in A. The bar graph on the bottom left shows expected and experimental *sir3*Δ fraction for two biological replicates at indicated time points (the time point after release from starvation was only taken in the first replicate).The error bars represent the standard error from the average of 3 plates as in A. **C.** Ratio between experimental (competition) and expected *sir3*Δ or *sir4*Δ fraction in the mixed population during starvation and exponential growth. The *sir3*Δ ratios are averaged over two biological replicates and time points for each replicate from B and A respectively (the starvation condition is averaged from two starvation time points from two replicates and 1 release time point from 1 replicate). The *sir4*Δ ratios are averaged over two biological replicates. The starvation condition is averaged from two starvation time points: 70.25hrs and 90hrs for replicates 1 and 2, respectively. *sir4*Δ colonies are also selected on YPD+G418 plates. The error bars represent the standard error from the average of replicates and time points.
