## Supplemental Methods for "The budding yeast heterochromatic protein Sir3 modulates genome-wide gene expression through transient direct contacts with euchromatin"

**SI Appendix**

### Materials and Methods

#### Yeast Strains

The Sir3-3XHA to 3xT7 RITE strain used in Figures 3, 4, 5, 7 and S3-S9 is MRL9.1 (S288C: *MATa his3d200 leu2d0 lys2d0 met15d0 trp1d63 ura3d0 bar1::HisG HIS3 Pgpd_CRE_EBD78 Sir3Histag-LoxP-3xHA-Hygro-LoxP-3xT7*) that was constructed by transforming NKI12318 (*MATa his3d200 leu2d0 lys2d0 met15d0 trp1d63 ura3d0 bar1::HisG HIS3 Pgpd_CRE_EBD78*; strain WT1 in Figures 3, 4, 7 and S6-S9 courtesy of Fred van Leeuwen) with the RITE-switch (*LoxP-3xHA-Hygro-LoxP-3xT7*) cassette amplified from the pFvL159 plasmid (Fred van Leeuwen) with primers (
forward:
5'GCCTTTTCGATGGATGAAGAATTCAAAAATATGGACTGCATTCATCACCATCACCATCACGGTGGATCTGGTGGATCT;
reverse:
5'CATAGGCATATCTATGGCGGAAGTGAAAATGAATGTTGGTGGTGATTACGCCAAGCTCG) compatible for homologous recombination with the C-terminal end of the Sir3 gene. Cassette incorporation was verified by PCR.

The Sir3-3xXHA to 3xT7 RITE strain without the His-tag linker used in Figure 4 is MM1 (S288C: *MATa his3d200 leu2d0 lys2d0 met15d0 trp1d63 ura3d0 bar1::HisG HIS3 Pgpd_CRE_EBD78 Sir3-LoxP-3xHA-Hygro-LoxP-3xT7*) that was constructed by transforming NKI12318 with the RITE-switch (*LoxP-3xHA-Hygro-LoxP-3xT7*) cassette amplified from the pFvL159 plasmid (Fred van Leeuwen) as above (the PCR primers are the same except they don't have the 6xHis tag linker).

The Sir3-1xT7 to 1xHA RITE strain used in Figures 4 and S3 is MM3 (S288C: *MATa his3d200 leu2d0 lys2d0 met15d0 trp1d63 ura3d0 bar1::HisG HIS3 Pgpd_CRE_EBD78 Sir3-LoxP-1xHA-Hygro-LoxP-1xT7*) that was constructed by transforming NKI12318 with the RITE-switch (*LoxP-1xHA-Hygro-LoxP-1xT7*) cassette amplified from the pTW081 plasmid (Fred van Leeuwen) as above.

The Sir3-1XHA to 1xT7 RITE strain used in Figure 4 is MM2 (S288C: *MATa his3d200 leu2d0 lys2d0 met15d0 trp1d63 ura3d0 bar1::HisG HIS3 Pgpd_CRE_EBD78 Sir3-LoxP-1xHA-Hygro-LoxP-1xT7*) that was constructed by transforming NKI12318 with the RITE-switch (*LoxP-1xHA-Hygro-LoxP-1xT7*) cassette amplified from the pFvL1118 plasmid (Fred van Leeuwen) as above.

The Sir3 over-expression strain oeSir3 in Figures 3, 4, 6 and 7 and S6-S9 is THC70 (W303: *MATa HMLa HMRa ade2-1 can1-100 his3-11,15 leu2-3,112 trp1-1 mat*Δ*::TRP1 hmr::rHMRa hml*Δ*::kanMX ura3-1::GAL10P–Sir3HA::URA3 bar1*Δ*::hisG lys2*Δ *sir3*Δ*::HIS3*) (courtesy of K.Struhl (1, 2)).

The WT2 strain from Figures 3, 7, and S6-S10 is JOY1 (S288C(BY4741 parent): *MATa ura3D leu2D his3D met15D bar1D:HIS5*) and has the same genetic background as the s*ir3Δ* strain from Figures 7, and S6-S10 (MRL5.1 ; S288C: *MATa ura3Δ leu2Δ his3Δ met15Δ sir3Δ:KANR*) and the *sir2Δ* and *sir4Δ* strains from Figures 7, and S6-S10 are from the YSC1053_KO deletion library (GE Healthcare/Dharmacon) (*MATa ura3Δ leu2Δ his3Δ 1 met15Δ Sir2|4Δ::KanR*).

The Sir3Dam strain (MATalpha *ade2-1 ura3-1 his3-11,15 trp1-1 leu2-3,112 can1-100 SIR3*:: Myc9-DAM (methylase from *E.coli*) Phleomycin) from Figures 1, 3A, and S1, were constructed by transformation of the W303 parent with a Dam-Phleomycin PCR fragment from the pBleomycin-Myc-Dam plasmid. The Dam gene was amplified from genomic DNA extracted from *E.coli* (DH5alpha) and inserted into the MC site of the pMX-Bleomycin vector. The addition of a 9xMyc tag at the N terminus of Dam resulted in the pBleomycin-Myc-Dam plasmid. The pBleomycin-Myc-DamK9A plasmid was constructed by introducing the K9A mutation by PCR into the Dam gene on the pBleomycin-Myc-Dam plasmid. The mutation was verified by sequencing. The Sir3DamK9A strain (MATa *ade2-1 ura3-1 his3-11,15 trp1-1 leu2-3,112 can1-100 SIR3*:: Myc9-DAMK9A (methylase from *E.coli*) Phleomycin) from Figures 1 and S1 was then constructed as the Sir3Dam strain using a PCR fragment from the pBleomycin-Myc-DamK9A plasmid.

The Sir3EcoG2 (*MATalpha ade2-1 ura3-1 his3-11,15 trp1-1 leu2-3,112 can1-100 SIR3:: Myc9-EcoGII* (methylase from *E.coli*) Phleomycin), Sir2EcoG2 (*MATa ade2-1 ura3-1 his3-11,15 trp1-1 leu2-3,112 can1-100 SIR2:: Myc9-EcoGII* (methylase from *E.coli*) Phleomycin) and Sir4EcoG2 (MATa *ade2-1 ura3-1 his3-11,15 trp1-1 leu2-3,112 can1-100 SIR4*:: Myc9-EcoGII (methylase from *E.coli*) Phleomycin) strains from Figures 1, 2, 8, and S2 were constructed as the Sir3Dam strain from the pBleomycin-Myc-EcoG2 plasmid, which was constructed by replacing the Dam gene in the pBleomycin-Myc-Dam plasmid with the EcoG2 gene amplified from the pLPC-M.EcoG2 plasmid (courtesy of L. Crabbe, (3)). The Sir3EcoG2_*sir2Δ* ((MATalpha *ade2-1 ura3-1 his3-11,15 trp1-1 leu2-3,112 can1-100 SIR3*:: Myc9-EcoGII (methylase from *E.coli*) Phleomycin *sir2*Δ::hygR), Sir3EcoG2_*sir4Δ* ((*MATalpha ade2-1 ura3-1 his3-11,15 trp1-1 leu2-3,112 can1-100 SIR3:: Myc9-EcoGII* (methylase from *E.coli*) Phleomycin *sir4*Δ::*HYGR*) and Sir3EcoG2_*sir2Δsir4Δ* ((*MATalpha ade2-1 ura3-1 his3-11,15 trp1-1 leu2-3,112 can1-100 SIR3:: Myc9-EcoGII* (methylase from *E.coli*) Phleomycin *sir2*Δ::hygR *sir4*Δ::*KANR*) were constructed by transforming the Sir3EcoG2 parent strain with a PCR product carrying the appropriate antibiotic resistance gene.

The NLSDamK9A (*MATa ura3-1 his3-11,15 trp1-1 leu2-3,112 can1-100 ade2-1:: promoterSIR3-Myc9-NLSDAM K9A ::ADE2*) and NLSEcoG2 (*MATa ura3-1 his3-11,15 trp1-1 leu2-3,112 can1-100 MATa ade2-1::promotorSIR3-NLS-EcoGII::ADE2*) strains were constructed by respectively transforming the Ade2 linearized (with StuI) integrative plasmids pADE2-pSIR3-NLS-DAMK9A and pADE2-pSIR3-NLS-EcoGII into a W303 parent. pSir3 is the Sir3 promoter amplified from W303 genomic DNA and cloned into the pADE2 integrative plasmid vector. The NLS

(ACGCTAGCATGCCAAAAAAGAAACGCAAGGTGCTTAATACTGTAAAAATATCCAGT) coding for SV40 minimum NLS MPKKKRKV, was cloned into the pADE2 vector downstream of the pSir3 sequence and the DAMK9A and EcoG2 sequences from the pBleomycin-Myc-DamK9A and pBleomycin-Myc-EcoG2 plasmid, respectively, were then inserted downstream of the NLS sequence.

**Sir3 Nanopore MetID cell cultures and genomic DNA preparation for nanopore sequencing**

For mid-log cultures, cell were grown overnight in YPD at 30°C until they reached saturation (~$10^8$ cells/ml). For the release from starvation time course (Figure 3A), cells were grown in YPD at 30°C for 72hrs, an aliquot was taken for the starvation time point. Cells were then pelleted and re-suspended in fresh YPD to OD=0.3. 300ml aliquots were then taken at indicated times, cells were pelleted and kept on ice until spheroplasting.

Pelleted cells were counted and divided into aliquots of $10^9$ cells, the amount needed for one sequencing run. Pellets were then re-suspended in 1M Sorbitol and incubated with 25 U of Zymolyase (AMSBIO) for 30min at 30°C for 30min with gentle shaking (300rpm, thermomixer).

High Molecular Weight DNA was then extracted from pelleted spheroplasts with the MagAttract HMW DNA Kit (Qiagen) according to the manufacturer's protocol.

400ng from each genomic DNA sample were used for nanopre sequencing library preparation with the Rapid Barcoding Kit (Oxford Nanopore) or with the Genomic DNA by Ligation kit (Oxford Nanopore) for the Sir3Dam saturated culture sample, according to the manufacturer protocol.

Libraries prepared with the Rapid Barcoding Kit were mixed, cleaned, and concentrated with Magna or Ampure beads as described in the Rapid Barcoding Kit protocol. The library mix was loaded on the R9.4.1 Flow cell (Oxford Nanopore) (Sir3Dam and Dam_without_NLS from a saturated culture were loaded on a Flongle R9.4.1 flow cell) and sequenced with the Minion device (Oxford Nanopre) for 18 to 24 hrs without the real time base calling option.

**Sir3 tag switch cell culture**

All cultures were incubated at 30˚C in an incubator shaker at 220 rpm, crosslinked for 20 min with 1% formaldehyde and quenched for 5 min with 125mM Glycine, unless indicated otherwise. Two 10 ml cultures were grown overnight in YPD (2% glucose). One culture contained hygromycin (0.3 mg/ml; the Hyg+ culture). Hyg+ and Hyg- saturated cultures were then transferred to flasks with 90 ml YPD and hygromycin was added to the Hyg+ culture and incubated for 48h until glucose was depleted and cells stopped dividing (monitored by OD measurements). A 20 ml aliquot from the Hyg+ culture was fixed, pelleted and flash frozen in liquid nitrogen and kept at -80°C as the "before switch" starvation sample. The rest of the Hyg+ and Hyg- cultures were pelleted and 80 ml of the Hyg- supernatant was used to re-suspend the Hyg+ pellet, and the Hyg+ supernatant and Hyg- pellet were discarded. The re-suspended 80 ml culture was inoculated with 1 µM estradiol (Sigma) in order to induce tag exchange, and incubated overnight for at least 16h. An aliquot for the "after switch" starvation sample was processed as above. The remaining culture was diluted to OD 0.3 with fresh pre-warmed (30°C) YPD (total volume: 1600-2000 ml) and incubated at 30°C to release the culture from starvation. 400 ml aliquots were taken at 5, 30 and 90 min after release and after indicated cell doublings (monitored by OD measurements; the first doubling typically takes place 3.5 hours after release and each subsequent doubling takes 1.5hrs). Aliquots for each time point were processed as above and fresh YPD (preheated at 30°C) was added to the rest of the culture in order to keep cell density constant (constant OD) and maintain cells in exponential growth up to the 4$^{th}$ or 5$^{th}$ doubling. For the 12$^{th}$ doubling after release in Figure S5, 400ml of fresh pre-warmed YPD were inoculated with a small volume of the cell culture after the 4$^{th}$ doubling and grown over night. The inoculation volume was calculated so the overnight culture reaches an OD between 0.3 and 0.5 the next day.

Small cell aliquots (50µl from a 1:100000 dilution) before and after tag switch were plated on YPD plates and replica-plated on YPD+hygromycin to estimate recombination efficiency. The average recombination efficiency in our cell culture conditions is 96.9% (from 11 independent experiments).

**Western blot**

10 ml aliquots from each time point were mixed with 2 ml 100% TCA and kept on ice for 10 min. Cells were then pelleted and washed twice with 500 µl 10% cold TCA. Pellets were re-suspended in 300 µl 10% cold TCA and bead beated with Zirconium Sillicate beads (0.5 mm) in a bullet blender (Next Advance) for 3 times 3 min (intensity 8). Zirconium beads were removed from the cell lysate by centrifugation and the entire cell lysate was washed twice with 200 µl 10% cold TCA. The cells lysate was then pelleted and re-suspended in 70 µl 2xSDS loading buffer (125 mM Tris pH 6.8, 20% glycerol; 4% SDS, 10% β-mercaptoethanol, 0.004% bromophenol blue) preheated at 95˚C. Approximately 30 µl Tris (1M, pH 8.7) was added to each sample to stabilize the pH. Samples were heated for 10 min at 95˚C, pelleted and the soluble protein extract in the supernatant was transferred to new tubes. Protein concentrations were measured by Bradford test kit (Sigma, B6916) and 40 µg/sample was loaded on a 7% polyacrylamide SDS-PAGE gel (30:1 acrylamide/bis-acrylamide). Proteins were transferred after electrophoresis onto a PDVF membrane (Bio-Rad, 1620177). The membrane was incubated for 1h at room temperature with either anti-HA (Abcam, ab9110 (lot# GR3245707-3)) for replicate 1 or anti-HA (Roche (Sigma) 11867423001 (lot# 27573500)) for replicate 2, to detect "old" Sir3 and anti-α-tubulin (Sigma T6199 (lot#116M48C2V)) to detect the α-tubulin loading control. Secondary goat anti-rat-HRP (Santa Cruz Biotechnology, SC2006, lot# G2514) and bovine anti-mouse-HRP (Santa Cruz Biotechnology sc-2375) were added after the corresponding primary antibody and incubated for 1hr at room temperature. Primary antibodies were diluted 1/1000 (replicate 1) or 1/500 (replicate 2) and secondary antibodies were diluted 1/10000 in 5% milk/TBS. The membrane was washed 3 times in 1xTBS-10%Tween after each antibody incubation step. The blot was then covered with 500 µl Immobilon Forte Western HRP substrate (Millipore WBLUF0500) for 2 min and protein bands were detected on a high-performance chemiluminescent film (Amersham 28906837).

**Microscopy and image analysis**

Cells were grown in YPD for 72hrs as above to obtain cultures in growth arrest (the tag switch step was omitted in this experiment) or overnight to OD=0.5 (for mid-log cells), and concentrated by centrifugation. 4µl of the cell pellet was injected under the 0.8% agarose/YPD/α-factor (0.2µg/ml) layer that had been poured into each well of an 8-well glass bottom microscopy plate (BioValley). oeSir3 cells (THC70) were grown as above in YPGal (2%) or YPD and injected under 0.8% agarose/YPD/α-factor (0.2µg/ml) or 0.8% agarose /YPGal/α-factor (0.2µg/ml) layers as indicated in Figure 4.

Images were acquired using a Nikon Ti2 Eclipse wide field inverted microscope in the triple channel LED DIC mode with a Nikon Plan Apochromat 60x water objective, NA 1.2, and a Photometrics Prime 95B CMOS camera (1200*1200, 11µm pixel size). Cells were kept at 30°C and imaged every 10min for 6 hrs, exposure time was 40 ms (Figure 4B-D). For oeSir3 cells, we used a wide field inverted microscope for TIRF acquisition (Nikon) under the HiLo setting, with a 60X water objective and water dispenser, and a EMCCD Evolve 512 Photometrics camera (512*512, 16µm pixel size). The time courses on growing cells were performed at 30°C. Pictures in bright field (Exposure time= 300ms and the Hilo angle= 62°) were taken every 10 or 20min for 6.5hrs (Figure 4E).

Shmoon and budding cells were counted manually using FIJI (Image J) for visualization.

**Chromatin Sonication**

Cross-linked frozen cell pellets were re-suspended in 500 µl cell breaking buffer (20% glycerol, 100 mM Tris pH 7.5, 1xEDTA-free protease inhibitor cocktail (Roche)). Zirconium Sillicate beads (400 µl, 0.5 mm) were then added to each aliquot and cells were mechanically disrupted using a bullet blender (Next Advance) for 5 x 3 min (intensity 8). Zirconium beads were removed from the cell lysate by centrifugation and the entire cell lysate was subject to sonication using the Bioruptor-Pico (Diagenode) for 3x10 cycles of 30 seconds ON/OFF each for a 500bp final median size of chromatin fragments. Cellular debris was then removed by centrifugation and the supernatant was used for ChIP.

**ChIP for ChIP-seq**

All steps were done at 4˚C unless indicated otherwise. For each aliquot, Buffer L (50 mM Hepes-KOH pH 7.5, 140 mM NaCl, 1 mM EDTA, 1% Triton X-100, 0.1% sodium deoxycholate) components were added from concentrated stocks (10-20X) for a total volume of 0.8 ml per aliquot. For anti-HA and anti-T7 ChIP, each aliquot was rotated for 1 hour with 100 µl 50% Sepharose Protein A Fast-Flow bead slurry (IPA400HC, Repligen) previously equilibrated in Buffer L. The beads were pelleted at 3000g for 30s, and approximately 200 µl of the supernatant was set aside for the input sample. The remainder (equivalent to 200 ml of cell culture of 0.5 OD) was separated into anti-HA and anti-T7 fractions. 10 µl anti-HA (Abcam, ab9110 (lot# GR3245707-3) and 10 µl polyclonal anti-T7 (Bethyl A190-117A (lot# A190-117A-7) (Figure 5) or 10 µl monoclonal anti-T7 (Cell Signaling Technology, DSE1X (lot#1)) (Supplementary Figure S5) were added to the corresponding aliquots. Beads were washed successively with Buffers L, W1(Buffer L with 500mM NaCl), W2(10mM Tris pH

8, 250mM LiCl, 0.5% NP-40, 0.5% Na-deoxycholate, 1mM EDTA) and TE (10mM Tris pH 8, 1mM EDTA), twice each, for 5min at 4°C with rotation. Samples were eluted twice in 125µl Elution buffer (TE, 1%SDS and 150mM NaCl with freshly added 5mM DTT) after 2x10min incubation at 65°C.

Purified DNA was treated with RNAse A (Qiagen) (5µg per sample for 1hr at 37°C) and purified once more with Phenol-Chloroform. Purified fragments were used for NGS library construction (Input, ChIP.

**Cell culture and RNA isolation for RNA-seq with *S.pombe* "spike in"**

All cells were grown as above in YPD for 72 hours and Sir3-3xHA was not subject to tag switching with estradiol. oeSir3 was grown in conditions of Sir3 over-expression in YPGalactose (2%). Cells were pelleted (3500rpm, 1min, 30°C) and flash frozen in liquid N2 at indicated times and total RNA was isolated from frozen cell pellets with TRIzol (Sigma). The 5min time point is taken immediately after release into new media but cells have effectively spent ~5min in new media during resuspension and centrifugation (at 30°C) before freezing. Consequently, some growth response genes have already been induced by that time. Frozen cell pellets were re-suspended directly in TRIzol and bead beated in the Bullet Blender (Next Advance) for 4 times, 3 min each (intensity 8). TRIzol extracted total RNA was then purified and DNAseI treated with the RNAeasy Column purification kit (Qiagen). RNA quality was checked by Bioanalyzer scan (Agilent) or with Qubit ™ RNA IQ Assay kit (Thermo Fisher Scientific). Purified total RNA amounts were measured in a nanodrop spectrophotometer. 2 µg (ML, 1D and 2D) or 8 µg (Starvation, 5, 30 and 90 min) of *S. cerevisiae* total RNA samples were mixed with total RNA from *S. Pombe* (strain FY2319, courtesy of S. Forsburg) mid-log cultures (grown in YES, flash frozen in liquid $N_2$ and extracted with TRIzol as above) at a 10:1 mass to mass ratio. NGS libraries for WT1 2, WT2 2, oeSir3 1, oeSir3 2, Sir3-3xT7 2, Sir3-3xHA 2, *sir4Δ* 1, *sir4Δ* 2, *sir2Δ* 2 and *sir3Δ* 2 were prepared using the Illumina Stranded mRNA Prep Ligation kit according to manufacturer's instructions. NGS Libraries for WT1 1, WT2 1, Sir3-3xT7 1, Sir3-3xHA 1, *sir2Δ* 1, and *sir3Δ* 1 were made using the Illumina TruSeq Stranded mRNA kit according to the manufacturer's protocol. 1 and 2 in the sample name refer to the replicate number. Libraries were sequenced on the Illumina NextSeq550 (2x75bp) (Plateforme Transcriptome, IRMB, Montpellier, France) or NovaSeq 6000 (2x75bp) (Illumina) at the CNAG, Barcelona.

**rRNA content calculation**

We checked the rRNA content of our purified total RNA samples in a 2100 Bioanalyzer Instrument (Agilent) using the Bioanalyzer RNA 6000 Nano assay or in a LabChip GX HT Touch Nucleic Acid Analyzer (Perkin Elmer) with the HT RNA Standard Sensitivity assay kit according the manufacturers' protocol. All samples were diluted to 500 ng/µl for Bioanalyzer and 100 ng/µl for LabChip assays. Gel images obtained from Bioanalyzer/LabChip were visualized in GelAnalyzer 19.1. (www.gelanalyzer.com). Lanes of equal length were selected manually, corresponding profiles were checked for automatic peak detection using the default parameters: peak threshold=1, minimum peak height=5 and maximum peak width in percentage of lane profile length. This allowed the detection of the rRNA peaks and the marker peaks (excluded manually). The peaks for the mRNA regions and the small RNA peaks, as shown in Figure S6A, were added manually. Baselines were subtracted using the rolling ball method with a diameter of 10% of the corresponding total lane profile length. The raw pixel intensities were then exported to Excel and the rRNA fractions were calculated as described in Figure S7.

**Competitive Fitness Assay**

For competition during exponential growth, o/n saturated *sir3Δ*, *sir4Δ* and WT2 cultures ( 5ml YPD inoculated with one colony and grown o/n at 30°C) were diluted ~20 fold (final OD ~0.1) with fresh YPD and grown for two generations either as individual cultures or as a mix with 1:1 OD ratio of WT2 and *sir3Δ* or *sir4Δ* cells (incubator shaker 30°C). OD measurements were taken at regular intervals during growth. The ODs collected from individual cultures were used to calculate the expected ratio of *sir3Δ* or *sir4Δ* and WT cells in the mix after the first and second doubling as described in Supplementary Figure S10. 1/50 dilutions of cells from the mixed culture were plated on 3 YPD plates after the first and after the second doubling. YPD plates were kept at 30°C for 2 to 4 days before replica plating on SCD-His (selects for WT2) and YPD+G418 (200µg/ml, selects for *sir3Δ* or *sir4Δ*) selection plates (incubated 1 night at 30°C). Colonies on each selection plate were counted the next day.

For competition during starvation, o/n saturated *sir3Δ* or *sir4Δ* and WT2 cultures were diluted 10 fold with fresh YPD and left in an incubator shaker at 30°C for ~90hrs. Individual *sir3Δ* or *sir4Δ* and WT2 and a mixture of WT2 with either mutant (1:1 OD ratio) were incubated in parallel as above. 1/10000 (replicate 1) or 1/5000 (replicate 2) dilutions of starved mixed populations were plated on 3 YPD plates for the starvation time point. Starved cultures (*sir3Δ* or *sir4Δ*, WT2 and the mix) were then diluted 300 fold (final OD ~0.1) and grown for two generations. OD measurements were taken at regular intervals during re-growth, as above, and 1/50 dilutions of the mixed populations were first plated on 3 YPD plates after the second doubling and then replica-plated as above.

**NGS Input and ChIP library construction and Illumina sequencing**

DNA fragments were blunt ended and phosphorylated with the Epicentre End-it-Repair kit. Adenosine nucleotide overhangs were added using Epicentre exo- Klenow. Illumina Genome sequencing adaptors with in line barcodes (

PE1-NNNNN: PhosNNNNNAGATCGGAAGAGCGGTTCAGCAGGAATGCCGAG

PE2-NNNNN: ACACTCTTTCCCTACACGACGCTCTTCCGATCTNNNNNT

, NNNNN indicates the position of the 5bp barcode, (IDT) were then ligated over night at 16°C using the Epicentre Fast-Link ligation kit. Ligated fragments were amplified using the Phusion enzyme

(NEB) for 18 PCR cycles with Illumina PE1 (AATGATACGGCGACCACCGAGATCTACACTCTTTCCCTACACGACGCTCTTCCGATCT) and PE2 (CAAGCAGAAGACGGCATACGAGATCGGTCTCGGCATTCCTGCTGAACCGCTCTTCCGATCT) primers (IDT). Reactions were cleaned between each step using MagNa beads.

Libraries were mixed in equimolar amounts (12 to 21 libraries per pool) and library pools were sequenced on the HiSeq 2000 (2x75bp) (Illumina) at the CNAG, Barcelona, Spain or the NextSeq550 (2x75bp) (Plateforme Transcriptome, IRMB, Montpellier, France).

**ChIP-seq data analysis**

Sequences were aligned to the *S.cerevisiae* genome using BLAT (Kent Informatics, http://hgdownload.soe.ucsc.edu/admin/). We kept reads that had at least one uniquely aligned 100% match in the paired end pair. Read count distribution was determined in 1bp windows and then normalized to 1 by dividing each base pair count with the genome-wide average base-pair count for time point (sum of all read counts per bp divided by the number of bps in the genome (~12Mbp)). Forward and reverse reads were then averaged and ChIP reads were normalized to their corresponding input reads.

The repetitive regions map was constructed by "BLATing" all the possible 70 bp sequences of the yeast genome and parsing all the unique 70bp sequences. All the base coordinates that were not in those unique sequences were considered repetitive.

Calculation of Sir3 exchange rates from the tag switch experiment (Figure 5):

Since Sir3 with the "old" tag stops being produced after the tag switch, we can determine Sir3p OFF rates during and after exit from growth arrest from the slope of the linear fit of the decrease in "old" Sir3p enrichment at time points during growth arrest and during the first cell cycle after release. Likewise, the Sir3p ON rate can be estimated from the initial rate of increase in "new" Sir3p enrichment up to the point when "new" Sir3p enrichment reaches equilibrium. The slope of the linear fit of the change in "new" Sir3p enrichment after release is approximately equal to the Sir3p ON rate even though it actually represents the ON/OFF rate ratio of "new" Sir3p, because the OFF rate of "new" Sir3p is initially negligible since most of the bound Sir3p is the one with the "old" tag in the early time points after the tag switch. ON and OFF in Δenrichment %/hr (ΔE) were therefore calculated according to the equation: $\Delta E = (2^{|b|} - 1) * 100$, where b is the slope of the linear fits from Figure 5C-E.

Calculation of Sir3 exchange rates from the ChIP seq experiment with the oeSir3 strain (Figure 6): The OFF rate is directly estimated from the rate of decrease of Sir3p enrichment after release from growth arrest into dextrose. The ON rate can then be calculated from the increase in Sir3 enrichment after release in galactose and the OFF rate measured in dextrose according to the formula:

$ON\ (\%/hr) = (2^{(b_i+b_d)} - 1) * 100$, where $b_i$ and $b_d$ are the slopes of the linear fits for the rates of increase in Sir3 enrichment in galactose and decrease in Sir3p enrichment in dextrose, respectively.

**RNA-seq data analysis**

*S.cerevisiae* and *S.pombe* reads were aligned to their respective genomes using BLAT and the read density distribution was determined for each species in each dataset separately. The average *S.pombe* genomic read density per bp (F and R reads were processed together) was determined for each dataset. For spike-in normalization, *S.cerevisiae* read densities per bp were then divided with the corresponding corrected average *S.pombe* genomic read density. The average *S.pombe* genomic read density was corrected by multiplying it with the corresponding rRNA correction coefficient (Figure S6) to account for variability in rRNA content between samples. Normalized read densities for each gene were aligned by the transcription start site and divided into sense and antisense transcripts. The median read density for each gene (from the tss to the end of the coding sequence) was then determined for each transcript. Intron regions were excluded from the calculation. Median read densities for each gene in each replicate of each time point were finally divided by the genome average mRNA *S.cerevisiae*/*S.pombe* count ratio for the whole time course of the corresponding replicate, to allow for direct comparisons between replicates and strains, as described in Figure S7.

**Nanopore sequencing data analysis**

Base calling was performed on raw fast5 files, using guppy basecaller (ONT, Community - Downloads (nanoporetech.com)) with dna_r9.4.1_450bps_modbases_dam-dcm-cpg_hac.cfg for Dam signals and the res_dna_r941_min_modbases-all-context_v001.cfg (rerio/basecall_models at master · nanoporetech/rerio · GitHub) for EcoG2 signals. Fastq files and modification probability tables were then extracted from base-called fast5 files using the ont-fast5-api package (GitHub - nanoporetech/ont_fast5_api: Oxford Nanopore Technologies fast5 API software). Demultiplexing and genome alignments were done with the guppy barcoder and the guppy aligner, respectively (S.cerevisiae reference genome: S288C_reference_sequence_R64-3-1_20210421.fasta). Adenines with a methylation probability higher or equal to 0.95 were considered as positive signals.

Sir3Dam and Sir3DamK9A analysis

The $^{me}$A count in each 400bp region of the genome was divided by the number of sequenced reads and the number of GATCs for that segment to account for the heterogeneous distribution of GATC motifs, which ranges from 0 to 6 per 400bp (Fig. S1B). 400bp regions without GATCs were excluded from the analysis. The normalized n($^{me}$A) signal ([n($^{me}$A)/(n(reads)*n(GATC))] per 400bp region) from the Sir3Dam and Sir3DamK9A strains were then subtracted from the normalized n($^{me}$A) signal from the untagged wt parent strain to eliminate false positives and from the (pSIR3)NLSDamK9A (standalone DamK9A with an NLS sequence under the control of the SIR3 promoter) signal to eliminate methylated sites that are not specific to Sir3. Positive and negative values consequently represent target sites that are specific for Sir3Dam|Sir3DamK9A and standalone Dam, respectively.

Sir3EcoG2, Sir2EcoG2 and Sir4EcoG2 analysis

The $^{me}$A count in the Watson or Crick strand of each 400bp region of the genome was divided by the number of sequenced reads of each strand from the Sir3|2|4 EcoG2 fusion strain and subtracted from the corresponding n($^{me}$A)/n(reads) ratio from the (pSIR3)NLSEcoG2 strain (standalone EcoG2 with an NLS sequence under the control of the SIR3 promoter). The resulting Δ (n($^{me}$A)/n(reads)) ratio of each 400bp genome segment was then divided by the number of Adenines or Thymidines in the Watson strand, to obtain the $^{me}$A density per 400bp region per cell in the Watson or the Crick strand, respectively. The fraction of cells with at least 1 $^{me}$A per 400bp genome segment was determined from the Δ(n($^{me}$A)/n(reads)) ratio for each 400bp segment. All 400bp segments with a ratio $\geq 1$ were considered as being contacted by Sir3 in 100% of cells in the population. All the ratios with values <1 and >0 represent the fraction of cells in the population in which Sir3 has contacted that particular genome segment.

### References


1. T. H. Cheng, M. R. Gartenberg, Yeast heterochromatin is a dynamic structure that requires silencers continuously. *Genes Dev* **14**, 452-463 (2000).
2. Y. Katan-Khaykovich, K. Struhl, Heterochromatin formation involves changes in histone modifications over multiple cell generations. *Embo J* **24**, 2138-2149 (2005).
3. M. Sobecki *et al.*, MadID, a Versatile Approach to Map Protein-DNA Interactions, Highlights Telomere-Nuclear Envelope Contact Sites in Human Cells. *Cell Rep* **25**, 2891-2903.e2895 (2018).
4. A. Weiner *et al.*, High-resolution chromatin dynamics during a yeast stress response. *Mol Cell* **58**, 371-386 (2015).
