## Supplementa Table S2 for "The budding yeast heterochromatic protein Sir3 modulates genome-wide gene expression through transient direct contacts with euchromatin"

| cluster 1 Sir3 | Colonne1 | Colonne2 | Colonne3 | Colonne4 | Colonne5 | Colonne6 | cluster 1 | Colonne7 | Colonne8 | Colonne9 | Colonne1 | Colonne11 | Colonne12 |
|---|---|---|---|---|---|---|---|---|---|---|---|---|---|
| **cluster 1 Sir3D most similar to WT** | | | | | | | **cluster 1 Sir3D least similar to WT** | | | | | | |
| N | X | LOD | P | P_adj | attrib ID | attrib name | N | X | LOD | P | P_adj | attrib ID | attrib name |
| 15 | 68 | 0.66671 | 9.14E-06 | 0.008 | GO:0010927 | cellular component assembly involved in morphogenesis | 4 | 4 | 2.14209277 | 1.41E-05 | 0.018 | GO:0000798 | nuclear cohesin complex |
| 35 | 238 | 0.45906 | 6.24E-07 | <0.001 | GO:1903046 | meiotic cell cycle process | 4 | 4 | 2.14209277 | 1.41E-05 | 0.018 | GO:0008278 | cohesin complex |
| 43 | 337 | 0.38985 | 1.71E-06 | 0.002 | GO:0044702 | single organism reproductive process | 4 | 4 | 2.14209277 | 1.41E-05 | 0.018 | GO:0030892 | mitotic cohesin complex |
| 51 | 450 | 0.33286 | 6.28E-06 | 0.007 | GO:0022414 | reproductive process | 4 | 4 | 2.14209277 | 1.41E-05 | 0.018 | GO:0034990 | nuclear mitotic cohesin complex |
| | | | | | | | 5 | 7 | 1.53123477 | 1.62E-05 | 0.024 | GO:0036297 | interstrand cross-link repair |
| | | | | | | | 7 | 12 | 1.32551433 | 1.90E-06 | 0.005 | GO:0043596 | nuclear replication fork |
| | | | | | | | 10 | 22 | 1.11794238 | 2.28E-07 | 0.001 | GO:0005657 | replication fork |
| | | | | | | | 12 | 27 | 1.10227341 | 1.84E-08 | <0.001 | GO:0007064 | mitotic sister chromatid cohesion |
| | | | | | | | 13 | 42 | 0.85632866 | 7.27E-07 | 0.001 | GO:0007062 | sister chromatid cohesion |
| | | | | | | | 10 | 34 | 0.8248244 | 2.37E-05 | 0.032 | GO:0015631 | tubulin binding |
| | | | | | | | 16 | 71 | 0.67053476 | 4.29E-06 | 0.007 | GO:0071103 | DNA conformation change |
| | | | | | | | 15 | 72 | 0.62672619 | 2.35E-05 | 0.032 | GO:0000725 | recombinational repair |
| | | | | | | | 21 | 106 | 0.60137223 | 1.33E-06 | 0.003 | GO:0006302 | double-strand break repair |
| | | | | | | | 22 | 117 | 0.57350716 | 1.87E-06 | 0.005 | GO:0051052 | regulation of DNA metabolic process |
| | | | | | | | 49 | 283 | 0.54796537 | 1.48E-11 | <0.001 | GO:0006281 | DNA repair |
| | | | | | | | 42 | 253 | 0.51965458 | 1.83E-09 | <0.001 | GO:0051276 | chromosome organization |
| | | | | | | | 28 | 171 | 0.50281025 | 1.41E-06 | 0.003 | GO:0005856 | cytoskeleton |
| | | | | | | | 26 | 159 | 0.50097032 | 3.47E-06 | 0.007 | GO:0005694 | chromosome |
| | | | | | | | 49 | 329 | 0.46673538 | 3.39E-09 | <0.001 | GO:0006974 | cellular response to DNA damage stimulus |
| | | | | | | | 40 | 274 | 0.44974193 | 1.84E-07 | 0.001 | GO:1903047 | mitotic cell cycle process |
| | | | | | | | 34 | 253 | 0.40270182 | 1.05E-05 | 0.01 | GO:0044454 | nuclear chromosome part |
| | | | | | | | 71 | 556 | 0.4008289 | 8.49E-10 | <0.001 | GO:0022402 | cell cycle process |
| | | | | | | | 51 | 389 | 0.40043458 | 1.22E-07 | 0.001 | GO:0044427 | chromosomal part |
| | | | | | | | 63 | 489 | 0.39962215 | 6.37E-09 | <0.001 | GO:0006259 | DNA metabolic process |
| | | | | | | | 46 | 401 | 0.3272861 | 2.15E-05 | 0.03 | GO:0007049 | cell cycle |
| | | | | | | | 59 | 568 | 0.28253726 | 2.97E-05 | 0.035 | GO:0003677 | DNA binding |
| | | | | | | | 79 | 806 | 0.26276258 | 9.61E-06 | 0.01 | GO:1902589 | single-organism organelle organization |
| | | | | | | | 99 | 1063 | 0.24618927 | 5.19E-06 | 0.008 | GO:0006996 | organelle organization |
| **cluster 2 Sir3D most similar to WT** | | | | | | | cluster 2 Sir3D least similar to WT | | | | | | |
| N | X | LOD | P | P_adj | attrib ID | attrib name | N | X | LOD | P | P_adj | attrib ID | attrib name |
| 102 | 231 | 0.26675 | 5.56E-06 | 0.009 | GO:0044451 | nucleoplasm part | 119 | 693 | 0.2619964 | 1.16E-07 | <0.001 | GO:0006464 | cellular protein modification process |
| | | | | | | | 119 | 693 | 0.2619964 | 1.16E-07 | <0.001 | GO:0036211 | protein modification process |
| | | | | | | | 532 | 4129 | 0.25970467 | 7.60E-12 | <0.001 | GO:0043227 | membrane-bounded organelle |
| | | | | | | | 532 | 4129 | 0.25970467 | 7.60E-12 | <0.001 | GO:0043231 | intracellular membrane-bounded organelle |
| | | | | | | | 81 | 478 | 0.24313745 | 2.65E-05 | 0.025 | GO:1902494 | catalytic complex |
| | | | | | | | 144 | 918 | 0.2164965 | 1.36E-06 | <0.001 | GO:0031326 | regulation of cellular biosynthetic process |
| | | | | | | | 144 | 919 | 0.21584845 | 1.45E-06 | <0.001 | GO:0009889 | regulation of biosynthetic process |
| | | | | | | | 134 | 854 | 0.21377974 | 3.45E-06 | 0.002 | GO:0043412 | macromolecule modification |
| | | | | | | | 145 | 936 | 0.20934357 | 2.60E-06 | 0.002 | GO:0051171 | regulation of nitrogen compound metabolic process |
| | | | | | | | 137 | 886 | 0.20598633 | 5.90E-06 | 0.005 | GO:0010556 | regulation of macromolecule biosynthetic process |
| | | | | | | | 171 | 1126 | 0.20492265 | 9.17E-07 | <0.001 | GO:0080090 | regulation of primary metabolic process |
| | | | | | | | 177 | 1179 | 0.19960865 | 1.21E-06 | <0.001 | GO:0031090 | organelle membrane |
| | | | | | | | 172 | 1148 | 0.19690455 | 2.10E-06 | 0.001 | GO:0031323 | regulation of cellular metabolic process |
| | | | | | | | 129 | 849 | 0.19354415 | 2.88E-05 | 0.028 | GO:0010468 | regulation of gene expression |
| | | | | | | | 132 | 872 | 0.19212737 | 2.74E-05 | 0.027 | GO:2000112 | regulation of cellular macromolecule biosynthetic process |
| | | | | | | | 165 | 1108 | 0.19118772 | 5.39E-06 | 0.002 | GO:0060255 | regulation of macromolecule metabolic process |
| | | | | | | | 546 | 4505 | 0.1850397 | 1.96E-06 | 0.001 | GO:0043229 | intracellular organelle |
| | | | | | | | 546 | 4506 | 0.18468453 | 2.05E-06 | 0.001 | GO:0043226 | organelle |
| | | | | | | | 192 | 1328 | 0.18037001 | 5.28E-06 | 0.002 | GO:0019222 | regulation of metabolic process |
| | | | | | | | 230 | 1676 | 0.15689934 | 2.19E-05 | 0.024 | GO:0050789 | regulation of biological process |
| | | | | | | | 259 | 1910 | 0.15683129 | 1.19E-05 | 0.007 | GO:0065007 | biological regulation |
| | | | | | | | 212 | 1543 | 0.15293645 | 5.19E-05 | 0.047 | GO:0050794 | regulation of cellular process |
| **cluster 3 Sir3D most similar to WT** | | | | | | | cluster 3 Sir3D least similar to WT | | | | | | |
| N | X | LOD | P | P_adj | attrib ID | attrib name | N | X | LOD | P | P_adj | attrib ID | attrib name |
| 8 | 14 | 1.43252 | 5.23E-08 | <0.001 | GO:0005736 | DNA-directed RNA polymerase I complex | 8 | 18 | 1.11324985 | 3.88E-06 | 0.003 | GO:0030173 | integral component of Golgi membrane |
| 6 | 13 | 1.25087 | 1.36E-05 | 0.015 | GO:0005751 | mitochondrial respiratory chain complex IV | 8 | 18 | 1.11324985 | 3.88E-06 | 0.003 | GO:0031228 | intrinsic component of Golgi membrane |
| 6 | 13 | 1.25087 | 1.36E-05 | 0.015 | GO:0045277 | respiratory chain complex IV | 28 | 201 | 0.43238282 | 1.99E-05 | 0.028 | GO:0000329 | fungal-type vacuole membrane |
| 11 | 34 | 1.00887 | 2.31E-07 | <0.001 | GO:0003899 | DNA-directed RNA polymerase activity | 28 | 201 | 0.43238282 | 1.99E-05 | 0.028 | GO:0098852 | lytic vacuole membrane |
| 11 | 34 | 1.00887 | 2.31E-07 | <0.001 | GO:0034062 | RNA polymerase activity | 33 | 258 | 0.39100783 | 2.23E-05 | 0.028 | GO:0005774 | vacuolar membrane |
| 9 | 29 | 0.98246 | 4.44E-06 | 0.002 | GO:0006360 | transcription from RNA polymerase I promoter | 34 | 269 | 0.3854284 | 2.15E-05 | 0.028 | GO:0044437 | vacuolar part |
| 11 | 36 | 0.97325 | 4.46E-07 | <0.001 | GO:0055029 | nuclear DNA-directed RNA polymerase complex | 365 | 5755 | 0.35845845 | 5.06E-05 | 0.05 | GO:0044464 | cell part |
| 11 | 38 | 0.94032 | 8.21E-07 | <0.001 | GO:0000428 | DNA-directed RNA polymerase complex | 358 | 5576 | 0.33199011 | 2.12E-05 | 0.028 | GO:0044424 | intracellular part |
| 11 | 38 | 0.94032 | 8.21E-07 | <0.001 | GO:0030880 | RNA polymerase complex | 74 | 782 | 0.25717406 | 2.42E-05 | 0.028 | GO:0044711 | single-organism biosynthetic process |
| 8 | 29 | 0.91193 | 4.01E-05 | 0.039 | GO:0098803 | respiratory chain complex | 80 | 857 | 0.25327759 | 1.73E-05 | 0.023 | GO:0098588 | bounding membrane of organelle |
| 11 | 53 | 0.75019 | 2.75E-05 | 0.027 | GO:0098781 | ncRNA transcription | 289 | 4129 | 0.24096136 | 1.17E-06 | 0.001 | GO:0043227 | membrane-bounded organelle |
| 16 | 80 | 0.73172 | 6.98E-07 | <0.001 | GO:0098800 | inner mitochondrial membrane protein complex | 289 | 4129 | 0.24096136 | 1.17E-06 | 0.001 | GO:0043231 | intracellular membrane-bounded organelle |
| 12 | 61 | 0.72118 | 2.11E-05 | 0.016 | GO:0009144 | purine nucleoside triphosphate metabolic process | 249 | 3399 | 0.2286253 | 6.52E-07 | <0.001 | GO:0044444 | cytoplasmic part |
| 33 | 195 | 0.65738 | 7.76E-11 | <0.001 | GO:0042254 | ribosome biogenesis | 303 | 4505 | 0.21337202 | 3.77E-05 | 0.042 | GO:0043229 | intracellular organelle |
| 36 | 217 | 0.65006 | 1.81E-11 | <0.001 | GO:0022613 | ribonucleoprotein complex biogenesis | 303 | 4506 | 0.21303318 | 3.87E-05 | 0.043 | GO:0043226 | organelle |
| 14 | 82 | 0.64618 | 2.39E-05 | 0.023 | GO:0030687 | preribosome, large subunit precursor | 254 | 3609 | 0.19431442 | 2.36E-05 | 0.028 | GO:0044699 | single-organism process |
| 19 | 120 | 0.60926 | 2.72E-06 | 0.001 | GO:0098798 | mitochondrial protein complex | | | | | | | |
| 16 | 105 | 0.58766 | 2.77E-05 | 0.027 | GO:0006457 | protein folding | | | | | | | |
| 294 | 5755 | 0.57524 | 5.50E-07 | <0.001 | GO:0044464 | cell part | | | | | | | |
| 17 | 115 | 0.57247 | 2.34E-05 | 0.023 | GO:0035770 | ribonucleoprotein granule | | | | | | | |
| 17 | 115 | 0.57247 | 2.34E-05 | 0.023 | GO:0036464 | cytoplasmic ribonucleoprotein granule | | | | | | | |
| 38 | 277 | 0.5518 | 1.53E-09 | <0.001 | GO:0044085 | cellular component biogenesis | | | | | | | |
| 43 | 343 | 0.51002 | 2.03E-09 | <0.001 | GO:0005730 | nucleolus | | | | | | | |
| 21 | 170 | 0.48308 | 4.32E-05 | 0.04 | GO:0044455 | mitochondrial membrane part | | | | | | | |
| 288 | 5576 | 0.46843 | 1.01E-06 | <0.001 | GO:0044424 | intracellular part | | | | | | | |
| 38 | 330 | 0.46105 | 1.93E-07 | <0.001 | GO:0098796 | membrane protein complex | | | | | | | |
| 55 | 545 | 0.40911 | 2.86E-08 | <0.001 | GO:0034660 | ncRNA metabolic process | | | | | | | |
| 257 | 4505 | 0.38821 | 9.33E-10 | <0.001 | GO:0043229 | intracellular organelle | | | | | | | |
| 257 | 4506 | 0.38787 | 9.66E-10 | <0.001 | GO:0043226 | organelle | | | | | | | |
| 285 | 5572 | 0.38765 | 1.88E-05 | 0.016 | GO:0008150 | biological_process | | | | | | | |
| 241 | 4129 | 0.34663 | 2.34E-09 | <0.001 | GO:0043227 | membrane-bounded organelle | | | | | | | |
| 241 | 4129 | 0.34663 | 2.34E-09 | <0.001 | GO:0043231 | intracellular membrane-bounded organelle | | | | | | | |
| 154 | 2160 | 0.32648 | 1.89E-10 | <0.001 | GO:0032991 | macromolecular complex | | | | | | | |
| 83 | 1062 | 0.29862 | 7.06E-07 | <0.001 | GO:0043228 | non-membrane-bounded organelle | | | | | | | |
| 83 | 1062 | 0.29862 | 7.06E-07 | <0.001 | GO:0043232 | intracellular non-membrane-bounded organelle | | | | | | | |
| 112 | 1509 | 0.29817 | 4.60E-08 | <0.001 | GO:0043234 | protein complex | | | | | | | |
| 52 | 638 | 0.29467 | 4.64E-05 | 0.042 | GO:1902582 | single-organism intracellular transport | | | | | | | |
| 266 | 5087 | 0.28326 | 4.39E-05 | 0.04 | GO:0009987 | cellular process | | | | | | | |
| 137 | 2130 | 0.23327 | 5.09E-06 | 0.003 | GO:0005634 | nucleus | | | | | | | |
| 138 | 2169 | 0.22682 | 8.62E-06 | 0.007 | GO:0003824 | catalytic activity | | | | | | | |
| **cluster 4 Sir3D most similar to WT** | | | | | | | cluster 4 Sir3D least similar to WT | | | | | | |
| N | X | LOD | P | P_adj | attrib ID | attrib name | N | X | LOD | P | P_adj | attrib ID | attrib name |
| 8 | 17 | 1.64781 | 6.02E-10 | <0.001 | GO:0006094 | gluconeogenesis | 4 | 14 | 1.47640083 | 3.84E-05 | 0.05 | GO:0004298 | threonine-type endopeptidase activity |
| 8 | 17 | 1.64781 | 6.02E-10 | <0.001 | GO:0019319 | hexose biosynthetic process | 4 | 14 | 1.47640083 | 3.84E-05 | 0.05 | GO:0010499 | proteasomal ubiquitin-independent protein catabolic process |
| 8 | 18 | 1.60427 | 1.06E-09 | <0.001 | GO:0046364 | monosaccharide biosynthetic process | 4 | 14 | 1.47640083 | 3.84E-05 | 0.05 | GO:0070003 | threonine-type peptidase activity |
| 7 | 18 | 1.50692 | 3.85E-08 | <0.001 | GO:0006407 | rRNA export from nucleus | 10 | 89 | 0.99022749 | 5.41E-07 | 0.001 | GO:0010494 | cytoplasmic stress granule |
| 7 | 18 | 1.50692 | 3.85E-08 | <0.001 | GO:0051029 | rRNA transport | 10 | 115 | 0.86553332 | 5.80E-06 | 0.007 | GO:0035770 | ribonucleoprotein granule |
| 9 | 24 | 1.48653 | 5.71E-10 | <0.001 | GO:0006096 | glycolytic process | 10 | 115 | 0.86553332 | 5.80E-06 | 0.007 | GO:0036464 | cytoplasmic ribonucleoprotein granule |
| 9 | 24 | 1.48653 | 5.71E-10 | <0.001 | GO:0006757 | ATP generation from ADP | 14 | 251 | 0.66506032 | 1.42E-05 | 0.024 | GO:0006520 | cellular amino acid metabolic process |
| 5 | 14 | 1.44857 | 6.30E-06 | 0.011 | GO:0000028 | ribosomal small subunit assembly | 19 | 397 | 0.60901292 | 3.35E-06 | 0.007 | GO:0019752 | carboxylic acid metabolic process |
| 9 | 26 | 1.43369 | 1.32E-09 | <0.001 | GO:0006165 | nucleoside diphosphate phosphorylation | 19 | 414 | 0.58869106 | 6.21E-06 | 0.007 | GO:0043436 | oxoacid metabolic process |
| 10 | 29 | 1.4335 | 1.59E-10 | <0.001 | GO:0046031 | ADP metabolic process | 19 | 415 | 0.58752123 | 6.43E-06 | 0.007 | GO:0006082 | organic acid metabolic process |
| 10 | 30 | 1.41171 | 2.34E-10 | <0.001 | GO:0009135 | purine nucleoside diphosphate metabolic process | 25 | 736 | 0.46238717 | 3.54E-05 | 0.046 | GO:0044281 | small molecule metabolic process |
| 10 | 30 | 1.41171 | 2.34E-10 | <0.001 | GO:0009179 | purine ribonucleoside diphosphate metabolic process | 30 | 926 | 0.45766896 | 1.12E-05 | 0.012 | GO:1901564 | organonitrogen compound metabolic process |
| 10 | 30 | 1.41171 | 2.34E-10 | <0.001 | GO:0009185 | ribonucleoside diphosphate metabolic process | 57 | 2274 | 0.45440656 | 4.45E-07 | 0.001 | GO:0005737 | cytoplasm |
| 44 | 165 | 1.39753 | 2.03E-38 | <0.001 | GO:0002181 | cytoplasmic translation | | | | | | | |
| 19 | 64 | 1.36515 | 1.25E-17 | <0.001 | GO:0022627 | cytosolic small ribosomal subunit | | | | | | | |
| 10 | 33 | 1.35218 | 6.83E-10 | <0.001 | GO:0009132 | nucleoside diphosphate metabolic process | | | | | | | |
| 10 | 33 | 1.35218 | 6.83E-10 | <0.001 | GO:0046939 | nucleotide phosphorylation | | | | | | | |
| 25 | 91 | 1.33855 | 4.91E-22 | <0.001 | GO:0022625 | cytosolic large ribosomal subunit | | | | | | | |

| | | | | | | |
|---|---|---|---|---|---|---|
| 48 | 222 | 1.29358 | 2.19E-37 | <0.001 | GO:0044445 | cytosolic part |
| 9 | 37 | 1.22112 | 4.32E-08 | <0.001 | GO:0006090 | pyruvate metabolic process |
| 44 | 232 | 1.20205 | 1.80E-31 | <0.001 | GO:0003735 | structural constituent of ribosome |
| 46 | 247 | 1.20096 | 1.29E-32 | <0.001 | GO:0005840 | ribosome |
| 44 | 238 | 1.18801 | 5.73E-31 | <0.001 | GO:0044391 | ribosomal subunit |
| 8 | 36 | 1.16937 | 5.37E-07 | <0.001 | GO:0006006 | glucose metabolic process |
| 37 | 203 | 1.15082 | 1.26E-25 | <0.001 | GO:0005622 | intracellular |
| 19 | 99 | 1.11492 | 8.40E-14 | <0.001 | GO:0015935 | small ribosomal subunit |
| 25 | 139 | 1.0992 | 3.35E-17 | <0.001 | GO:0015934 | large ribosomal subunit |
| 7 | 36 | 1.09655 | 7.39E-06 | 0.011 | GO:0000027 | ribosomal large subunit assembly |
| 52 | 369 | 1.07834 | 7.04E-31 | <0.001 | GO:0006412 | translation |
| 52 | 373 | 1.07261 | 1.22E-30 | <0.001 | GO:0043043 | peptide biosynthetic process |
| 10 | 56 | 1.0542 | 1.75E-07 | <0.001 | GO:0046034 | ATP metabolic process |
| 52 | 399 | 1.03694 | 3.70E-29 | <0.001 | GO:0006518 | peptide metabolic process |
| 10 | 59 | 1.02684 | 2.93E-07 | <0.001 | GO:0009205 | purine ribonucleoside triphosphate metabolic process |
| 48 | 365 | 1.02336 | 6.72E-27 | <0.001 | GO:0005198 | structural molecule activity |
| 52 | 415 | 1.01622 | 2.65E-28 | <0.001 | GO:0043604 | amide biosynthetic process |
| 10 | 61 | 1.0095 | 4.05E-07 | <0.001 | GO:0009144 | purine nucleoside triphosphate metabolic process |
| 11 | 69 | 0.99668 | 1.40E-07 | <0.001 | GO:0016051 | carbohydrate biosynthetic process |
| 12 | 76 | 0.9936 | 4.10E-08 | <0.001 | GO:0019362 | pyridine nucleotide metabolic process |
| 10 | 63 | 0.99282 | 5.55E-07 | <0.001 | GO:0009199 | ribonucleoside triphosphate metabolic process |
| 8 | 52 | 0.97474 | 1.01E-05 | 0.016 | GO:0019318 | hexose metabolic process |
| 12 | 79 | 0.97364 | 6.42E-08 | <0.001 | GO:0009126 | purine nucleoside monophosphate metabolic process |
| 12 | 79 | 0.97364 | 6.42E-08 | <0.001 | GO:0009167 | purine ribonucleoside monophosphate metabolic process |
| 12 | 79 | 0.97364 | 6.42E-08 | <0.001 | GO:0030686 | 90S preribosome |
| 53 | 464 | 0.97231 | 7.07E-27 | <0.001 | GO:0043603 | cellular amide metabolic process |
| 11 | 75 | 0.95385 | 3.38E-07 | <0.001 | GO:0046496 | nicotinamide nucleotide metabolic process |
| 10 | 71 | 0.93174 | 1.74E-06 | 0.002 | GO:0009141 | nucleoside triphosphate metabolic process |
| 12 | 88 | 0.91866 | 2.19E-07 | <0.001 | GO:0072524 | pyridine-containing compound metabolic process |
| 13 | 97 | 0.91189 | 8.24E-08 | <0.001 | GO:0009161 | ribonucleoside monophosphate metabolic process |
| 8 | 59 | 0.91081 | 2.63E-05 | 0.029 | GO:0005996 | monosaccharide metabolic process |
| 12 | 90 | 0.90731 | 2.82E-07 | <0.001 | GO:0006733 | oxidoreduction coenzyme metabolic process |
| 12 | 91 | 0.90174 | 3.20E-07 | <0.001 | GO:0042278 | purine nucleoside metabolic process |
| 12 | 91 | 0.90174 | 3.20E-07 | <0.001 | GO:0046128 | purine ribonucleoside metabolic process |
| 9 | 68 | 0.89929 | 9.90E-06 | 0.016 | GO:0044724 | single-organism carbohydrate catabolic process |
| 13 | 100 | 0.89653 | 1.19E-07 | <0.001 | GO:0009123 | nucleoside monophosphate metabolic process |
| 12 | 93 | 0.89081 | 4.07E-07 | <0.001 | GO:0009150 | purine ribonucleotide metabolic process |
| 10 | 78 | 0.88443 | 4.19E-06 | 0.008 | GO:0016052 | carbohydrate catabolic process |
| 73 | 926 | 0.88177 | 4.57E-28 | <0.001 | GO:1901564 | organonitrogen compound metabolic process |
| 21 | 175 | 0.87658 | 5.07E-11 | <0.001 | GO:0030684 | preribosome |
| 12 | 97 | 0.86972 | 6.49E-07 | <0.001 | GO:0006163 | purine nucleotide metabolic process |
| 13 | 107 | 0.86261 | 2.68E-07 | <0.001 | GO:0009119 | ribonucleoside metabolic process |
| 62 | 733 | 0.85835 | 1.24E-24 | <0.001 | GO:1901566 | organonitrogen compound biosynthetic process |
| 13 | 108 | 0.85797 | 2.99E-07 | <0.001 | GO:0009259 | ribonucleotide metabolic process |
| 14 | 128 | 0.81246 | 3.42E-07 | <0.001 | GO:0019693 | ribose phosphate metabolic process |
| 13 | 119 | 0.80985 | 9.34E-07 | 0.001 | GO:0009116 | nucleoside metabolic process |
| 57 | 723 | 0.7968 | 8.48E-21 | <0.001 | GO:0030529 | intracellular ribonucleoprotein complex |
| 57 | 723 | 0.7968 | 8.48E-21 | <0.001 | GO:1990904 | ribonucleoprotein complex |
| 13 | 124 | 0.78958 | 1.50E-06 | 0.002 | GO:1901657 | glycosyl compound metabolic process |
| 12 | 116 | 0.78124 | 4.47E-06 | 0.008 | GO:0072521 | purine-containing compound metabolic process |
| 14 | 150 | 0.73458 | 2.38E-06 | 0.003 | GO:0022618 | ribonucleoprotein complex assembly |
| 14 | 160 | 0.70317 | 5.13E-06 | 0.008 | GO:0071826 | ribonucleoprotein complex subunit organization |
| 14 | 165 | 0.68824 | 7.35E-06 | 0.011 | GO:0006913 | nucleocytoplasmic transport |
| 14 | 166 | 0.68531 | 7.89E-06 | 0.011 | GO:0051169 | nuclear transport |
| 16 | 195 | 0.67599 | 2.44E-06 | 0.004 | GO:0009117 | nucleotide metabolic process |
| 16 | 199 | 0.66614 | 3.18E-06 | 0.008 | GO:0006753 | nucleoside phosphate metabolic process |
| 94 | 2274 | 0.64353 | 1.39E-16 | <0.001 | GO:0005737 | cytoplasm |
| 66 | 1228 | 0.6319 | 1.67E-15 | <0.001 | GO:0044271 | cellular nitrogen compound biosynthetic process |
| 59 | 1062 | 0.61847 | 3.43E-14 | <0.001 | GO:0043228 | non-membrane-bounded organelle |
| 59 | 1062 | 0.61847 | 3.43E-14 | <0.001 | GO:0043232 | intracellular non-membrane-bounded organelle |
| 16 | 220 | 0.61758 | 1.16E-05 | 0.016 | GO:0055086 | nucleobase-containing small molecule metabolic process |
| 58 | 1085 | 0.59315 | 3.69E-13 | <0.001 | GO:0044267 | cellular protein metabolic process |
| 88 | 2216 | 0.57098 | 1.06E-13 | <0.001 | GO:0034641 | cellular nitrogen compound metabolic process |
| 122 | 4685 | 0.56985 | 1.26E-07 | <0.001 | GO:0003674 | molecular_function |
| 82 | 2010 | 0.55101 | 5.52E-13 | <0.001 | GO:0009058 | biosynthetic process |
| 81 | 1979 | 0.5473 | 7.79E-13 | <0.001 | GO:1901576 | organic substance biosynthetic process |
| 90 | 2394 | 0.54652 | 1.29E-12 | <0.001 | GO:0006807 | nitrogen compound metabolic process |
| 62 | 1319 | 0.53808 | 1.24E-11 | <0.001 | GO:0019538 | protein metabolic process |
| 124 | 5087 | 0.50188 | 1.77E-05 | 0.02 | GO:0009987 | cellular process |
| 61 | 1400 | 0.49056 | 5.60E-10 | <0.001 | GO:0034645 | cellular macromolecule biosynthetic process |
| 61 | 1424 | 0.48074 | 1.15E-09 | <0.001 | GO:0009059 | macromolecule biosynthetic process |
| 99 | 3257 | 0.44428 | 2.55E-08 | <0.001 | GO:0044238 | primary metabolic process |
| 72 | 1927 | 0.44314 | 6.05E-09 | <0.001 | GO:0044249 | cellular biosynthetic process |
| 77 | 2160 | 0.43507 | 9.30E-09 | <0.001 | GO:0032991 | macromolecular complex |
| 103 | 3623 | 0.4129 | 4.82E-07 | <0.001 | GO:0044237 | cellular metabolic process |
| 107 | 3918 | 0.40569 | 1.83E-06 | 0.003 | GO:0008152 | metabolic process |
| 99 | 3399 | 0.40505 | 3.89E-07 | <0.001 | GO:0044444 | cytoplasmic part |
| 102 | 3675 | 0.38024 | 3.06E-06 | 0.006 | GO:0071704 | organic substance metabolic process |
